# Transposon expansion is associated with reorganization of small RNA and DNA methylation landscapes in the morphologically minimal angiosperm *Wolffia brasiliensis*

**DOI:** 10.64898/2026.02.27.708460

**Authors:** Daniel Buendía-Ávila, Verónica Barragán-Borrero, Pablo Luna-Rodriguez, Mary Akinyuwa, Laura Morello, Arturo Marí-Ordóñez

## Abstract

Genome expansion in angiosperms is largely driven by transposable element (TE) proliferation, counteracted by epigenetic silencing. To investigate how TE amplification reshapes silencing landscapes, we compared two closely related, clonally propagating duckweeds, the TE-rich *Wolffia brasiliensis*, for which we report a draft genome assembly, and the TE-poor *Spirodela polyrhiza*, that share a broadly conserved silencing and methylation machinery. *W. brasiliensis* displays extensive and recent TE amplification with pervasive TE–gene interspersion, elevated genome-wide CG methylation, and high levels of both 22- and 24-nt siRNAs. Systematic cross-species comparison reveals that per-element silencing rules are conserved between the two duckweeds: TE length predicting siRNA-producing capacity, inverted-repeat-forming TEs as productive PTGS-associated loci, 24-nt siRNA production predicting non-CG methylation, and intragenic constraint on RdDM-associated methylation. What differs is the genome-wide deployment of these rules across dramatically different TE loads and genome architectures: some divergent outcomes, including elevated 24-nt siRNA abundance and pervasive TE-wide CG methylation, scale directly with TE content, whereas others: near-exclusive 22-nt processing of PTGS substrates, gene-proximity-dependent RdDM at intergenic TEs, and a strong correlation between gene body CG methylation and intragenic TE content that is not detectable in *S. polyrhiza*; require additional *W. brasiliensis*-specific contributions. These findings position TEs as central determinants of the epigenetic architecture that emerges from plant genome expansion, and reveal previously underappreciated plasticity in conserved silencing pathways.

**Significance statement:** Comparing closely related duckweeds with contrasting TE loads but conserved silencing machinery, we show that per-element silencing rules are shared between the two species while the genome-wide outcomes those rules produce diverge profoundly with TE amplification and genome architecture. This reframes TE proliferation as a determinant not only of genome size but of the epigenetic architecture that emerges from it, uncovers a *Wolffia*-specific coupling between intragenic TE content and gene body CG methylation, and reveals unexpected plasticity in small RNA biogenesis pathways.

## Introduction

Transposable elements (TEs) are pervasive components of plant genomes and major contributors to genome size variation. Their activity profoundly influences chromosomal organization and regulatory landscapes across species (Lisch, 2013; Bennetzen and Wang, 2014). While TE mobilization can generate genetic novelty, uncontrolled transposition poses substantial risks, including insertional mutagenesis and genome instability (Quadrana and Henderson, 2025; Roulin, 2026; Tossolini *et al*., 2025; Wen *et al*., 2026). To mitigate these threats, plants have evolved multilayered epigenetic silencing strategies (Slotkin and Martienssen, 2007; Fedoroff, 2012; Sigman and Slotkin, 2016; Vaucheret and Voinnet, 2023). RNA-mediated silencing and epigenetic repression constitute central layers of genome defense, integrating small RNA pathways, DNA methylation, and histone modifications, which cooperate to repress transposable elements (TEs), regulate gene expression, and maintain genome stability (Slotkin and Martienssen, 2007; Lisch, 2009; Law and Jacobsen, 2010; Matzke and Mosher, 2014).

In angiosperms, TEs are repressed both transcriptionally, through DNA methylation and repressive histone modifications (transcriptional gene silencing, TGS), and post-transcriptionally, through degradation of their transcripts (post-transcriptional gene silencing, PTGS) (Matzke and Mosher, 2014; Vaucheret and Voinnet, 2023). CG methylation is maintained by METHYLTRANSFERASE 1 (MET1), ensuring stable inheritance across cell divisions, whereas non-CG methylation is maintained by a self-reinforcing loop with histone H3 lysine-9 dimethylation (H3K9me2): SUPPRESSOR OF VARIEGATION 3-9 HOMOLOG (SUVH) methyltransferases read DNA methylation to deposit H3K9me2, which the CHROMOMETHYLASEs CMT3 and CMT2 in turn recognize to reinforce CHG and CHH methylation, respectively (Erdmann and Picard, 2020). *De novo* methylation is established by RNA-directed DNA methylation (RdDM): RNA Polymerase IV (Pol IV) - recruited to target loci by CLASSY (CLSY) chromatin remodelers through the H3K9me2 reader SHH1, or directed to specific sequence motifs by DNA-binding transcription factors - transcribes precursors that RNA-DEPENDENT RNA POLYMERASE 2 (RDR2) and DICER-LIKE 3 (DCL3) process into 24-nt siRNAs; these load into the AGO4/6 ARGONAUTE clade and, through base-pairing with RNA Polymerase V (Pol V) transcripts, recruit the methyltransferase DRM2 to methylate cytosines in all sequence contexts (Erdmann and Picard, 2020; Wang *et al*., 2022; Wu *et al*., 2025; Xu *et al*., 2025; Xie *et al*., 2025). PTGS, by contrast, mainly relies on 21- and 22-nt small RNAs: micro RNAs (miRNAs) processed from imperfect hairpin precursors by DICER-LIKE 1 (DCL1) and siRNAs generated from double-stranded (ds)RNA mainly by DCL4 (21-nt) and DCL2 (22-nt). These small RNAs are loaded into members of the AGO1/5/10 and AGO2/7 clades to guide cleavage of complementary transcripts and are amplified into secondary siRNAs by RNA-DEPENDENT RNA POLYMERASE 6 (RDR6), providing an antiviral-like first line of defense against transcriptionally active TEs that can as well initiate RdDM (Slotkin *et al*., 2009; Marí-Ordóñez *et al*., 2013; Creasey *et al*., 2014; Fang and Qi, 2016; Zhan and Meyers, 2023).

Whereas PTGS acts mainly in active or epigenetically reactivated TEs, most TEs are kept silenced transcriptionally through small RNA-dependent (RdDM) and independent (CMT-H3K9me2) pathways. However, the relative contribution of these silencing pathways is not uniform across the genome. Extent and mode of TE repression depend on intrinsic properties of individual elements - such as their age, length, and transcriptional activity - as well as their genomic context, including proximity to genes and local chromatin environment. TE regulation is therefore dynamically shaped according to the structural and positional features of each TE (Zemach *et al*., 2013; Gent *et al*., 2013; Stroud *et al*., 2014; Sigman and Slotkin, 2016; Wang and Baulcombe, 2020; Quadrana and Henderson, 2025).

Diversity in epigenetic landscapes is also remarkable within clades of flowering plants. Differences in DNA methylation levels, small RNA populations, and chromatin organization have been associated with variation in evolutionary history, TE content and distribution, genome architecture, reproductive strategy, and composition of silencing pathways (Seymour *et al*., 2014; Bennetzen and Wang, 2014; Alonso *et al*., 2015; Niederhuth *et al*., 2016; Baduel and Sasaki, 2023; Baduel *et al*., 2024). However, because these variables often change simultaneously, it remains difficult to disentangle the specific contribution of TE load from broader divergence in genome structure and regulatory machinery (Seymour *et al*., 2014; Niederhuth *et al*., 2016; Bewick *et al*., 2017; Baduel and Sasaki, 2023). Especially variation in TE load represents a particularly compelling yet difficult-to-isolate parameter, as differences in TE abundance and configuration may influence not only the number of silencing targets but also the relative engagement of transcriptional and post-transcriptional pathways (Chen *et al*., 2011; Matzke and Mosher, 2014; Bennetzen and Wang, 2014; Wang *et al*., 2020; Baldrich *et al*., 2022; Arce *et al*., 2023; Martin *et al*., 2023). Whether quantitative differences in TE abundance alone are sufficient to reorganize epigenetic and small RNA regulatory regimes remains unresolved.

Comparative analyses within species or among closely related taxa have provided a powerful framework for investigating how TE landscapes and epigenetic regulation co-evolve, although limited to a small number of angiosperm families (Zemach *et al*., 2010; Seymour *et al*., 2014; Bewick *et al*., 2016; Eichten *et al*., 2016; Niederhuth *et al*., 2016; Stritt *et al*., 2017; Quadrana *et al*., 2019; Wyler *et al*., 2020; Igolkina *et al*., 2025). Disentangling the effects of TE proliferation from lineage-specific changes in silencing pathway composition requires a system in which phylogenetic distance and chromosomal organization are tightly controlled and allow a direct comparison between species that share a conserved regulatory background yet differ markedly in TE content.

Duckweeds (family *Lemnaceae*), free-floating flowering plants characterized by extreme morphological reduction and predominantly clonal reproduction (Landolt, 1986; Acosta *et al*., 2021), offer such a system. Closely related duckweed species exhibit striking variation in genome size and TE abundance (Hoang *et al*., 2019; Acosta *et al*., 2021; Tippery *et al*., 2021; Hoang *et al*., 2022; Ernst *et al*., 2025). Despite frequent polyploidization and interspecific hybridization (Braglia *et al*., 2021; Hoang *et al*., 2022; Michael *et al*., 2025; Lee *et al*., 2025), their genomes retain relatively conserved chromosome numbers and high levels of synteny, indicating that genome expansion has largely occurred through differential TE accumulation rather than extensive chromosomal rearrangement (Hoeck *et al*., 2015; Hoang *et al*., 2019; Hoang *et al*., 2020; Michael *et al*., 2020; Abramson *et al*., 2021; Hoang *et al*., 2022; Ernst *et al*., 2025). Additionally, due to their extensive clonal propagation, duckweeds provide a unique opportunity to evaluate how asexual reproduction influences small RNA- and epigenetic-based TE regulation in plants.

Analysis of the compact genome of *Spirodela polyrhiza* revealed unconventional TE regulation characterized by globally reduced DNA methylation across both genes and TEs, with RdDM activity largely restricted to a small subset of young, intact TEs, resulting in extremely low abundance of 24-nt siRNAs (An *et al*., 2019; Harkess *et al*., 2024; Dombey *et al*., 2025). This has been proposed to reflect developmental and reproductive features of duckweeds (Dombey *et al*., 2025). However, recent analyses across duckweed species indicate that those with higher TE content display higher global DNA methylation levels and greater accumulation of TE-derived 24-nt siRNAs (Ernst *et al*., 2025). This occurs despite lineage-wide losses of silencing components otherwise conserved across angiosperms. On the PTGS side, duckweeds lack the 22-nt–generating DCL2 among several other antiviral factors. On the TGS side, they have lost the Pol IV–recruiting chromatin factors CLSY1/2 and the H3K9me2 reader SHH1 - retaining only CLSY3 - together with the RdDM-clade AGO6. CHH methylation maintenance is also compromised by absence of CMT2 (Dombey *et al*., 2025; Ernst *et al*., 2025). Together, these observations indicate that in TE-rich duckweeds increased TE DNA methylation and 24-nt siRNAs can still be achieved with a reduced silencing toolkit, raising the possibility that differences in TE abundance and distribution may influence the extent and mode of silencing pathway deployment. Hence, duckweeds offer a suitable material to study this connection.

## Results

### 24-nt siRNAs are lineage-specifically enriched across duckweeds

To address whether TE load influences small RNA deployment across duckweeds, we profiled small RNAs across representative duckweed species spanning all five duckweed genera. Because reference genomes are unavailable for most accessions (clones), preventing locus-specific mapping of small RNAs to TEs, we quantified the relative abundance of 24-nt small interfering RNAs (siRNAs) in TraPR libraries representing ARGONAUTE (AGO)-loaded RNAs (Grentzinger *et al*., 2020) as a proxy for RdDM activity, including additional species and accessions not investigated in earlier studies (Dombey *et al*., 2025; Ernst *et al*., 2025).

Small RNA size distributions differed markedly among genera. Species belonging to *Spirodela*, *Landoltia*, or *Lemna* displayed relatively low proportions of 24-nt siRNAs, whereas species from *Wolffiella* and *Wolffia* exhibited higher relative abundance of 24-nt siRNAs, in some cases accompanied by a concomitant enrichment of 22-nt small RNAs (**Figure 1a**). Although the focus on AGO-associated small RNAs and differences in AGO composition or loading efficiency across species could modify the proportional representation, these data validate and extend previous observations (Ernst *et al*., 2025), indicating that enrichment of RdDM-associated small RNA signatures correlates with specific duckweed lineages. This provides a system in which the genomic and epigenetic basis of this lineage-specific increase of 24-nt siRNA relative to the 24-nt-poor *S. polyrhiza* can be investigated in detail.

**Figure 1.**
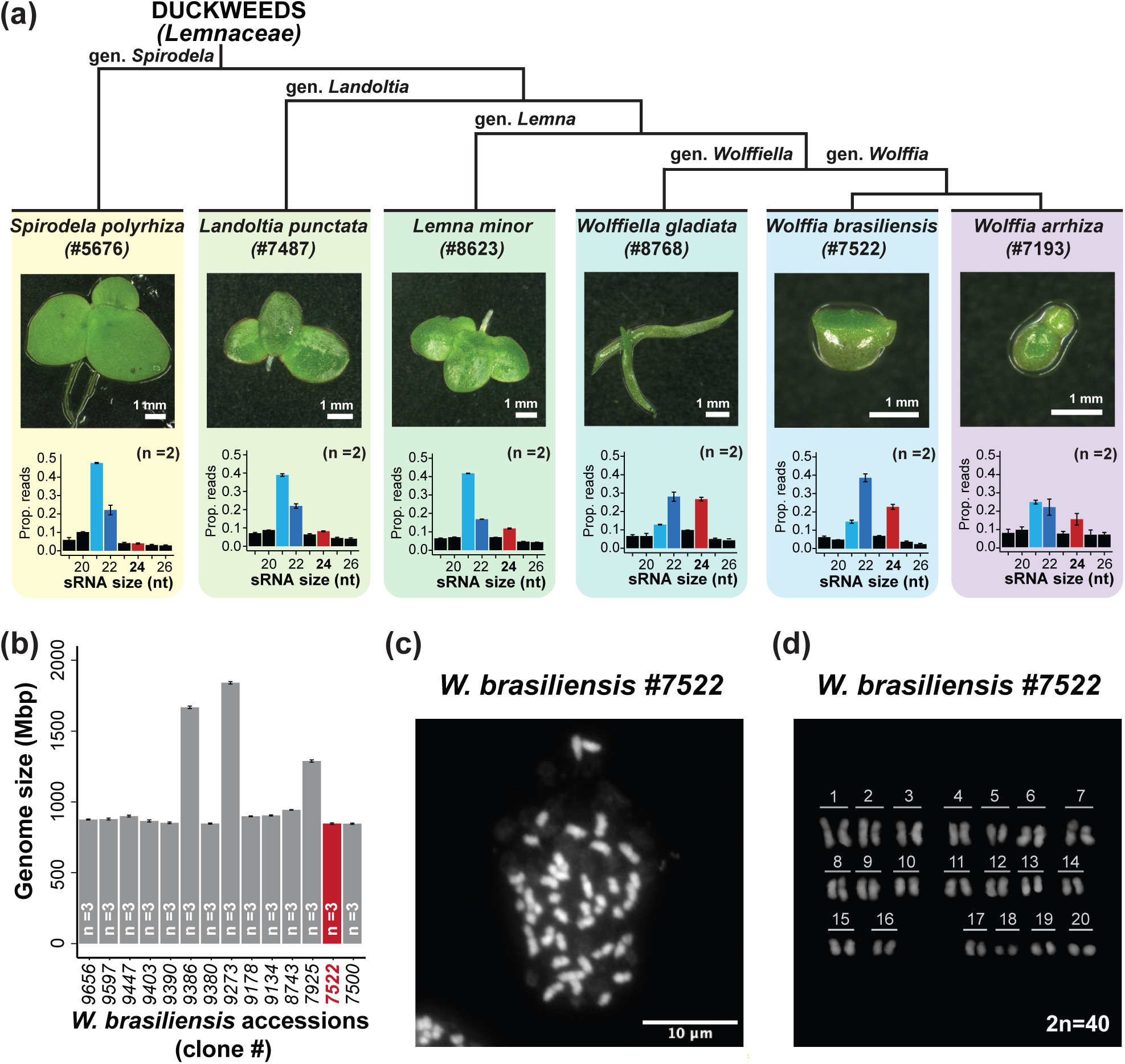
Identification of a *W. brasiliensis* accession with increased proportion of 24-nt siRNAs. **(a)** Phylogeny of selected duckweed species with representative images and corresponding small RNA size distribution profiles. Bars and error bars indicate the mean and the SD respectively (n=2). **(b)** Genome size estimation of *Wolffia brasiliensis* accessions by flow cytometry. Bars and error bars indicate the mean and the SD respectively (n=3). (**c)** DAPI-stained mitotic metaphase and (**d)** Representative karyogram of *W. brasiliensis* #7522.

### A diploid *Wolffia brasiliensis* accession with elevated 24-nt siRNAs is suitable for genome-wide analyses

Given the enrichment of 24-nt siRNAs observed in *Wolffia* species and the difficulty of maintaining *Wolffiella* cultures under our growth conditions, we focused subsequent analyses on *Wolffia brasiliensis* and *Wolffia arrhiza*. To enable genome-wide analyses while minimizing confounding effects of polyploidy or interspecific hybridization, we screened multiple accessions of both species by flow cytometry to select for diploid accessions.

Genome size measurements revealed substantial intraspecific variation in both species (**Figure 1b; Figure S1**), consistent with previous reports (Hoang *et al*., 2022). *W. brasiliensis* accessions displayed genome sizes clustered around 900 Mbp with comparatively limited variation, whereas *W. arrhiza* accessions segregated into two groups around ∼1500 and ∼2100 Mbp. Based on the more uniform genome size, we prioritized *W. brasiliensis* for detailed analysis. We selected *W. brasiliensis* accession #7522, which was included in the small RNA survey and exhibited a representative genome size for the species (Hoang *et al*., 2022) (**Figure 1b**). Cytological analysis of metaphase chromosome spreads confirmed a diploid chromosome number of 2n = 40 (**Figure 1c,d**), consistent with the predominant karyotype reported for *W. brasiliensis* and related duckweeds (Hoang *et al*., 2022).

Species of the genus *Wolffia* are among the smallest (∼ 1 mm) and morphologically simplest in angiosperms, lacking roots and vasculature, exemplifying the extreme morphological reduction and clonal growth characteristic of duckweeds (Landolt, 1986; Lemon and Posluszny, 2000; Acosta *et al*., 2021) (**Figure S2a**). Morphological inspection of accession #7522 revealed the characteristic dorsal papilla associated with *W. brasiliensis* (Landolt, 1986; Bog *et al*., 2020) (**Figure S2b**). Because morphological traits alone are often insufficient for precise taxonomic assignment in duckweeds, we further validated the identity of accession #7522 using Tubulin-Based Polymorphism (TBP) analysis (Bog *et al*., 2020; Braglia *et al*., 2021), confirming its classification as *W. brasiliensis* (**Figure S2c**). Together, these analyses establish *W. brasiliensis* #7522 as a diploid, taxonomically validated accession with elevated 24-nt siRNAs, providing a robust system for subsequent genome-wide investigation of TE regulation. Despite identical chromosome numbers, *W. brasiliensis* and *S. polyrhiza* differ dramatically in genome size (∼847 versus ∼138 Mbp) (Dombey *et al*., 2025), suggesting TE-driven expansion and motivating genome sequencing.

### Genome sequencing and annotation of *W. brasiliensis* #7522 allows characterization of TE content, structure and distribution

To determine whether the pronounced genome size expansion relative to *S. polyrhiza* is due to TE proliferation and to characterize the TE landscape underlying small RNA production in *W. brasiliensis*, we generated a long-read draft genome assembly for accession #7522 using PacBio HiFi sequencing (Wenger *et al*., 2019). Approximately 40× genome coverage yielded an assembly comprising 250 contigs with a total length of 771 Mbp, representing ∼90% of the genome size estimated by flow cytometry (**Figure S3a**). The assembly showed high continuity (N50 = 9.23 Mbp; L90 = 88) and completeness, with 91.4% of conserved plant genes recovered based on BUSCO analysis (Manni *et al*., 2021) (**Figure S3a**).

To support structural analysis and enable gene annotation, we complemented the genome assembly with transcriptomic data generated by short-read Illumina RNA sequencing and long-read PacBio Iso-Seq full-length transcript sequencing (Mortazavi *et al*., 2008; Sharon *et al*., 2013). Integration of these datasets resulted in the annotation of 25,161 gene models, of which 70.6% were supported by transcript evidence from one or both platforms (**Figure 2a**; **Figure S3b**). To further assess assembly completeness, we assembled a de novo transcriptome from RNA-seq reads that failed to map to the genome (Grabherr *et al*., 2011); 98.4% of the resulting transcripts remapped to our assembly, and only 39 of the residual transcripts had plant BLAST hits, only raising the BUSCO score from 91.4% to 92.1% (**Figure S3c, Table S1**) indicating that most of the transcribed genes were included in the assembly. This high-quality assembly and annotation provided a foundation for comprehensive characterization of TE content, structure, and genomic distribution.

**Figure 2.**
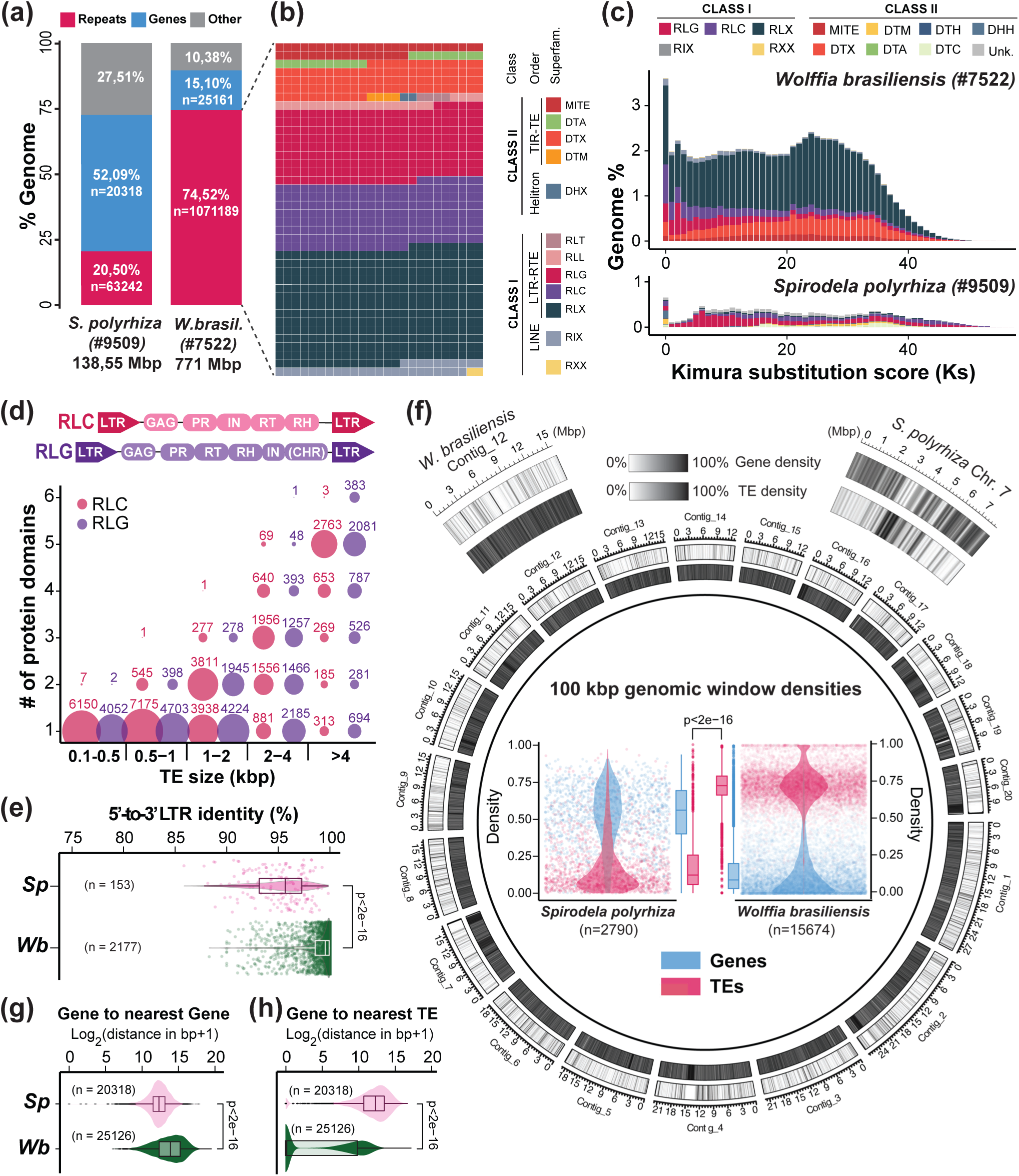
Genome composition and TE landscape of *W. brasiliensis*. **(a)** Genome size and occupancy of genes/TEs in *S. polyrhiza* and *W. brasiliensis*. **(b)** Waffle plot of different TE superfamilies in *W.* brasiliensis labelled by their three-letter classification codes (defined in Figure S3d). Each square represents 0.1% of the total repeat content. **(c)** Kimura plot of TE divergence and genome occupancy in *W. brasiliensis* and *S. polyrhiza*. **(d)** Bubble plot of protein domains identified per size category for RLC and RLG with at least one domain. **(e)** LTR identity (%) of intact TEs identified by EDTA for *S. polyrhiza* and *W. brasiliensis.* **(f)** Circos plot of gene (outer) and TE (inner) density across the 20 longest *W. brasiliensis* contigs, using 100 kb genomic windows. *S. polyrhiza* chromosome 7 is shown for comparison. Center: gene/TE density plots of all 100 kb genomic widows in *W. brasiliensis* and *S. polyrhiza*. **(g-h)** Distance between neighboring genes and genes to nearest TEs in *S. polyrhiza* (*Sp*) and *W. brasiliensis* (*Wb*). Unless otherwise stated, all boxplots in this study depict the median as a solid bar, with box upper and bottom limits representing the first and third quartiles. Whisker range is 1.5 times the interquartile range. P-values were calculated using Wilcoxon rank sum test.

### TE proliferation in the *W. brasiliensis* genome is extensive and recent

To assess the contribution of transposable elements to genome expansion, we performed *de novo* repeat annotation of the *W. brasiliensis* assembly. This revealed that TEs comprise 74.52% of the assembled *W. brasiliensis* genome (**Figure 2a**, **Figure S3d,e**). Because highly repetitive regions are often under-represented in *de novo* assemblies (Alkan *et al*., 2011; Jain *et al*., 2018), this is likely a conservative lower bound estimate; the ∼10% of the genome absent from our assembly is likely to consist of such regions rather than to reflect missing gene content (**Figure S3c**). This contrasts sharply with the compact genome of *S. polyrhiza*, in which TEs account for only ∼20.5% of the genome (**Figure 2a**) (Dombey *et al*., 2025).

Retrotransposons (RTE, Class I) dominate the *W. brasiliensis* genome, accounting for ∼57% of assembled sequence. Long terminal repeat (LTR) retrotransposons (LTR-RTEs) are particularly abundant, with *Ty3/Gypsy* and *Ty1/Copia* elements representing the largest superfamilies (15.37% and 13.73% of the genome, respectively; **Figure 2b**; **Figure S3d**). To assess the TE accumulation history, we estimated divergence from family consensus sequences using Kimura substitution levels (Kimura, 1980; SanMiguel *et al*., 1998). Approximately 3.5% of the genome consists of TE copies showing no detectable divergence from their consensus, indicative of very recent insertions. Other copies are moderately diverged, consistent with sustained TE activity over time and dominated by LTR-RTE families (**Figure 2c**). The much smaller fraction of TEs in the *S. polyrhiza* genome lacks a substantial fraction of moderately diverged (1–5%) elements observed in *W. brasiliensis* (**Figure S4a**), suggesting distinct accumulation histories.

Structural analyses further support recent activity of LTR-RTEs in *W. brasiliensis*. A substantial fraction of LTR-RTEs exceed 4 kb in length (**Figure S4b**), consistent with relatively intact insertions. Functional domain annotation confirmed that *Ty1/Copia* and *Ty3/Gypsy* elements longer than 4 kb encode complete sets of protein domains characteristic of autonomous LTR-RTEs (Neumann *et al*., 2019) (**Figure 2d**). Moreover, many intact elements display high (>98%) 5′–3′ LTR sequence identity (**Figure 2e, Figure S4c**), a hallmark of recent insertion (SanMiguel *et al*., 1998; Ma and Bennetzen, 2004). In contrast, the few intact LTR-RTEs detected in *S. polyrhiza* exhibit lower LTR identity values (**Figure 2e**). Together, these analyses indicate that the *W. brasiliensis* genome has undergone extensive and relatively recent TE proliferation, particularly of LTR-RTE, generating a repeat-rich genome.

### Pervasive TE integration reshapes gene organization in *W. brasiliensis*

We next examined how TE proliferation reshaped genome architecture in *W. brasiliensis*. Unlike *S. polyrhiza*, where TEs are largely confined to discrete TE-rich regions (Dombey *et al*., 2025), the *W. brasiliensis* genome is uniformly TE-rich, with genes distributed across contigs (**Figure 2f**). Consequently, genes in *W. brasiliensis* are more widely spaced than in *S. polyrhiza*, with significantly increased intergenic distances (**Figure 2g**). Moreover, many genes are located in close proximity to, or directly harbor, TE insertions (**Figure 2h**). Thus, TE proliferation in *W. brasiliensis* is associated not only with increased genome size but has also altered both genome-wide gene spacing and local genic environments.

### Transposable elements are major sources of 22- and 24-nt siRNAs in *W. brasiliensis*

Given the pervasive and recent TE amplification in *W. brasiliensis*, we asked how this impacts small RNA production. To validate the lineage-wide enrichment of 24-nt siRNAs observed in the initial survey, we generated additional small RNA sequencing libraries from both total RNA and AGO-loaded (TraPR-purified) fractions. As 20–25-nt small RNAs represent only a minor fraction of total libraries (**Figure S5a**), we focused subsequent analyses on AGO-loaded populations in which approximately half of all mapped reads aligned to annotated TEs. While 21-nt small RNAs were largely derived from microRNA (miRNA) loci, 24-nt siRNAs were enriched over TEs. Notably, TE-derived 22-nt siRNAs were also abundant and constituted the most prevalent size class among TE-mapping small RNAs in AGO-loaded datasets (**Figure 3a**), contrasting with data from most angiosperms in which TE-derived small RNAs are typically dominated by the 24-nt class (Patel *et al*., 2021).

**Figure 3.**
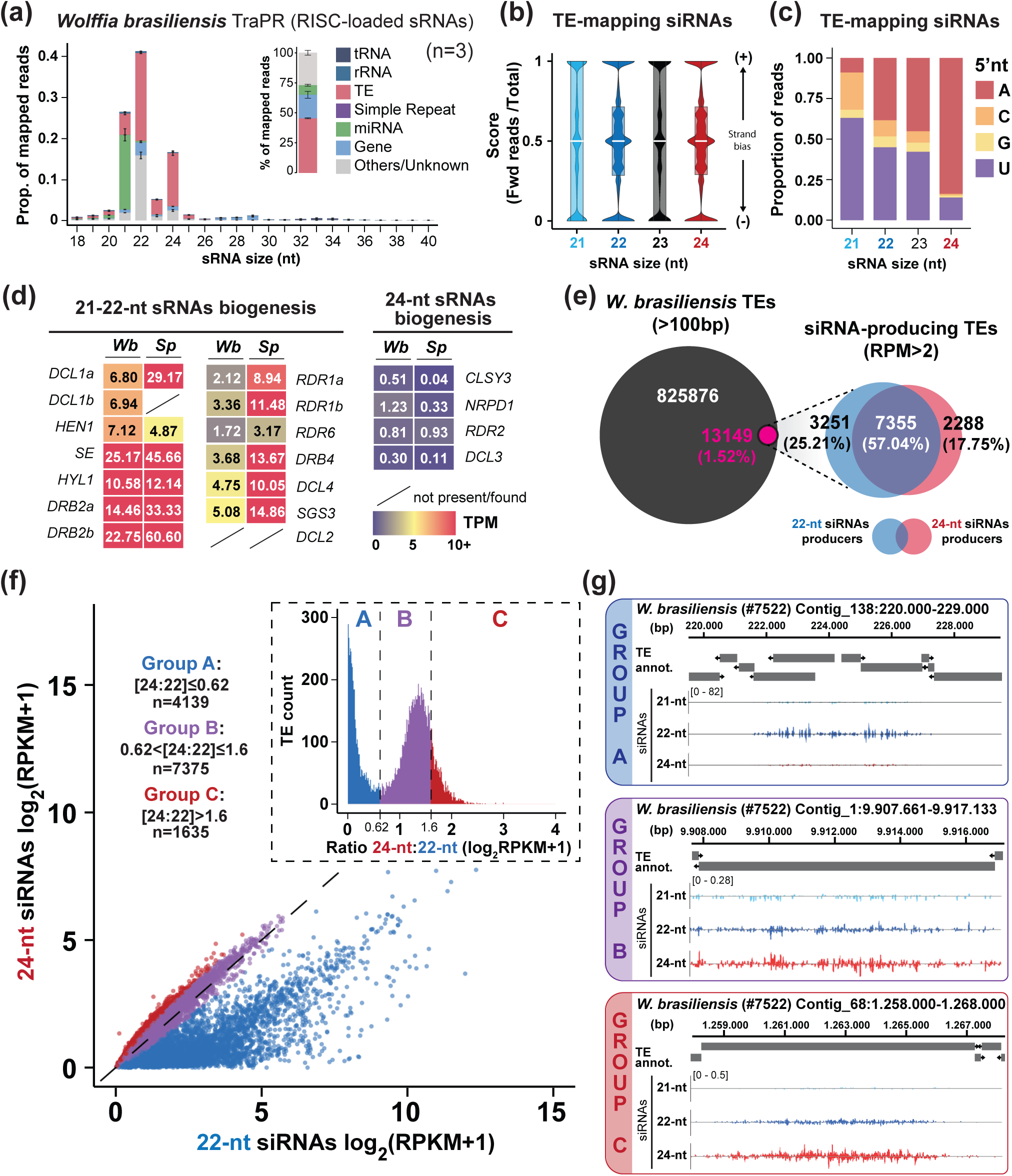
Analysis of TE-derived small RNA in *W. brasiliensis*. **(a)** Size distribution and genomic mapping of TraPR-purified small RNAs. Inset shows percentage (%) of all mapped siRNA per feature. Bars and error bars indicate the mean and the SD respectively (n=3). **(b)** Strand bias distribution of TE mapping siRNAs across all TE loci. A strand-bias score (0-1) for each TE locus shown as the ratio of forward strand (+) mapping reads to total mapped reads independently of the orientation of the TE annotation. A score of 0 indicates exclusive mapping to the reverse strand (-), while a score of 1 indicates exclusive mapping to the forward strand (+). **(c)** 5’ nt bias distribution of TE-mapping siRNAs. **(d)** Expression in transcripts per million (TPM) of small RNA biogenesis components in *S. polyrhiza* (*Sp*) and *W. brasiliensis* (*Wb*). **(e)** Venn diagrams of siRNA-producing (>2 reads per million; RPM) over total TE annotations and overlap of 22- and 24-nt siRNA-producing TEs. **(f)** Abundance (in reads per kb per million reads; RPKM+1) of 22- and 24-nt siRNAs for each siRNA-producing TE colored by groups determined by their 24-nt:22-nt ratio as defined in inlet. Diagonal line represents the 1:1 ratio. **(g)** Genome browser captures of selected examples of TEs in Groups defined in (f). siRNA values in RPM.

TE-derived small RNAs displayed canonical features of plant siRNAs. Their genomic mapping was balanced between the forward and reverse strands independently of the annotated TE orientation, consistent with double-stranded RNA precursors, in clear contrast to miRNAs, which showed strong strand bias (**Figure 3b**, **Figure S5b,c**), and exhibited size-dependent 5′ nucleotide biases consistent with the known loading preferences of *A. thaliana* AGOs (Mi *et al*., 2008; Bologna and Voinnet, 2014), supporting functional AGO loading. miRNAs and TE-derived 21-nt siRNAs were enriched for 5′ uridine (5’U, AtAGO1-associated), whereas 24-nt siRNAs preferentially carried a 5′ adenosine (5’A, AtAGO4-associated). The 22-nt siRNA population showed a mixed 5′ composition, with predominant 5′U and 5’A biases (**Figure 3c**, **Figure S5b**), suggesting engagement with both PTGS- and RdDM-associated AGOs (AGO1 and AGO4, respectively). However, as several AGOs share 5’nt preferences (Mi *et al*., 2008; Bologna and Voinnet, 2014) and those of *W. brasiliensis* AGOs have not been determined, the exact AGOs involved cannot be fully assigned based on the observed 5’nt biases and might be contributed by other AGOs than WbAGO1 and WbAGO4.

### 22-nt siRNAs accumulate in *W. brasiliensis* despite no detectable DCL2 orthologue

The prevalence of TE-derived 22- and 24-nt siRNAs, largely absent in *S. polyrhiza* (Dombey *et al*., 2025), prompted us to examine the composition and expression of small RNA biogenesis pathways in *W. brasiliensis*. We identified orthologues of genes encoding known biogenesis factors and compared them to those of *S. polyrhiza* and *A. thaliana* (**Figure S6**).

The predominance of TE-derived 22-nt siRNAs in *W. brasiliensis* contrasts with the 24-nt bias typical of TE-derived siRNAs in most angiosperms and raised the question of whether altered mechanisms in small RNA biogenesis contribute to this shift. In angiosperms, 22-nt siRNAs are primarily generated by DICER-LIKE 2 (DCL2) (Xie *et al*., 2004; Gasciolli *et al*., 2005; Wang *et al*., 2018; Katsarou *et al*., 2019; Berube *et al*., 2024). A gene encoding a corresponding protein has not been detected in previously sequenced duckweed genomes (Ernst *et al*., 2025). We therefore examined the complement of DCL genes encoded in the *W. brasiliensis* genome. Four DCL genes were identified: two tandemly duplicated DCL1 paralogues, one DCL4, and one DCL3 (**Figure S7, S8**). Phylogenetic analysis confirmed the absence of DCL2 and validated the assignment of the remaining DCLs (**Figure S9**). Given that our assembly represents ∼90% of the estimated genome size, we cannot fully exclude that a highly diverged or lowly expressed DCL2 orthologue escaped detection. However, no evidence for such a gene was previously detected in any other duckweed, and DCL2 was also not recovered from a de novo transcriptome assembly of RNA-seq reads unmapped to the genome (**Figure S3c, Table S1**), further supporting its absence from *W. brasiliensis*.

In addition to the absence of DCL2, *W. brasiliensis* DCL3 exhibited less similarity to its *Spirodela* counterpart and lacks the conserved N-terminal helicase domain (Margis *et al*., 2006; MacRae and Doudna, 2007) (**Figure S7**, **S10a–c**). Analysis of the genomic locus and supporting transcriptomic data confirmed that this truncation is not attributable to mis-annotation (**Figure S10d**).

Despite these structural differences, the overall complement and expression levels of genes involved in 21–22-nt siRNA biogenesis were broadly comparable to those observed in *S. polyrhiza* (**Figure 3d**; **Figure S11**) (Dombey *et al*., 2025). Thus, the accumulation of 22-nt siRNAs in *W. brasiliensis* in the absence of a DCL2 orthologue suggests altered functionality of conserved pathways.

### Elevated 24-nt siRNA are abundant despite reduced biogenesis factors

Given the elevated abundance of 24-nt siRNAs in *W. brasiliensis* relative to *S. polyrhiza*, we next examined whether this enrichment reflects expansion or upregulation of canonical RdDM components, focusing on genes associated with 24-nt siRNAs biogenesis. Consistent with previous analyses of duckweed genomes (Dombey *et al*., 2025; Ernst *et al*., 2025), several conserved components of this pathway are absent in *W. brasiliensis*. Notably missing are *CLASSY 1* (*CLSY1*) and *SHAWADEE HOMEODOMAIN HOMOLOGUE 1* (*SHH1*), which in *A. thaliana* promote 24-nt siRNA production from TEs located in gene-rich chromosomal arms (Law *et al*., 2013; Zhou *et al*., 2022). The retained RdDM-associated genes—including CLSY3, NUCLEAR RNA POLYMERASE D1 (NRPD1), RNA-DEPENDENT RNA POLYMERASE 2 (RDR2), and DCL3, were expressed at markedly lower levels relative to *A. thaliana*, closely mirroring expression patterns observed in *S. polyrhiza* (**Figure 3d**; **Figure S11**) (Dombey *et al*., 2025). Together with the above results, these observations indicate that altered small RNA size distributions in *W. brasiliensis*, relative to *S. polyrhiza*, likely arise from differential deployment of the reduced, but conserved silencing principles.

### Individual transposable elements generate overlapping 22- and 24-nt siRNA populations in *W. brasiliensis*

To determine whether 22- and 24-nt siRNAs originate from distinct or overlapping TE populations, we quantified siRNA production at individual annotated TEs. Only 1.52% of annotated TEs produced detectable levels of siRNAs, but with substantial overlap: 57% generated both 22- and 24-nt siRNAs (**Figure 3e**), suggesting that the two size classes frequently originate from the same elements. To characterize this overlap, we classified siRNA-producing TEs according to the ratios between 24-nt and 22-nt siRNAs. This analysis identified three groups: Group A, strongly biased toward 22-nt production; Group B, producing comparable levels of 22- and 24-nt siRNAs; and Group C, modestly biased toward 24-nt siRNAs (**Figure 3f**-**g**; **Figure S12a**). Restricting the analysis to uniquely mapping siRNAs yielded similar patterns (**Figure S12b**), excluding the possibility that apparent overlap resulted from multi-mapping reads assigned to different insertions of the same TE family.

To test whether the overlap of siRNA size classes at individual TEs extends beyond *W. brasiliensis*, we applied the same framework to *S. polyrhiza* (**Figure S13**). Despite the markedly lower overall abundance of TE-derived siRNA (**Figure S13a-c**), siRNA producing TEs frequently generated overlapping multiple size classes (**Figure S13d**), confirming that this feature is shared with *W. brasiliensis*. To accommodate the predominant 21- and 22-nt species at *S. polyrhiza* TEs (**Figure S13b,d**), the group defining ratio was calculated as 24-nt:(21+22)-nt while keeping the threshold values identical to those used in *W. brasiliensis* (**Figure S13e**).

We next asked whether specific TE families preferentially associate with distinct siRNA profiles. Because the smaller TE complement of *S. polyrhiza* yields markedly fewer siRNA-producing TEs per family and per group, this analysis was restricted to *W. brasiliensis.* Among all siRNA-producing TEs, several superfamilies were enriched relative to their genomic representation, including LTR-RTEs *Copia* (RLC), *Gypsy* (RLG), non-LTR RTE (RIX), DNA transposons of the *hAt* (DTA) and *Mutator* (DTM) superfamilies, indicating that siRNA production is restricted to specific classes of elements (**Figure S14**). However, when stratified by siRNA bias groups, no TE superfamily was exclusively associated with a particular size-class profile. Although some families were enriched among siRNA-producing elements, such as MITEs in Group A, enrichment patterns varied across groups, implying that family identity alone does not determine siRNA output (**Figure S14**). Together, these results indicate that many siRNA-producing TEs in *W. brasiliensis* generate overlapping 22-and 24-nt populations, a pattern of overlapping size classes also observed at siRNA-producing TEs in *S. polyrhiza*, and that variation in siRNA size bias is only partially explained by TE family identity, suggesting additional structural or contextual determinants.

### Canonical PTGS substrates are predominantly processed into 22-nt siRNAs in *W. brasiliensis*

As Group A TEs produce predominantly 22-nt siRNAs, typically associated with PTGS, we asked whether they represent PTGS substrates in *W. brasiliensis*. However, in most angiosperms PTGS substrates are primarily processed into 21-nt siRNAs by DCL4, with 22-nt species typically arising from DCL2 activity (Gasciolli *et al*., 2005; Bologna and Voinnet, 2014). Given the absence of DCL2, we first examined how well-established PTGS substrates are processed in this species.

We first analyzed endogenous TAS3 loci, trans-acting siRNA (tasiRNA)-producing genes conserved in plants and initiated by miR390 (Axtell *et al*., 2006; Xia *et al*., 2017). Using conserved miR390 target complementarity, we identified two TAS3 loci in the *W. brasiliensis* genome (TAS3a and TAS3b), each containing the canonical dual miR390 target sites (**Figure S15**). Both loci produced siRNAs from the expected regions **(Figure 4a**). However, tasiRNAs were dominated by 22-nt species, with minimal representation of other size classes (**Figure 4a,b**). TAS3a-derived 22-nt tasiRNAs showed the expected 5′U bias associated with AGO1 loading, whereas TAS3b-derived tasiRNAs exhibited a strong 5′G bias (**Figure 4c**). Although a plant AGO with strict 5′G preference has not been described, a similar bias has been observed for 5S rRNA–derived siRNAs in *S. polyrhiza* (Dombey *et al*., 2025), suggesting precursor or AGO-loading specific properties in duckweeds.

**Figure 4.**
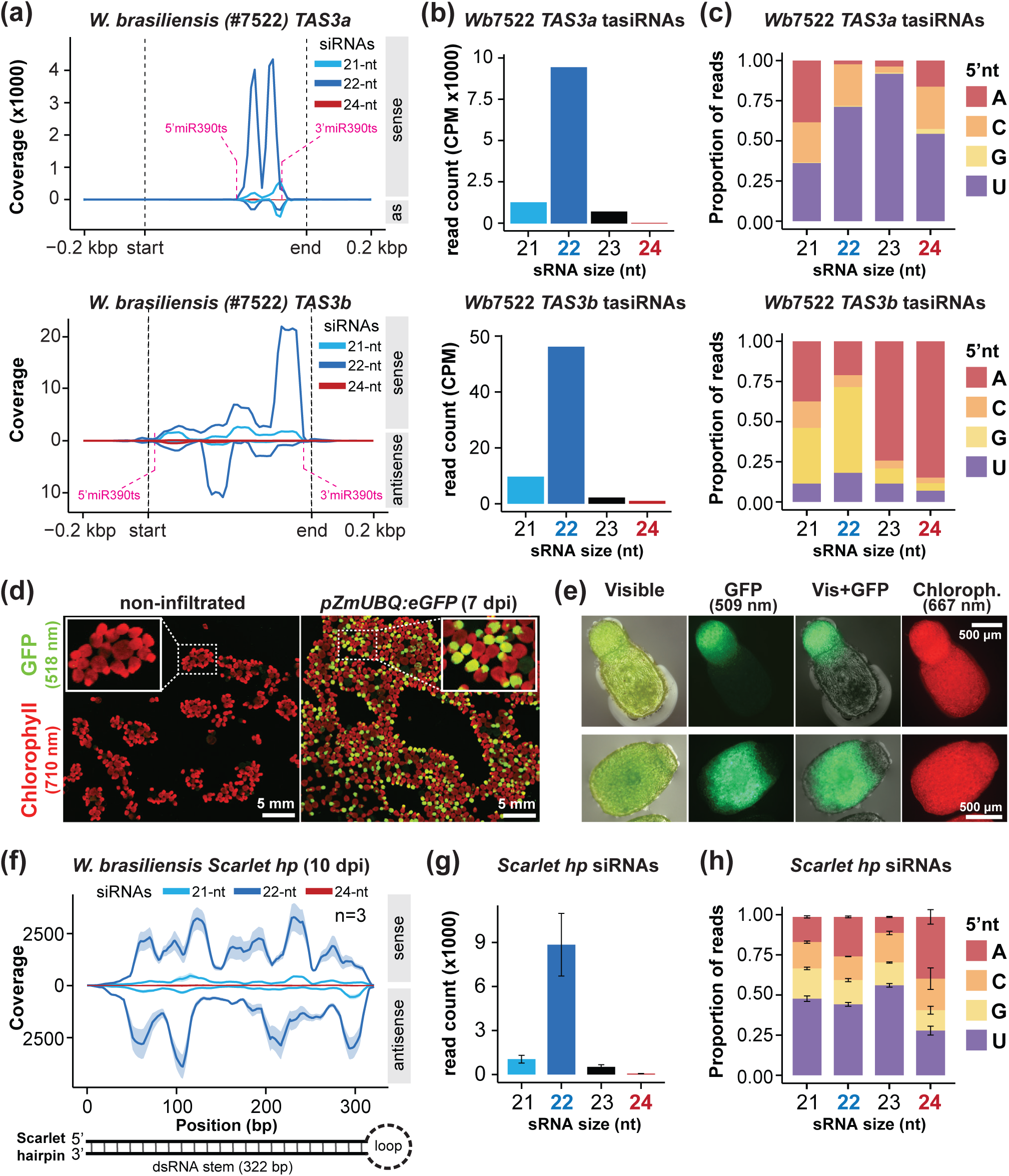
PTGS substrates are processed as 22-nt siRNAs in *W. brasiliensis*. **(a)** Metaplots of normalized siRNA coverage on the *TAS3* loci and their flanking regions. Pink dashed lines indicate the miR390 target sites (ts). **(b-c)** Size distribution and abundance (b), and 5’ nt bias (c) of *TAS3* mapping 21-24-nt siRNAs. **(d)** Representative images of *W. brasiliensis* of non-transformed and fronds showing transient GFP expression at 7 days post infiltration (dpi) obtained in a biomolecular scanner. **(e)** Representative fluorescent microscopy images of GFP-expressing fronds at 7 dpi. **(f)** Metaplot of siRNA distribution across *Scarlet* hairpin (hp) transiently expressed in *W. brasiliensis* at 10 dpi. Solid line and shaded area represent the average of the non-normalized coverage calculated from 3 independent replicates and the SD, respectively. **(g-h)** Size distribution and abundance (g), and 5’ nt bias (h) of *Scarlet hp* mapping 21-24-nt siRNAs. Bars and error bars indicate the mean and the SD respectively (n=3).

To test PTGS processing independently of endogenous loci and in absence of stable transformation procedures, we employed a transient expression system adapted from *S. polyrhiza* (Dombey *et al*., 2025; Barragán-Borrero *et al*., 2026). Expression of an eGFP reporter demonstrated robust and sustained transgene activity in *W. brasiliensis* (**Figure 4d,e**; **Figure S16**). Using this system, we expressed a 300-bp RNA hairpin (*Scarlet hp*), another well-established PTGS substrate in plants (Fusaro *et al*., 2006; Parent *et al*., 2015), previously shown to generate mixed 21- and 22-nt siRNAs in *S. polyrhiza*. In TraPR libraries from *W. brasiliensis*, the hairpin sequence was almost exclusively present as 22-nt siRNAs, with a predominant 5′U bias (**Figure 4f-h**).

To exclude that the 22-nt siRNAs arose from non-templated nucleotide addition to canonical 21-nt products (Wang *et al*., 2016), we examined mismatch patterns in uniquely mapping reads (**Figure S17**). Only 1.85% of reads carried a 3′-terminal mismatch, the position where non-templated nucleotides are added, lower than the internal-position mismatch rate (5.38%), while 92.19% matched the genome perfectly. This 3′ rate is comparable to the ∼2.6% reported for genuine DCL products and is far below the ∼35% reported for small RNAs one nucleotide longer than their dominant size, in other plants (Wang *et al*., 2016). Therefore, the 22-nt siRNAs represent primary DCL products.

Together, these results show that canonical PTGS substrates in *W. brasiliensis* are predominantly processed into 22-nt siRNAs despite the absence of DCL2, supporting a 22-nt–biased PTGS regime in this species.

### Inverted repeats define a subset of highly productive 22-nt–biased TEs in *W. brasiliensis*

In many plant genomes, TE insertions can generate inverted repeat (IR) configurations capable of forming long hairpin structures that serve as potent substrates for DCL processing (Slotkin *et al*., 2005; Chen *et al*., 2011; Gao *et al*., 2012). Because PTGS is typically triggered by double-stranded RNA, we then investigated whether structural configurations within Group A TEs could intrinsically generate such substrates.

Visual inspection of a representative Group A locus in *W. brasiliensis* revealed symmetric distribution of 22-nt siRNAs, suggestive of IR organization (**Figure 5a**). Sequence analysis of the region confirmed the presence of a large IR structure (**Figure S18**), predicted to form a ∼2.9 kb near-perfect hairpin with extensive double-stranded RNA stem (**Figure 5b**). Similar siRNA patterns and IR structures were observed at additional Group A loci (**Figure S19**), indicating that IR structures may underlie 22-nt–biased siRNA production at a subset of elements.

**Figure 5.**
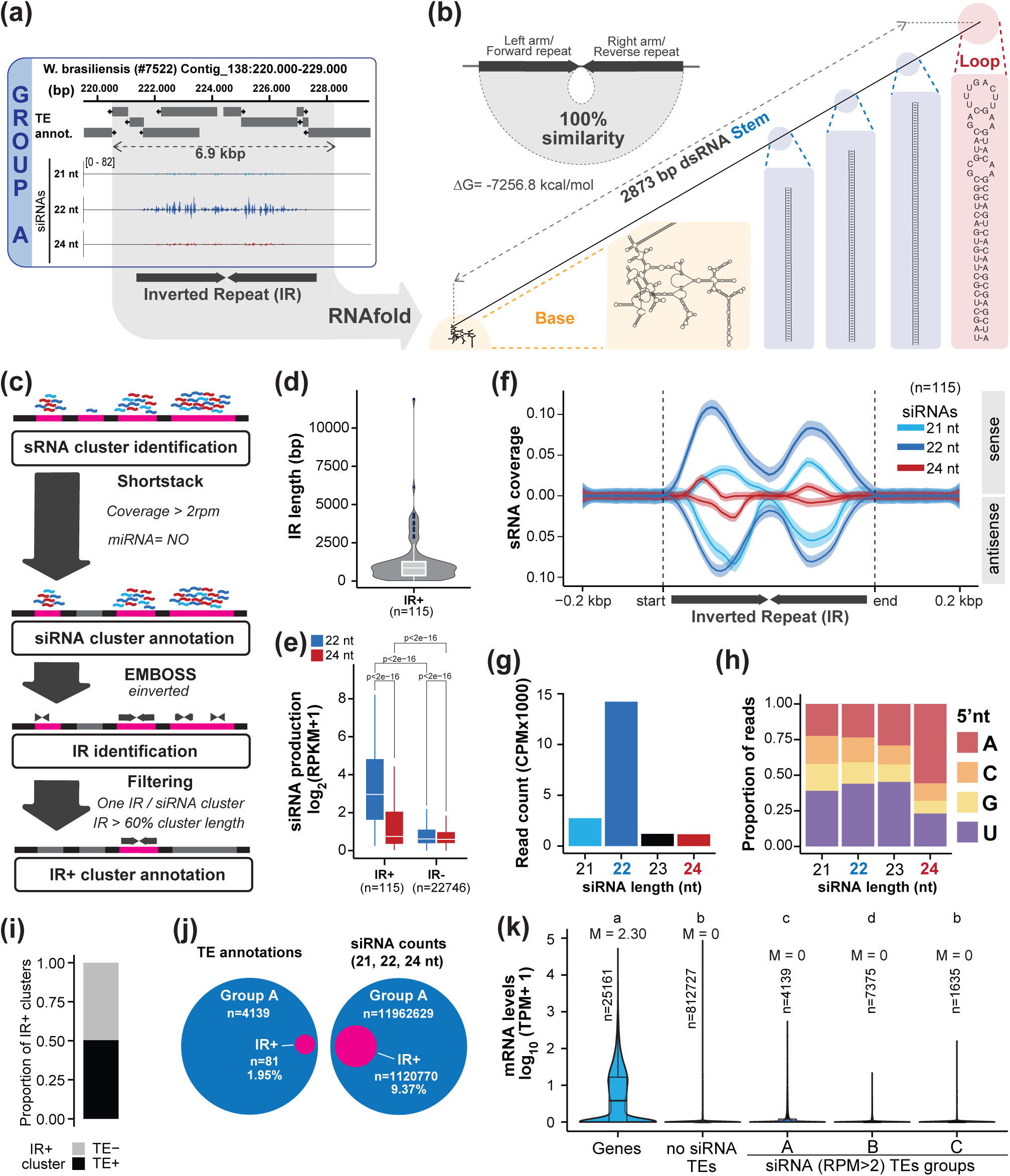
Identification and characterization of endogenous inverted repeats in *W. brasiliensis*. **(a)** Genome browser capture of the Group A locus sown in Figure 3g. **(b)** Secondary structure analysis of the locus shown in (a). **(c)** Schematic representation of the workflow strategy used to identify and annotate inverted repeat-containing (IR+) siRNA clusters. **(d)** Length distribution in bp of identified IRs. **(e)** 22-and 24-nt siRNA levels (RPKM+1) of IR+ and IR- siRNA clusters. P-values determined by Wilcoxon rank sum test. **(f)** siRNA distribution across identified IRs and their flanking regions. Solid line and shaded area represent the mean normalized coverage and the SD respectively. **(g-h)** Size distribution and abundance (g), and 5’ nt bias (h) of IR+ cluster-derived siRNAs. **(i)** Proportion of IR+ siRNA clusters overlapping with TE annotations **(j)** Venn diagrams of Group A TEs overlapping with IR+ clusters and their contribution to Group A-mapping siRNAs. **(k)** Expression level (TPM) comparison between genes and TEs grouped by siRNA outcomes. Different letters indicate significant differences between groups (Kruskal-Wallis followed by Dunn’s test, p<0.05). All P-values can be found in Table S2. M, median values.

To assess the genome-wide contribution of IRs to siRNA production, we systematically searched siRNA clusters for inverted repeat configurations. Starting from annotated siRNA clusters, we defined IR-positive (IR+) clusters as those containing a single inverted repeat encompassing more than 60% of the siRNA-producing region (**Figure 5c**). This stringent filtering identified 115 IR+ clusters, spanning a broad size range (**Figure 5d**). IR+ clusters were significantly more productive than those classified as lacking IRs (IR-) (**Figure 5e**). siRNA accumulation was concentrated over the inverted repeat arms and was composed almost exclusively of 22-nt species with predominant 5′U bias (**Figure 5f-h**), mirroring the patterns observed for Group A TEs and the *Scarlet hp* (**Figure 4f-h**, **Figure S12a**).

Intersection with TE annotations revealed that approximately half of IR+ clusters overlapped annotated TEs (**Figure 5i**). Although TEs in IR+ loci represented only ∼2% of Group A elements, they contributed ∼9.5% of all siRNAs derived from Group A TEs (**Figure 5j**). Thus, IR–forming TEs represent a minor but highly productive subset of 22-nt siRNA–biased loci.

An equivalent IR analysis in *S. polyrhiza* identified 77 IR+ clusters that produced significantly higher levels of 21- and 22-nt siRNAs than IR- clusters (**Figure S20a,b**), with comparable siRNA accumulation patterns over IR arms (**Figure S20c,d**). Although less that 25% of IR+ clusters overlapped with TE annotations and those represented only 1.8% of *S. polyrhiza* Group A elements, they contributed to about 37% of all Group A siRNAs (**Figure S20e,f**), echoing the disproportionate contribution of TE-derived IRs to siRNAs within this group observed in *W. brasiliensis*. This pattern is also consistent with the small RNA patterns observed in transiently expressed IR previously described in *S. polyrhiza* (Dombey *et al*., 2025), supporting the same IR-mediated PTGS-like framework in both species.

Because PTGS is typically associated with transcriptional activity, we examined whether Group A TEs in *W. brasiliensis* exhibited elevated steady-state mRNA levels. Although PTGS lowers steady-state transcript levels, in *A. thaliana* TEs subject to PTGS that produce 21–22-nt siRNAs nonetheless retain readily detectable steady-state mRNA, since their silencing depends on sustained transcription (Marí-Ordóñez *et al*., 2013; Creasey *et al*., 2014; Oberlin *et al*., 2022). Steady-state mRNA therefore offers an informative, if conservative, proxy for their transcriptional activity. However, Group A steady-state transcript levels were not markedly increased relative to other siRNA-producing or non-producing TEs (**Figure 5k**), and 22-nt siRNA abundance and TE mRNA abundance were only weakly correlated (**Figure S21**).

Together, these analyses indicate that IR structures constitute a prominent structural determinant of highly productive 22-nt siRNA loci in *W. brasiliensis* and similarly 21-22-nt siRNAs in *S. polyrhiza*. For the broader Group A TE population, the weak correlation between 22-nt siRNAs and steady-state mRNA levels may reflect either a transcriptional contribution that remains below detection, owing to ongoing PTGS-mediated mRNA degradation, or additional mechanisms contributing to 22-nt siRNA generation at these loci.

### Widespread CG methylation characterizes both TEs and genes in *W. brasiliensis*

Because 24-nt siRNAs are canonically associated with RdDM and 22-nt siRNAs with PTGS, we next asked whether the distinct TE classes defined by siRNA output exhibit corresponding differences in DNA methylation and examined genome-wide DNA methylation patterns. Unlike *S. polyrhiza*, which displays globally less DNA methylation largely confined to limited TE-rich regions (Dombey *et al*., 2025), *W. brasiliensis* exhibited elevated CG methylation (mCG) broadly distributed across the genome. By comparison, non-CG methylation (mCHG and mCHH) remained globally low (**Figure 6a**). At the level of TEs, mCG was uniformly high, reaching ∼75% on average (**Figure 6b**). In contrast, mCHG and mCHH at TEs was not elevated compared to the overall level. This pattern differs markedly from that in *S. polyrhiza*, in which TE-associated methylation levels are substantially lower across all contexts (**Figure 6b**).

**Figure 6.**
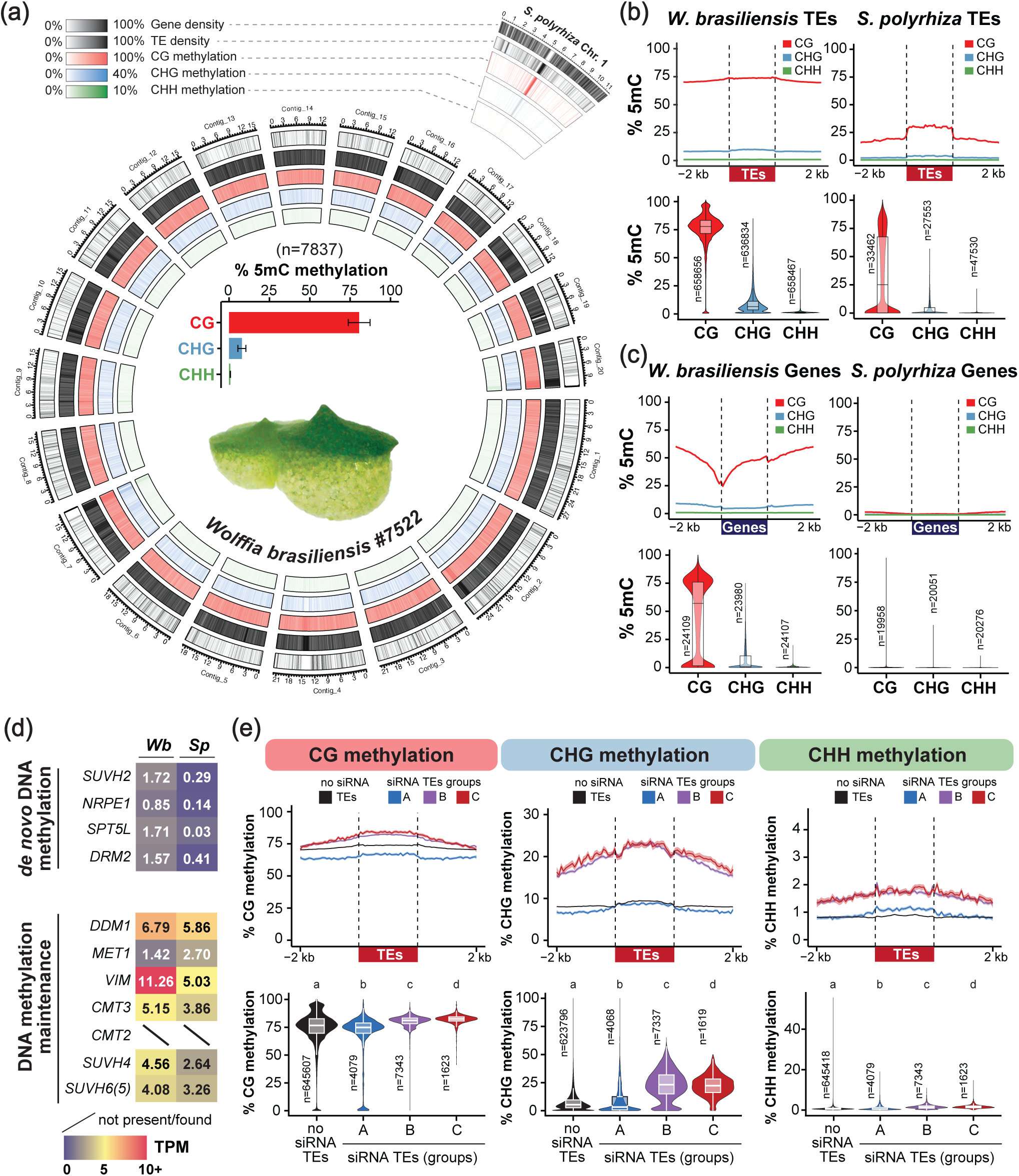
Distribution of DNA methylation in *W. brasiliensis*. **(a)** Circos plot of gene/TEs density as in Figure 2f and 5mC for CG, CHG, or CHH contexts (from outside to inside) across the 20 longest *W. brasiliensis* contigs, using 100 kb windows. *S. polyrhiza* chromosome 1 is shown for comparison. Center: Global DNA methylation levels per context. Bars and error bars indicate the mean and the SD from the mean respectively, across all genomic windows. **(b-c)** Metaplots of average weighted DNA methylation and 5mC distribution in all three contexts for TEs (b) and genes (c) in *S. polyrhiza* and *W. brasiliensis*. **(d)** Expression in TPM of genes involved in DNA methylation in *S. polyrhiza* (*Sp*) and *W. brasiliensis* (*Wb*). **(e)** Metaplots of average weighted DNA methylation and 5mC distribution in all three contexts for TEs grouped by siRNA outcomes in *W. brasiliensis*. Different letters indicate significant differences between groups (Kruskal-Wallis followed by Dunn’s test, p<0.05). All P-values can be found in Table S3.

Gene-associated methylation also differed strikingly between species. Whereas most genes in *S. polyrhiza* lack detectable gene body methylation (gbM) (Harkess *et al*., 2024; Dombey *et al*., 2025; Ernst *et al*., 2025), *W. brasiliensis* displayed widespread gbM with substantial heterogeneity across loci. While some genes remained largely unmethylated, many exhibited mCG levels approaching those observed at TEs (**Figure 6c**).

To determine whether this methylation landscape reflects expansion or upregulation of the factors installing methylation, we examined the complement and expression of genes involved in methylation establishment and maintenance (Xie *et al*., 2025) (**Figure 6d**, **Figure S11**). Similar to *S. polyrhiza*, *W. brasiliensis* retains a streamlined set of DNA methylation components and lacks CMT2, which is absent in all duckweeds examined to date (Harkess *et al*., 2024; Dombey *et al*., 2025; Ernst *et al*., 2025). Genes associated with RdDM showed modest expression levels relative to *A. thaliana* (**Figure 6d**, **Figure S11**), consistent with the reduced number of RdDM factors described above. Within the pathway, however, the downstream, Pol V–dependent arm that directs de novo methylation (SUVH2, NRPE1, SPT5L and DRM2) was more expressed than in *S.polyrhiza* (**Figure 6d**, **Figure S11**) in contrast to the upstream 24-nt–biogenesis arm (NRPD1, RDR2 and DCL3) (**Figure 3d**, **Figure S11**). Genes involved in DNA methylation maintenance exhibited comparable or lower expression relative to *S. polyrhiza*. Notably, *MET1* expression was lower in *W. brasiliensis* despite the substantially higher mCG levels observed genome-wide, although its cofactor *VARIANT IN METHYLATION* (*VIM*) was more expressed (**Figure 6d**; **Figure S11**).

Together, these results reveal a methylome in *W. brasiliensis* characterized by pervasive CG methylation across both TEs and genes despite a methylation machinery that is largely shared with *S. polyrhiza*. This indicates that the divergent methylation landscapes of the two duckweed species are unlikely to reflect pathway divergence and motivates examining how the conserved methylation pathways behave when applied to the much larger TE complement of *W. brasiliensis*.

### 24-nt siRNA–producing TEs preferentially acquire non-CG methylation in both duckweed species

Given the pervasive CG methylation observed across TEs in *W. brasiliensis*, we next examined whether siRNA production predicts differences in DNA methylation at individual elements. TEs were again grouped according to their siRNA profiles (Groups A–C and non–siRNA-producing). CG methylation was uniformly high across most TEs. However, TEs producing 24-nt siRNAs (Groups B and C) exhibited significantly higher mCG levels than non–siRNA-producing elements, whereas Group A TEs, biased toward producing 22-nt siRNAs, displayed the lowest mCG levels among all categories (**Figure 6e**). Although these differences were modest relative to the overall high mCG background, they indicate quantitative variation in CG methylation linked to siRNAs.

More pronounced differences emerged in non-CG contexts. Despite globally low mCHG and mCHH levels in *W. brasiliensis*, TEs producing 24-nt siRNAs showed significantly elevated non-CG methylation relative to both Group A and non–siRNA-producing TEs (**Figure 6e**). In contrast, non-CG methylation was not elevated at Group A TEs, consistent with their predominant association with 22-nt siRNAs. Within 24-nt–producing elements, Group C TEs, characterized by a higher 24-nt:22-nt ratio, showed modest but significant increases in methylation across all sequence contexts compared with Group B (**Figure 6e**), suggesting that the relative dominance of 24-nt siRNAs correlates with strengthened methylation signatures. These results indicate that while CG methylation is broadly maintained across TEs in *W. brasiliensis*, elevated non-CG methylation is selectively associated with 24-nt siRNA production, consistent with targeted engagement of RdDM at a subset of TEs.

Applying the same framework to *S. polyrhiza* revealed that the coupling of 24-nt siRNAs to elevated methylation is conserved between the two species: as in *W. brasiliensis*, Group B and C TEs accumulated significantly higher methylation than non–24-nt-producing TEs (Group A and non-siRNA) in all three sequence contexts (**Figure S22**). The principal divergence between species lay outside the 24-nt-producing classes: Group A and non-siRNA TEs in *S. polyrhiza* did not display the high CG methylation observed at their *W. brasiliensis* counterparts (**Figure S22**), consistent with the globally lower TE methylation in *S. polyrhiza*. The conservation of this siRNA-group–methylation rule across both species indicates that the substantially higher overall non-CG methylation of *W. brasiliensis* TEs reflects the larger absolute number of 24-nt-producing TEs in this species rather than altered per-element RdDM efficiency. The pervasive CG methylation at non–24-nt-producing TEs in *W. brasiliensis*, by contrast, points to an additional siRNA-independent contribution, most likely maintenance methylation acting on the expanded TE complement.

### TE length and, in *W. brasiliensis*, gene proximity influence siRNA production

Because only a subset of TEs produce siRNAs and acquire elevated non-CG methylation in *W. brasiliensis*, we next sought to identify intrinsic and genomic features that distinguish these elements from the broader TE population. TE length emerged as a correlate shared between species: longer TEs were more heavily methylated overall in *W. brasiliensis* (**Figure 7a**; **Figure S23a,b**), in agreement with previous observations in *S. polyrhiza* (Dombey *et al*., 2025), and in both species the proportion of siRNA-producing TEs, and of 24-nt-producing Groups B and C in particular, rose with TE length (**Figure 7b,c**; **Figure S24a,b**). These analyses suggest that long, relatively intact, TEs are more likely to engage RdDM-associated pathways across duckweeds, independently of overall TE complement.

**Figure 7.**
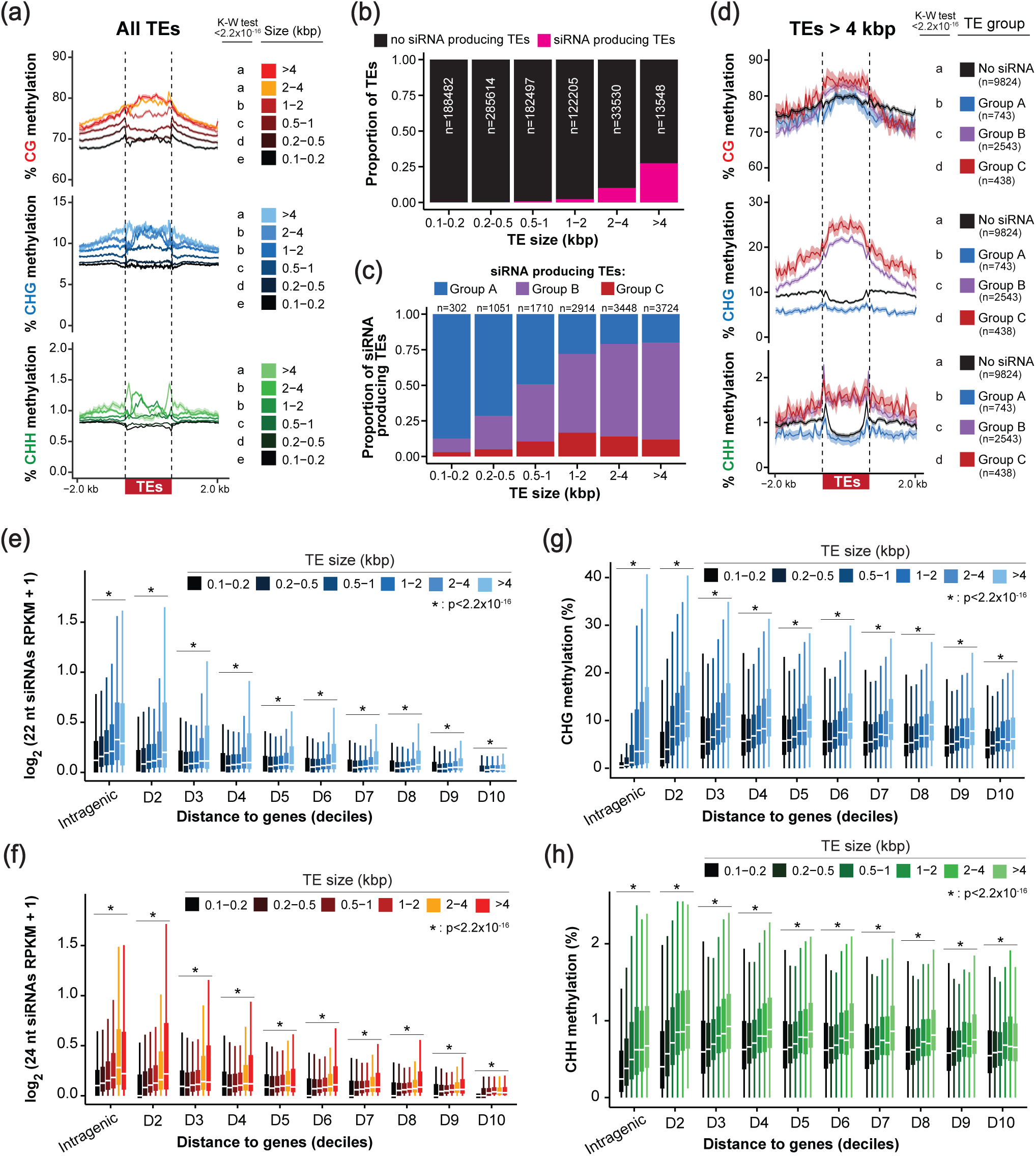
Impact of size and distance to genes on TE DNA methylation and siRNA production. **(a)** Metaplots of the average weighted 5mC in all three contexts for TEs grouped by size categories and their flanking regions. **(b-c)** Proportion of siRNA-producing TEs (b) and of TE groups by 24-nt:22-nt ratios (c), per size category. **(d)** Metaplots of the average weighted 5mC in all three contexts for TEs >4 kb and their flanking regions grouped by 24-nt:22-nt ratios. In (a) and (d), different letters indicate significant differences between groups (Kruskal-Wallis followed by Dunn’s test p<0.05) **(e-f)** TE 22-nt (e) and 24-nt (f) siRNA levels (RPKM+1) grouped by distance to genes (deciles) and TE sizes. **(g-h)** Average CHG (g) and CHH (h) methylation levels (%) at TEs grouped by distance to genes (deciles) and TE sizes. (*) P-values<2.2 x 10^-16^ (Kruskal-Wallis test) between size categories within each decile and intragenic group in (e-h). All P-values and n-values can be found in Tables S3, S4, S6.

However, TE length alone was not sufficient to predict siRNA production. Even among long elements (>4 kb), only slightly more than 25% produced detectable siRNAs in *W. brasiliensis* (**Figure 7b**) markedly lower than the 43% observed for comparably sized TEs in *S. polyrhiza* (**Figure S24a**), indicating other factors beyond length also shape this engagement. Within this long-TE subset, the association between siRNA output and methylation remained evident: TEs producing 24-nt siRNAs exhibited significantly higher non-CG methylation than either non–siRNA-producing or Group A TEs (**Figure 7d**, **Figure S23b**). Group C elements, characterized by higher 24-nt:22-nt ratios, displayed modest but significantly more methylation across all contexts relative to Group B. Thus, while TE length is an important correlate of siRNA production and methylation, it does not fully explain selective engagement of 24-nt–associated pathways.

We next examined whether genomic context contributes to this selectivity. In *A. thaliana* and maize, RdDM preferentially targets TEs located near genes, reinforcing chromatin boundaries at TE–gene interfaces (Gent *et al*., 2013; Zemach *et al*., 2013; Stroud *et al*., 2014; Gent *et al*., 2014; Li *et al*., 2015). TEs in *W. brasiliensis* were therefore partitioned according to distance to the nearest gene, including intragenic insertions (**Figure S25a**). The likelihood of TEs producing siRNAs—both 22- and 24-nt—is reduced with larger distance from genes (**Figure S25b**). TEs located within or immediately adjacent to genes accumulated higher siRNA levels. When TE length and gene proximity were considered jointly, a clearer pattern emerged: long TEs located near or within genes produced the highest siRNA levels across both size classes (**Figure 7e-f**). Together, these analyses indicate that both intrinsic properties (length and likely structural integrity) and genomic context (gene proximity) contribute to selective siRNA production in *W. brasiliensis*, with long, gene-proximal TEs showing the strongest engagement of 24-nt–associated methylation pathways.

To test whether the same gene-proximity bias operates in *S. polyrhiza*, we partitioned its TEs in the same way (**Figure S25a**). The TE-length effect was again observed, but siRNA production did not decline with distance from genes; instead, siRNA accumulation correlated with TE size independently of gene-TE distance (**Figure S25b,d,e**), and global 24-nt siRNA accumulation actually rose with distance, paralleling the increase in average TE size at gene-distal positions in this species (**Figure S25b,c**). The preferential targeting of gene-proximal TEs by siRNA-generating pathways is therefore a feature of *W. brasiliensis*, not shared with *S. polyrhiza*. Taken together, the analyses in this section distinguish two layers of TE selection by silencing pathways in duckweeds: an element-intrinsic component (length) shared across both species, and a genomic-context component (gene proximity) that operates in *W. brasiliensis* but not in *S. polyrhiza*.

### DNA methylation at intergenic TEs is largely decoupled from gene-distance and length gradients in *W. brasiliensis*

We then asked whether DNA methylation at TEs followed the same gene-proximity and size gradient. As observed genome-wide, CG methylation at TEs was consistently high across all distance-to-gene deciles and TE size classes (**Figure S26a**). However, although 22- and 24-nt siRNA accumulation declined with distance from genes, CHG and CHH methylation at intergenic TEs remained largely uniform across distance-to-gene deciles, and the same pattern held when TEs were jointly stratified by length (**Figure 7g,h; Figure S26b**). Nonetheless, within this overall uniformity, a gradient was apparent at the most gene-proximal intergenic TEs. There, non-CG methylation co-varied with TE length, like siRNAs, but this co-variation was progressively lost with increasing distance from genes, where methylation became homogeneous across TEs irrespective of length (**Figure 7g,h; Figure S26b**). Intragenic TEs were the exception: although they also displayed a similar co-variance of DNA methylation with size as proximal intergenic TEs, they showed lower non-CG methylation despite the production of siRNAs (**Figure 7e-h; Figure S26b**). Thus, DNA methylation at intergenic TEs is broadly decoupled from the gene proximity gradient that shapes siRNA production, with siRNA-associated variation in methylation mainly observed at the most gene-proximal TEs.

By contrast, in *S. polyrhiza* (**Figure S26c,d**), methylation at intergenic TEs co-varied with both TE length across all distance-to-gene deciles in all three sequence contexts, broadly tracking the patterns of siRNA production and without the homogenization observed at gene-distal positions in *W. brasiliensis*. The lower non-CG methylation of intragenic TEs relative to intergenic ones of similar size, however, was shared between the two species, indicating that intragenic constraint on RdDM-associated methylation is a common feature of the duckweed silencing landscape. This pattern of divergence, uniformity of methylation at gene-distal TEs in *W. brasiliensis* but preserved size-tracking in *S. polyrhiza*, fits the substrate-scaling framework raised earlier: in *W. brasiliensis*, the expanded TE substrate sustains a high CG-methylation baseline that masks or buffers the size-dependent siRNA-driven variation otherwise visible against the much lower TE-methylation baseline of *S. polyrhiza*.

### Intragenic TEs that produce siRNAs acquire little non-CG methylation in *W. brasiliensis* and *S. polyrhiza*

Given that intragenic TEs showed reduced non-CG methylation despite siRNA production, a feature shared between *W. brasiliensis* and *S. polyrhiza* (**Figure 7e-h**; **Figure S26b,d**), we next examined their siRNA and DNA methylation patterns in detail. Genome-wide, approximately 12% of annotated TEs in *W. brasiliensis* reside within genes, with nearly 75% of these insertions located in introns (**Figure 8a**), and more than half of all annotated genes (53%) contain at least one TE insertion (**Figure 8b**). The proportional partitioning of TEs between intergenic and intragenic compartments, and between intronic and exonic locations within gene bodies, was comparable in *S. polyrhiza* (**Figure S27a**); however, only ∼21% of its genes carry a TE insertion in this species. The most striking species divergence was found in intronic TE content: *W. brasiliensis* has substantially more intronic DNA than *S. polyrhiza* (74.30 vs 45.12 Mbp) and a much larger fraction of it is TE-derived (59.18% vs 7.56%; **Figure S27b,c**), corresponding to roughly an order-of-magnitude greater TE-derived intronic sequences in *W. brasiliensis*.

**Figure 8.**
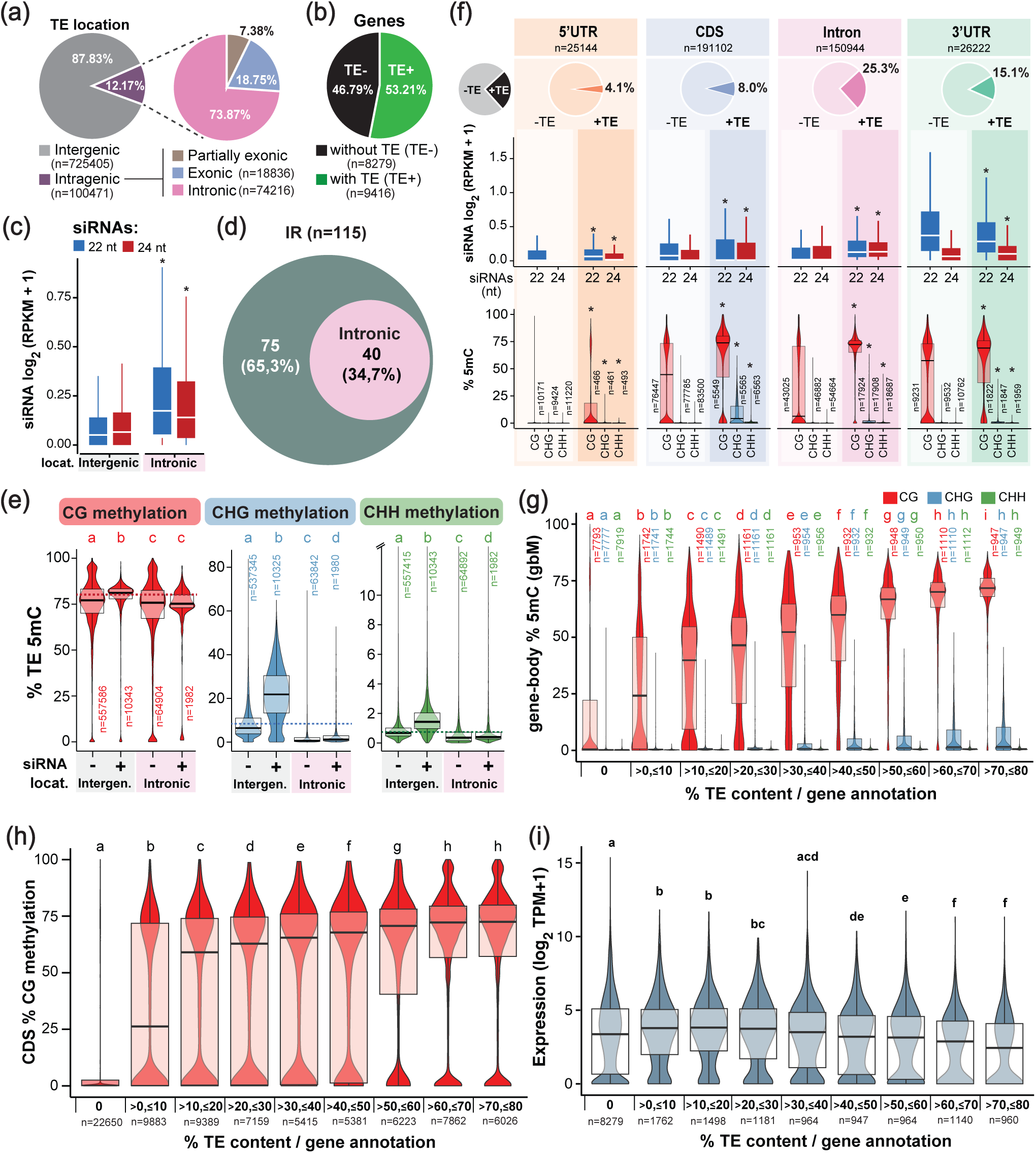
Impact of intragenic TEs in genic siRNAs and DNA methylation. **(a)** Percentage of intragenic TEs and their distribution within genes. **(b)** Proportion of genes overlapping with at least one TEs annotation. **(c)** siRNA levels (RPKM+1) levels of intergenic and intronic TEs. **(d)** Proportion of IR+ clusters within introns. **(e)** 5mC levels in all three contexts for siRNA- and non-siRNA-producing TEs in intragenic or intronic locations. Dashed line indicates average global methylation in each context. **(f)** siRNA and 5mC levels of different gene features (UTRs, CDSs, introns) overlapping (+) or not (-) with TE annotations. Pie charts show the proportion of TE-overlapping features. In (c) and (f), (*) indicates p<0.05 Wilcoxon rank-sum test between same siRNA size or 5mC context in different TE location (c) or TE+/TE- features (f). **(g)** 5mC levels in all three contexts for gene annotations without or within 10% incremental levels of relative TE content (sequence percentage over total annotation length). **(h)** mCG levels for CDS annotations within genes sorted by relative TE content as in (g). **(i)** Expression levels (TPM) for gene annotations sorted by relative TE content as in (g). Genes with a TE coverage higher than 80% were considered as TE genes and excluded from all analysis. In (e-i), different letters indicate significant differences between groups (Kruskal-Wallis followed by Dunn’s test p<0.05). All P-values can be found in Tables S7, S8, S9.

Due to the reduced number of intronic TEs in *S. polyrhiza*, we first investigated the relationship between siRNAs and DNA methylation in *W. brasiliensis.* To avoid confounding effects from misannotated TE-derived genes, we excluded “TE-genes” (∼30%), defined as gene models in which more than 80% of the gene body overlapped TE annotations (**Figure S28**) from the analysis. These models typically correspond to TE sequences for which no canonical TE protein domains have been identified and are therefore retained in the gene set despite their TE origin (Ou *et al*., 2019; Hufford *et al*., 2021). Intronic TEs accumulated higher levels of siRNAs than intergenic elements, including both 22- and 24-nt species, with a modest bias toward 22-nt siRNAs (**Figure 8c**). Notably, ∼35% of siRNA clusters encompassing IRs were located within introns (**Figure 8d**), including the long IR described above (**Figure 5a,b**, **Figure S29a**), indicating that intragenic contexts can support IR-driven siRNA production.

We next explored whether siRNA production correlates with DNA methylation, comparing intronic TEs, the predominant intragenic class (**Figure 8a**), with exonic and intergenic TEs. As observed genome-wide (**Figure 6b**), CG methylation remained high across TEs regardless of location; siRNA presence was associated with modestly elevated mCG at intergenic TEs but not at intronic TEs relative to their non–siRNA-producing counterparts (**Figure 8e**). Differences were more pronounced in non-CG contexts: intergenic TEs showed higher CHG and CHH methylation than intronic and exonic TEs, and siRNA production was associated with a clear increase in non-CG methylation at intergenic TEs and a much smaller, though still significant, increase at intronic TEs (**Figure 8e**, **Figure S29, Figure S30a**). Albeit less abundant, a similar pattern was observed for exonic TEs (**Figure S30a,b**).

Thus, although intragenic TEs frequently produce siRNAs, including 24-nt species, this yields only a limited gain of non-CG methylation relative to intergenic elements. An equivalent analysis in *S. polyrhiza* recapitulated this pattern: non-CG methylation at intragenic TEs was lower than at intergenic TEs producing siRNAs, although less than 200 intragenic TEs produce siRNAs in this species (**Figure S30c,d**). This low intragenic siRNA output (**Figure S30c**) likely reflects the substantially smaller number of long, intact TEs residing within gene bodies in *S. polyrhiza*. The shared reduction in intragenic non-CG methylation across both species therefore operates independently of overall TE-load, indicating that the intragenic environment constrains RdDM-associated methylation in *S. polyrhiza* and *W. brasiliensis*.

### Intragenic TE insertions are associated with gene body methylation in *W. brasiliensis*

Given that intragenic TEs frequently produce siRNAs, display high CG methylation, but acquire only limited non-CG methylation, we next examined how intragenic insertions influence gene-associated small RNA accumulation and DNA methylation patterns beyond the TE sequence itself. TE insertions within 5′ UTRs and introns were associated with modestly higher siRNA accumulation, whereas insertions within coding sequences (CDS) were associated with lower siRNA levels (**Figure 8f**). Elevated 22-nt siRNAs were also observed over 3′ UTRs even in the absence of annotated TE insertions (**Figure 8f**, **Figure S31**), indicating that gene-associated siRNA production is not exclusively TE-derived. In parallel, 3′ UTRs displayed high CG methylation regardless of TE presence (**Figure 8f**), correlating with this TE-independent siRNA accumulation. In contrast to the modest effects on siRNA levels, TE insertions were consistently associated with increased CG methylation across introns, CDS, and 3′ UTRs, except 5′ UTRs, which remained largely unmethylated (**Figure 8f**). We thus expanded the analysis to entire gene annotations. Genes lacking TE insertions displayed little or no gene body methylation (gbM), whereas increasing TE content within genes correlated with progressively higher mCG levels (**Figure 8g**, **Figure S32a**). Although non-CG methylation showed a similar trend, overall levels remained low (**Figure S32b**).

To ask whether high gbM is due to TE sequences within the gene annotation, we restricted the analysis to coding sequences (CDS). CDS methylation was nearly absent in genes without TE insertions but markedly elevated in genes containing TEs (**Figure 8h**, **Figure S33**), indicating that methylation extends beyond TE-derived sequence into adjacent coding regions. Within each TE-content category, CDS methylation levels were heterogeneous: some remained weakly methylated, whereas others exhibited high mCG levels. However, the proportion of highly methylated CDS increased with TE content (**Figure 8h**, **Figure S34**), indicating progressive impact of TE content on gbM rather than uniform elevation across all genes. By contrast, in *S. polyrhiza* no comparable relationship between gene methylation and TE coverage was apparent in any of the three sequence contexts (**Figure S35**), likely reflecting the substantially lower fraction of intronic sequence occupied by TEs in this species (**Figure S27b,c**) or that additional, *Wolffia*-specific features contribute to TE-driven gbM. Together, these analyses demonstrate that intragenic TE insertions are strongly associated with the emergence and quantitative enrichment of gene body CG methylation in *W. brasiliensis*, extending beyond the TE sequence itself. Despite these methylation changes, gene expression was detected across all TE-content categories. However, genes with high TE content and elevated CG methylation were enriched for lowly expressed transcripts, although low expression was not restricted to TE-containing genes and was also found among those lacking TE insertions (**Figure 8i**).

Collectively, our cross-species analyses indicate that *W. brasiliensis* and *S. polyrhiza* share the same per-element rules governing small RNA biogenesis and DNA methylation: TE length predicts siRNA-producing capacity, inverted-repeat-forming TEs underpin highly productive PTGS-associated siRNAs, 24-nt-producing TEs preferentially acquire non-CG methylation, and intragenic TEs face an intrinsic constraint on RdDM-associated methylation. Against this conserved framework, *W. brasiliensis* displays several genome-wide outcomes not observed in the smaller-TE-load *S. polyrhiza*: pervasive CG methylation across the TE complement, gene-proximity targeting of RdDM at intergenic TEs, and gene body CG methylation that correlates with intragenic TE content.

## Discussion

We integrated genome, methylome, and small RNA analyses of the TE-dense *W. brasiliensis* with cross-species comparison to the closely related but TE-poor *S. polyrhiza.* Despite major differences in their small RNA populations and DNA methylation patterns, *W. brasiliensis* does not exhibit expansion or broad transcriptional upregulation of canonical siRNA biogenesis or DNA methylation components relative to *S. polyrhiza*. This conservation of core silencing and methylation machinery enables a comparison in which the consequences of TE load and genome architecture can be evaluated with minimal confounding by pathway divergence. Although contributions from yet-unidentified regulators or genetic variation within known epigenetic components cannot be excluded, and although our analyses are based on a single accessions for both species, the high conservation of epigenetic regulators across duckweeds together with the substantial differences in genome size and TE load across the family suggests that the patterns described here are likely to recur in other accessions and additional duckweed species.

Within this comparative framework, the cross-species data resolve into two coherent layers. First, the per-element rules governing siRNA biogenesis and DNA methylation are conserved between *W. brasiliensis* and *S. polyrhiza*. Second, several genome-wide epigenetic features are markedly more pronounced or qualitatively distinct in W. brasiliensis: pervasive CG methylation across both TEs and genes, gene-proximity-dependent targeting of RdDM at intergenic TEs, near-exclusive 22-nt processing of PTGS substrates, and a quantitative correlation between gene body CG methylation and intragenic TE content. These divergences fall into two classes: those well-explained by the simple scaling of conserved silencing logic across the much larger TE complement of *W. brasiliensis*, and those that, in addition, point to *W. brasiliensis*-specific contributions whose mechanistic basis remains to be resolved.

Two prominent epigenetic features distinguishing *W. brasiliensis* from *S. polyrhiza* appear well-accounted for by the expanded TE content of *W. brasiliensis* acting on conserved per-element rules. The elevated abundance of 24-nt siRNAs in *W. brasiliensis* follows from the conserved preference of RdDM-associated pathways for long, intact TEs (Dombey *et al*., 2025): the extended and recent TE amplification documented here has produced a much larger pool of such elements in *W. brasiliensis*, particularly LTR retrotransposons. Similarly, the pervasive CG methylation observed across TEs and genes is consistent with a TE-load-driven scaling of maintenance methylation, in line with the broader angiosperm pattern where genome size, largely driven by TE accumulation, correlates positively with global CG methylation but not necessarily with non-CG methylation (Alonso *et al*., 2015; Niederhuth *et al*., 2016), and with models in which TE content promotes stable CG methylation as a basal repressive layer (Fedoroff, 2012). TEs in *W. brasiliensis* retain high CG methylation regardless of siRNA production, whereas in *S. polyrhiza* degenerated TE relics progressively lose methylation (Dombey *et al*., 2025). In both cases, the existing silencing machinery appears to be sufficient to handle this elevated TE load without global upregulation or expansion of small RNA biogenesis or DNA methylation factors.

By contrast, three further *W. brasiliensis*-specific features are not fully accounted for by the simple scaling discussed above. The first is the increase and near-exclusive 22-nt processing of PTGS substrates in *W. brasiliensis*, to which both increased TE-load and altered DCL processing likely contribute. Increased TE density in *W. brasiliensis* is expected to elevate the formation of inverted repeats (IRs) (SanMiguel *et al*., 1996; Slotkin *et al*., 2005; Chen *et al*., 2011; Gao *et al*., 2012; Arce *et al*., 2023; Trasser *et al*., 2024). We indeed identified IR-forming TEs as a minor but highly productive subset of 22-nt–biased loci (Group A) that resemble PTGS substrates. However, many Group A TEs lack detectable long IRs and shorter IR-containing TEs known to produce small RNAs, such as miniature inverted-repeat TEs (MITEs) (Li *et al*., 2011; Gagliardi *et al*., 2019; Arce *et al*., 2023; Guo *et al*., 2022; Martin *et al*., 2023), which are enriched in Group A, might have escaped our stringent analysis biased for long IRs. Additional triggers of PTGS-like processing, including miRNA targeting and translation-dependent silencing, may further contribute (Creasey *et al*., 2014; Kim *et al*., 2021; Oberlin *et al*., 2022). Beyond substrate availability, however, the near-exclusive 22-nt size at IRs and Group A TEs itself points to an altered DCL processing regime, distinct from that of *S. polyrhiza* where the same type of loci, and even transgenic hairpin precursor, produces a mix of 21- and 22-nt species despite the equivalent absence of DCL2 in both species (Dombey *et al*., 2025; Ernst *et al*., 2025).(Ernst *et al*., 2025)(Dombey *et al*., 2025)DCL4, which canonically processes such substrates into 21-nt siRNAs, might therefore display altered processing and/or modulated activity in duckweeds, and in *W. brasiliensis* in particular, producing both 21-22-nt siRNA in *S. polyrhiza* but only 22-nt siRNAs in *W. brasiliensis*. A non-exclusive alternative is that DCL1 contributes to 22-nt siRNA biogenesis from these substrates: in maize, DCL1 has been shown to produce 22-nt secondary siRNAs in absence of DCL4 (Petsch *et al*., 2015). The two tandemly duplicated DCL1 paralogues encoded by *W. brasiliensis* may provide a comparable source of 22-nt siRNAs at PTGS loci (Vaucheret and Voinnet, 2023). A 22-nt biased PTGS regime could partially compensate for the lack of DCL2 by retaining transitivity, the small-RNA amplification process for which 22-nt species are required (Vaucheret and Voinnet, 2023). We did not, however, detect transitivity at endogenous genes producing 22-nt siRNAs from their 3’UTRs. Moreover, the assignment of these siRNAs to PTGS may not be exclusive: in *A. thaliana*, Pol IV–derived 21–22-nt siRNAs can also direct RdDM (Panda *et al*., 2020), leaving open the possibility that a fraction of 22-nt siRNAs, particularly those co-produced with 24-nt species at Group B and C elements, contribute to DNA methylation rather than acting solely post-transcriptionally. The coincidence of TE-independent 22-nt siRNAs and high CG methylation at 3′ UTRs further suggests that 22-nt siRNAs and DNA methylation might be coupled. While the biogenesis and role of 22-nt siRNAs in duckweeds remains to be further investigated, these observations reveal previously underappreciated plasticity within conserved PTGS pathways.

A second feature is the gene-proximity-dependent targeting of RdDM at intergenic TEs in *W. brasiliensis*, a pattern not observed in *S. polyrhiza*. In *W. brasiliensis*, both 22- and 24-nt siRNA accumulation declines with distance from genes, and the strongest siRNA production is observed at long TEs located near or within genes. This gene-boundary-biased RdDM activity resembles patterns observed in other angiosperms, where RdDM reinforces chromatin boundaries at TE–gene interfaces rather than uniformly targeting bulk heterochromatin (Zemach *et al*., 2013; Gent *et al*., 2013; Stroud *et al*., 2014; Li *et al*., 2015; Sigman and Slotkin, 2016; Guo *et al*., 2021). In *S. polyrhiza*, however, the TE-length effect on siRNA production is conserved but no gene-proximity dependence is observed, indicating that gene-targeted RdDM is not a generic consequence of higher TE load but a feature specific to *W. brasiliensis*. At the molecular level, the relatively higher expression of the Pol V–dependent effector arm of RdDM (SUVH2, NRPE1, SPT5L, DRM2) in *W. brasiliensis* compared with *S. polyrhiza*, in contrast to the upstream 24-nt biogenesis arm (NRPD1, RDR2, DCL3) which is not, provides one candidate molecular correlate of this gene-proximity bias. A non-exclusive possibility is that the gene-proximity bias also reflects the chromatin accessibility of the underlying TEs to the RdDM machinery: at heterochromatinized gene-distal regions, Pol IV/V transcription of TEs may be more limited, restricting active RdDM to TEs in more accessible chromatin contexts. However, despite the gene proximity siRNA gradient, DNA methylation at intergenic TEs in *W. brasiliensis* is broadly uniform across distance deciles: only at the most gene-proximal TEs does non-CG methylation co-vary with siRNA production and TE length, a signature of RdDM-driven establishment, whereas at more distal positions methylation becomes homogeneous across the TE population, as expected of steady-state ZMET/CMT3–H3K9me2 maintenance. Selective RdDM activity at gene-proximal TEs may therefore primarily contribute to establishing and reinforcing non-CG methylation at TE-gene boundaries, while bulk methylation across the rest of the genome is likely sustained by siRNA-independent DNA methylation maintenance pathways. Even at regions far away from genes, individual TEs that produce siRNAs still gain additional non-CG methylation, consistent with siRNA-mediated reinforcement operating on top of this maintenance baseline.

The third and most pronounced *W. brasiliensis*-specific feature is the strong positive correlation between gene body CG methylation (gbM) and intragenic TE content, a relationship not detectable in *S. polyrhiza*. The intragenic constraint on non-CG methylation despite siRNA production at intronic TEs is itself shared between the two species: intronic TEs are typically heterochromatinized in plants to limit interference with splicing and other transcriptional processes (Liu *et al*., 2004; Chen *et al*., 2011; West *et al*., 2014; Le *et al*., 2015; Espinas *et al*., 2020; Zhou *et al*., 2020). Our data extend this conserved feature to both duckweeds, consistent with mechanisms that restrict heterochromatin spreading within genes (Saze *et al*., 2008). The gbM correlation itself, however, appears to be a *W. brasiliensis*-specific outcome. In most plant systems, gbM has been interpreted as largely independent of TEs (Zilberman *et al*., 2006; Takuno and Gaut, 2012; Bewick *et al*., 2016; Picard and Gehring, 2017). Recent models propose that transient heterochromatic incursions introduce DNA methylation within genes, with active removal of H3K9me2 preventing stable non-CG accumulation while CG methylation persists through its efficient maintenance (Wendte *et al*., 2019; Zhang *et al*., 2020; Papareddy *et al*., 2021; Muyle *et al*., 2022; Zhang *et al*., 2024). Our observations in *W. brasiliensis* are consistent with this framework: intragenic TE insertions may generate localized heterochromatic pressure within otherwise euchromatic genes, with CG methylation persisting beyond the TE sequence into adjacent gene body regions. Crucially, the absence of an equivalent gbM gradient in *S. polyrhiza*, including genes that reach high TE-content categories, indicates that gene-level TE content alone is not sufficient to produce this regime; additional *W. brasiliensis*-specific contributions must therefore be involved. Three non-exclusive possibilities can be considered. First, the substantially higher intronic TE content of *W. brasiliensis,* approximately an order of magnitude more TE-derived intronic sequence than in *S. polyrhiza,* would be expected to have produced more frequent heterochromatic incursions per gene than in *S. polyrhiza*. Second, the elevated CG-maintenance baseline observed at TEs in *W. brasiliensis* would favor the persistence of mCG once it is introduced into gene bodies, in line with the model in which CG methylation persists by maintenance after the transient heterochromatic source has been cleared. Third, the body plan simplification and predominantly clonal life history of *W. brasiliensis* may itself stabilize emergent methylation states: As one of the most morphologically reduced angiosperms, *W. brasiliensis* is characterized by a highly simplified body plan and continuous formation of meristems. Although direct links between morphological reduction and epigenetic organization remain unexplored, such developmental minimalism may further constrain how silencing pathways are deployed across tissues. Vegetative propagation can transmit somatic epigenetic states across generations (Wibowo *et al*., 2018; Wibowo *et al*., 2022), potentially stabilizing methylation landscapes that might otherwise be removed during chromatin reorganization and non-CG methylation reprogramming accompanying flowering, gametogenesis, and embryogenesis (Peng and Karpen, 2008; Ingouff *et al*., 2010; Jullien *et al*., 2012; She and Baroux, 2015; Bouyer *et al*., 2017; Kawakatsu *et al*., 2017; Papareddy *et al*., 2020; Gutzat *et al*., 2020; Papareddy *et al*., 2021; Borg *et al*., 2021; Parent *et al*., 2021; Hemenway and Gehring, 2023). The rarity of sexual reproduction may reduce opportunities to prevent non-CG methylation from stabilizing and spreading within TE-containing genes, favoring mechanisms that restrict non-CG methylation in intragenic contexts. As other duckweeds with increased TE content also display increased global gbM (Ernst *et al*., 2025), disentangling the relative contributions of these factors will require comparative analyses across additional duckweed species, particularly those with intermediate TE loads and varying reproductive modes.

Beyond the cross-species divergences discussed above, the co-occurrence of multiple siRNA sizes at individual TEs in both duckweeds points to additional layers of TE regulatory complexity. Such overlap may reflect spatial or temporal partitioning of PTGS and RdDM activities across tissues (Pachamuthu and Borges, 2023). A non-exclusive possibility is that all size classes arise from a shared pool of precursor transcripts processed by different DCLs rather than from dedicated pathways. In *A. thaliana*, Pol IV transcripts that feed the 24-nt pathway also yield 21–22-nt siRNAs, and, when Pol IV is absent, Pol II–transcribed TEs give rise to both 21–22-nt and 24-nt siRNAs (Panda *et al*., 2020). The strong positional overlap of 22- and 24-nt siRNAs in Group B TEs, and the residual 22-nt signal over the 24-nt–producing regions of Group C TEs, is consistent with such shared processing, whether from Pol IV- or Pol II–derived precursors. In addition, the truncated DCL3 protein in *W. brasiliensis* may permit mixed 24/22-nt outputs from siRNA precursors due to reduced processing precision (Liu *et al*., 2012). Similar DCL3 truncations have been reported in other angiosperms (Šečić *et al*., 2019; Cervantes-Pérez *et al*., 2021), suggesting potential flexibility in DCL3 function. Although functional validation will be required, these possibilities, together with the candidate DCL1 contribution to 22-nt siRNA biogenesis discussed above, underscore the plasticity of small RNA biogenesis pathways in duckweeds and in angiosperms in general.

Taken together, our analyses of *W. brasiliensis* and its cross-species comparison with *S. polyrhiza* reveal a duckweed silencing system in which conserved per-element rules, shared between the two species, produce divergent genome-wide outcomes when applied to dramatically different TE loads and genome architectures. While some of these outcomes follow naturally from the scaling of this conserved logic across the much larger TE complement of *W. brasiliensis*, others require additional *W. brasiliensis*-specific contributions that remain to be characterized. More broadly, these findings highlight the central role of TEs in shaping plant epigenomes and establish duckweeds as a uniquely informative comparative system for dissecting the evolutionary interplay between TE accumulation and host silencing mechanisms.

## Materials and Methods

### Plant material and growth conditions

*Wolffia brasiliensis* Wess. accessions were obtained from the Landolt Collection (CNR-IBBA-MIDW, Milan, Italy) (Morello *et al*., 2024). Axenic cultures were established as previously described (Barragán-Borrero *et al*., 2026) immersed for 2 min. in 1% Danklorix and maintained on N medium under long-day conditions (16 h light/8 h dark) at 21°C. Biological replicates represent independently propagated cultures.

### Flow cytometry

Genome size was estimated by flow cytometry using propidium iodide-stained nuclei as in (Temsch *et al*., 2010; Barragán-Borrero *et al*., 2026) co-processed with internal standards. Three independent biological replicates were analyzed per accession.

### Genome sequencing, assembly and annotation

High-molecular-weight DNA was isolated from purified nuclei as in (Lutz *et al*., 2011; Rabanal *et al*., 2022) and sequenced using PacBio HiFi technology. HiFi reads were assembled with Hifiasm (v0.16.1) (Cheng *et al*., 2021). Chloroplast-derived contigs and redundant overlapping contigs were removed prior to generating the final assembly. Gene models were predicted using Funannotate (Palmer and Stajich, 2020) integrating PacBio Iso-Seq and Illumina RNA-seq evidence, with *ab initio* prediction supported by BUSCO (Simão *et al*., 2015) (embryophyta dataset). Protein domains were annotated using InterProScan (Paysan-Lafosse *et al*., 2022), and gene models were curated based on transcript support.

### Transposable element annotation and divergence analysis

TEs were annotated using a combined *de novo* and homology-based approach integrating EDTA and RepeatModeler2, followed by genome masking with RepeatMasker (Smit *et al*., n.d.; Ou *et al*., 2019; Flynn *et al*., 2020). TE families were curated and classified based on structural characteristics and conserved protein domains using MCHelper (Orozco-Arias *et al*., 2024). Sequence divergence relative to TE consensus sequences was estimated using Kimura substitution levels in RepeatMasker alignments. Elements were classified as intragenic or intergenic based on overlap with annotated genes.

### Gene and TE overlap analysis

Classification of TEs based on the overlap with genes was performed using ParasiTE (Berthelier *et al*., 2023), using the Iso-Seq reads as input. In brief, fully exonic TEs were those with a >80% overlap with exons, as Intronic if <1% of the TE length overlapped with an exon and partially exonic those TEs within 1%-80% overlap with exonic sequences. Intragenic TEs that were not overlapping with a predicted gene model were filtered out as Intergenic.

### Transcriptome profiling

Total RNA was extracted from ∼100 mg tissue and Illumina RNA-seq libraries were prepared and sequenced (150 bp paired-end). Reads were quality-filtered with fastp (v0.20.1) (Chen *et al*., 2018) and quantified against the *W. brasiliensis* genome using pseudo-alignment with Kallisto (Bray *et al*., 2016). Transcript abundance values (TPM) were averaged across three independent biological replicates. Iso-Seq long-read transcript data were processed independently and aligned to the genome using minimap2 (v2.17) (Li, 2021) to support gene model refinement. For *A. thaliana* and *S. polyrhiza,* publicly available datasets were used (GSM6892967, GSM6892968, GSM6892969, SRX26200034, SRX26200035, SRX26200036) (Li *et al*., 2023; Dombey *et al*., 2025).

### Small RNA sequencing

Small RNAs were isolated either from total RNA or from TraPR as described previously (Grentzinger *et al*., 2020; Dombey *et al*., 2025). Libraries were generated using sequential 3′ and 5′ adapter ligation with randomized terminal nucleotides to minimize ligation bias, followed by reverse transcription and PCR amplification. Two independent biological replicates were sequenced per accession unless otherwise stated. Adapter-trimmed reads were aligned to the *W. brasiliensis* genome using Bowtie 2 (v1.2.2) (Langmead and Salzberg, 2012), allowing at most one mismatch and retaining uniquely mapping reads unless specified. Size-specific read counts were extracted from mapped reads. Genome-wide small RNA-producing loci were identified using ShortStack (v3.8.5) (Axtell, 2013) with a minimum coverage threshold of 0.6 RPM. Coverage tracks were normalized to counts per million (CPM). For *hpScarlet*, processed reads were aligned to the hairpin sequence with zero mismatches and normalized to CPM.

### Small RNA locus classification

For each ShortStack-defined locus, the ratio of 22-nt to 24-nt reads was calculated using uniquely mapped reads. To define objective classification thresholds, the empirical distribution of locus-specific 22-nt:24-nt ratios was modeled using kernel density estimation in R (v4.2.2; https://www.r-project.org). Local minima in the density curve were identified computationally, and the minimum closest to unity (ratio = 1) was selected as the primary separation point. Symmetric thresholds were defined as LT and 1/LT, generating three classes: 22-nt enriched, balanced, and 24-nt enriched loci. Classification was applied uniformly across biological replicates.

### DNA methylation profiling

Genomic DNA was extracted and enzymatic methyl-seq (EM-seq) libraries were prepared and sequenced (150 bp paired-end). Three independent biological replicates were generated. Reads were adapter-trimmed and aligned to the *W. brasiliensis* genome using Bismark (Krueger and Andrews, 2011) in non-directional mode. PCR duplicates were removed prior to methylation calling. Weighted methylation levels were calculated per cytosine as the proportion of methylated reads over total coverage. Conversion efficiency was >98.6% in all replicates, monitored using chloroplast sequences. Cytosines with coverage below four reads were excluded. For genome-wide and feature-level analyses, biological replicates were first analyzed independently and subsequently merged after confirming high concordance. Global methylation levels and feature-specific methylation profiles were computed using deepTools (Ramírez *et al*., 2016). Only features containing at least four informative cytosines were retained for downstream analysis.

### Phylogenetic and domain analysis

Putative orthologues of RNA silencing components were identified by homology searches using BLAST+ (Camacho *et al*., 2009) against the *W. brasiliensis* proteome and genome using *Arabidopsis* and *Spirodela* queries. Candidate genes were validated based on conserved domain architecture, and phylogenetic trees built in CLC Main Workbench (v20).

### Agrobacterium-mediated transformation

Transient transformation was performed using *Agrobacterium tumefaciens* strain EHA105 via vacuum infiltration following the *Spirodela* protocol in (Barragán-Borrero *et al*., 2026) with minimal modifications. Plasmids used: *pZmUbq:eGFP* (EPR <u>#869</u>) (Barragán-Borrero *et al*., 2026); *pGGSun-pUBIPars-hpScarlet* (Incarbone *et al*., 2023). Fluorescence images were obtained in an Azure biomolecular scanner and quantified using Fiji (v2.16.0) (Schindelin *et al*., 2012). Plants were scored as positive when signal exceeded control means by more than two standard deviations.

### Statistical analysis

All statistical analyses were performed in R (v4.2.2). Statistical tests are specified in the corresponding figure legends.

Fully detailed Materials and Methods can be found in the associated Supplemental Information.

## Acknowledgements

We thank current and former members of the Marí-Ordóñez group, in particular Rodolphe Dombey for assistance with computational analysis and colleagues at the Gregor Mendel Institute and the Vienna BioCenter (VBC) for valuable discussions and feedback. We are grateful to Marco Incarbone (Max Planck Institute of Plant Physiology, Golm) for providing the hpScarlet plasmid, Hanna Weiss-Schneeweiss and Eva Temsch from the University of Vienna for training and advice on flow cytometry and chromosome spreads. We also thank the Vienna BioCenter Core Facilities (VBCF): NGS for PacBio, EM-seq, RNA-seq, and small RNA library preparation and sequencing; BioOptics for microscopy and flow cytometry support; and the IMP Molecular Biology Service for reagents.

## Author contributions

A.M.-O. and D.B.-A. conceived and designed the study. D.B.-A. performed most experiments with assistance from V.B.-B. M.A., supervised by D.B.-A., optimized agroinfiltration-mediated transient expression in *Wolffia*. P.L.-R., supervised by D.B.-A., assisted with the cross-species small RNA screen. D.B.-A. conducted computational and statistical analyses. L.M. performed the TBP analysis. A.M.-O. and D.B.-A. analyzed the data, assembled the figures, and wrote the manuscript.

## Data availability

