## Supplemental Information for "Transposon expansion is associated with reorganization of small RNA and DNA methylation landscapes in the morphologically minimal angiosperm *Wolffia brasiliensis*"

**Figure S1**: Genome size estimations of *W. brasiliensis* and *W. arrhizal* accessions by flow cytometry.

**Figure S2:** *Wolffia* morphology and vegetative growth

**Figure S3:** *W. brasiliensis* genome assembly and annotation.

**Figure S4:** Evaluation of TE relative age based on length asn sequence divergence.

**Figure S5:** Profiling of total and miRNA-mapping siRNAs in *W. brasiliensis.*.

**Figure S6:** List of gene IDs.

**Figure S7:** Pairwise comparison of DCL proteins identified in *W. brasiliensis.*

**Figure S8:** Genome duplication of DCL1 in *W. brasiliensis*.

**Figure S9:** Phylogenetic analysis of DCL proteins in *W. brasiliensis*.

**Figure S10:** Structural analysis of *WbDCL3.*

**Figure S11:** Gene expression of silencing components in *W.brasiliensis*.

**Figure S12:** Profile of siRNAs from siRNA producing TEs in *W. brasiliensis*.

**Figure S13:** Analysis of TE-derived small RNAs in *S. polyrhiza*.

**Figure S14:** TE Family siRNA enrichment analysis in *W. brasiliensis.*

**Figure S15:** TAS3 loci identified in *W. brasiliensis*

**Figure S16:** Agrobacterium-mediated Transient expression on *W. brasiliensis* fronds.

**Figure S17:** Analysis of non-templated nucleotides in uniquely mapped siRNA in Wolffia.

**Figure S18:** Example of an Inverted Repeat in Wolffia.

**Figure S19:** Examples of additional siRNA-producing IR+ clusters in Wolffia.

**Figure S20:** Identification and characterization of endogenous IRs in Spirodela.

**Figure S21:** Correlation between 22nt-siRNA and expression on TEs.

**Figure S22:** Distribution of DNA methylation in siRNA TE-groups in Spirodela.

**Figure S23:** Effect of TE size on DNA methylation.

**Figure S24:** Impact of TE size on siRNA production in Spirodela.

**Figure S25:** TE siRNA production correlates with gene distance and TE size.

**Figure S26:** Analysis of DNA methylation by distance to genes and TE size in Wolffia and Spirodela.

**Figure S27:** Intragenic TE-distribution in Spirodela and Wolffia.

**Figure S28:** TE-genes and TE coverage distribution across gene annotations.

**Figure S29:** Examples of siRNA-producing intergenic and intronic regions in Wolffia.

**Figure S30:** TE siRNAs and DNA methylation by genomic compartment in Wolffia and Spirodela.

**Figure S31:** Examples of siRNA producing 3'UTRs in Wolffia.

**Figure S32:** DNA methylation levels on gene annotations correlate with gene TE coverage in Wolffia.

**Figure S33:** Examples of gene body methylation on genes containing or not TEs in in Wolffia.

**Figure S34:** DNA Methylation analysis on CDS sequences in *W. brasiliensis*.

**Figure S35:** DNA Methylation analysis on CDS sequences in *S. polyrhiza*.

**Extended Materials and Methods**

### FIGURE S1

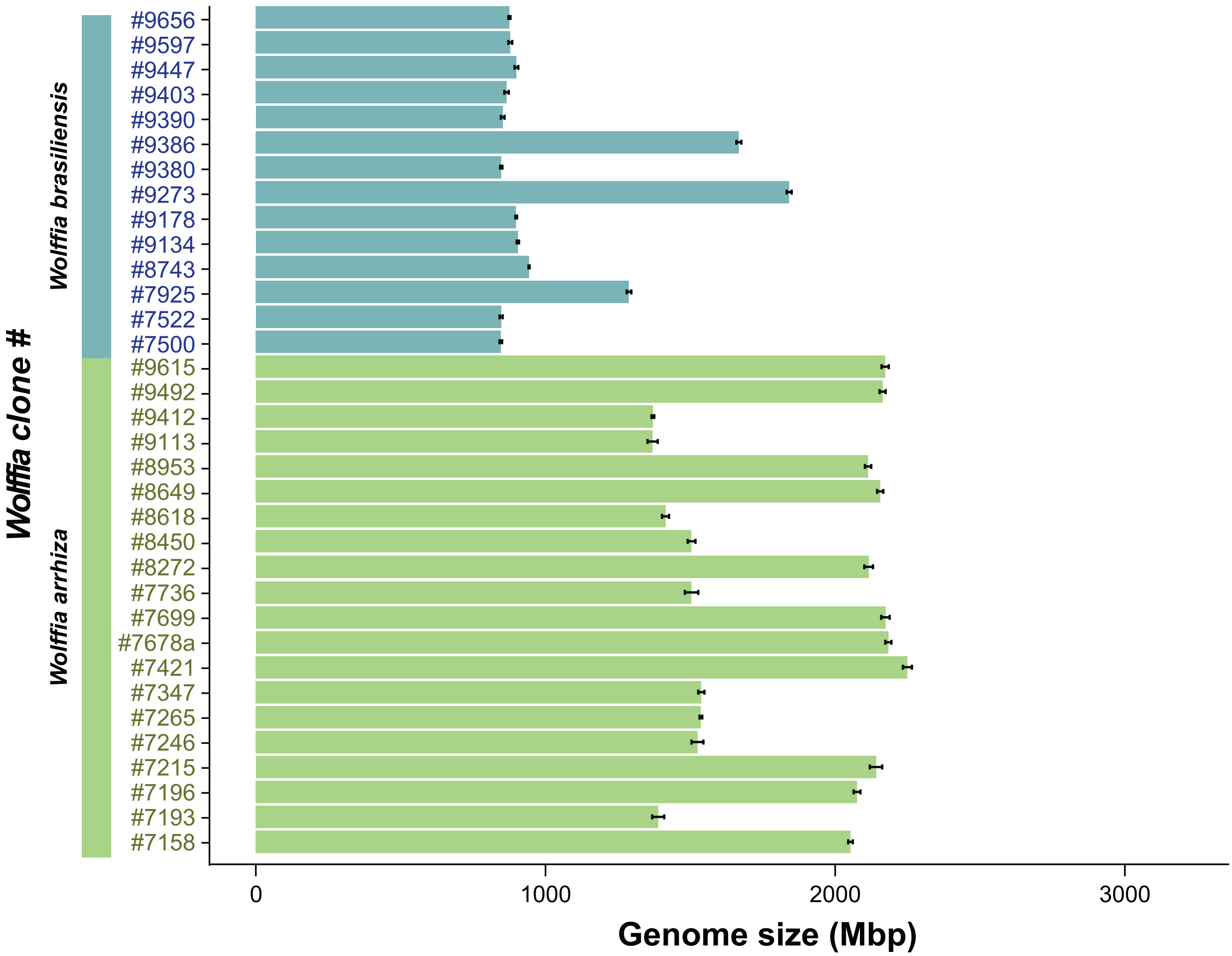


**Figure S1. Genome size estimation of *W. brasiliensis* and *W. arrhiza* accessions by flow cytometry**. Bar length represent the mean of 3 replicates, and error bars the SD from the mean.

### SUPPLEMENTAL FIGURE S2

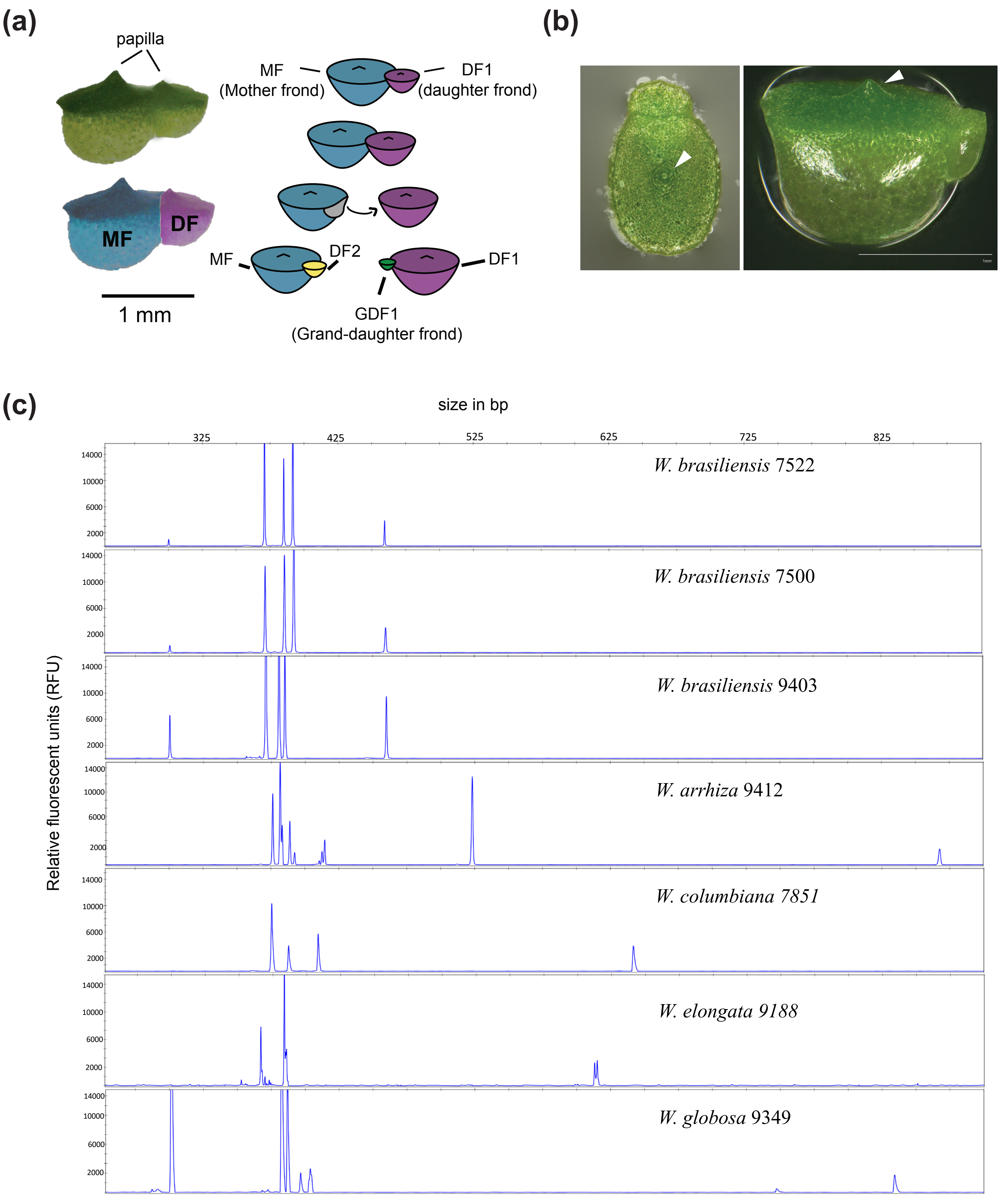


**Figure S2. Morphology and vegetative growth in *Wolffia.* (a)** Image and schematic representation of *W. brasiliensis* #7522 fronds and asexual propagation during vegetative growth. MF: Mother frond, DF: Daughter frond. **(b)** Representative images of *W. brasiliensis* fronds, white arrowhead points to the papilla**. c)** Electropherograms of Tubullin-Based-Polymorphism (TBP) analysis of several *Wolffia* species based on 1st intron-length polymorphism.

### SUPPLEMENTAL FIGURE S3

(a)

| *W. brasiliensis* (#7522) | |
|---|---|
| Genome Size (FC) | 846,88 Mbp |
| Sequencing platform | PacBio HiFi |
| Assembly size | 771 Mbp |
| Contigs | 250 |
| L90 | 88 |
| N50 | 9.23 Mbp |
| BUSCO score | 91.4 % |
| % Repeats | 74,52 % |
| No of Genes | 25161 |

(b)

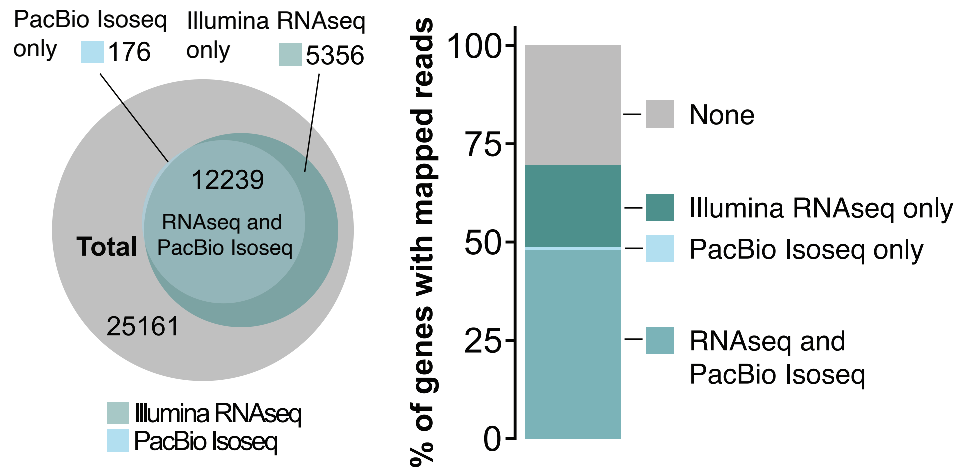


(c)

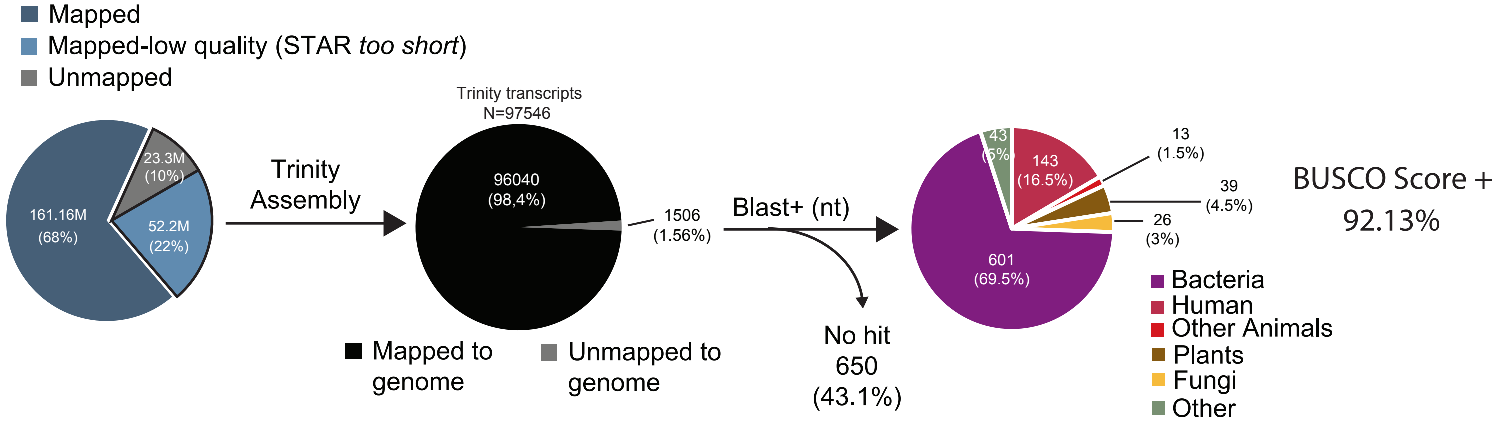


(d)

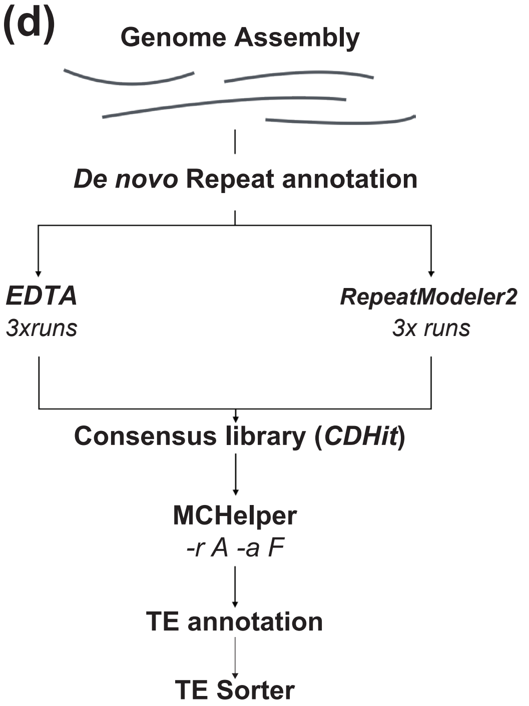


(e)

| | Class | Order | Superfamily/code | Elements | bp masked | % of repeats | % of genome |
|---|---|---|---|---|---|---|---|
| TRANSPOSABLE ELEMENTS (TEs) | *Class I: Retrotransposons ("copy-paste")* (R) | *Unknown* (X) | *Unknown* (X) RXX | 4256 | 1263859 | 0,22 | 0,16 |
| | | Long Terminal Repeat (LTR) (L) | *Ty1/Copia* (C) RLC | 140075 | 105839482 | 18,42 | 13,73 |
| | | | *Ty3/Gypsy* (G) RLG | 115987 | 118545063 | 20,63 | 15,37 |
| | | | *Unknown* (X) RLX | 447633 | 185906516 | 32,35 | 24,11 |
| | | | *LARDs* (L) RLL | 25078 | 17297083 | 3,01 | 2,24 |
| | | | *TRIMs* (T) RLT | 13622 | 11988895 | 2,09 | 1,55 |
| | | non-LTR LINE (I) | *LINE/Unknown* (L/X) RIL(X) | 6176 | 2085046 | 0,36 | 0,27 |
| | *Class II: DNA transposons (cut-paste)* (D) | Terminal Inverted Repeat (TIR) (T) | MITE | 87919 | 21685221 | 3,77 | 2,81 |
| | | | *Mutator* (M) DTM | 2955 | 2180045 | 0,38 | 0,28 |
| | | | *Unknown* (X) DTX | 164271 | 53325470 | 9,28 | 6,92 |
| | | | *hAT* (A) DTA | 7256 | 10434449 | 1,82 | 1,35 |
| | | | *Merlin* (E) DTE | 3 | 1666 | 0,00 | 0,00 |
| | | non-TIR (H) | *Unknown* (X) DHX | 1788 | 823994 | 0,14 | 0,11 |
| OTHER REPEATS | | | Unknown | 62 | 103119 | 0,02 | 0,01 |
| | | | Satellite | 1973 | 486103 | 0,08 | 0,06 |
| | | | Telomeric | 2209 | 623383 | 0,11 | 0,08 |
| | | | Simple repeat | 27299 | 22242058 | 3,87 | 2,88 |
| | | | Pararetrovirus | 7 | 1420 | 0,00 | 0,00 |
| | | | Organellar | 83418 | 19762632 | 3,44 | 2,56 |
| | | | **Total:** | 1131987 | 574595504 | 100 | **74,52** |

**Figure S3. *W. brasiliensis* genome assembly and annotation. (a)** Table showing the main genome assembly statistics. FC: Flow cytometry. **(b)** Number and proportion of genes with associated transcripts. Left: Venn Diagram showing the amount of gene annotations covered by PacBio Isoseq reads, Illumina RNA seq reads or both. Right: Stacked barplot showing the % of genes with mapping reads from each or both platforms. **(c)** De novo transcriptome assembly from RNA-seq reads with low quality mapping (STAR "too short") and unmapped to the genome. 98.4% of assembled transcripts remapped to the assembly; of the remaining 1.56%, 39 transcripts returned plant hits by BLAST. Adding these to the BUSCO analysis raised the recovered fraction from 91.4% to 92.1% (See Table S1). **(d)** Schematic representation of the strategy followed to annotate Transposable Elements (TEs) in (b) **(e)** Transposon classification and nomenclature used in this study, alongside with the number of elements and genome occupancy of each TE superfamily.

### SUPPLEMENTAL FIGURE S4

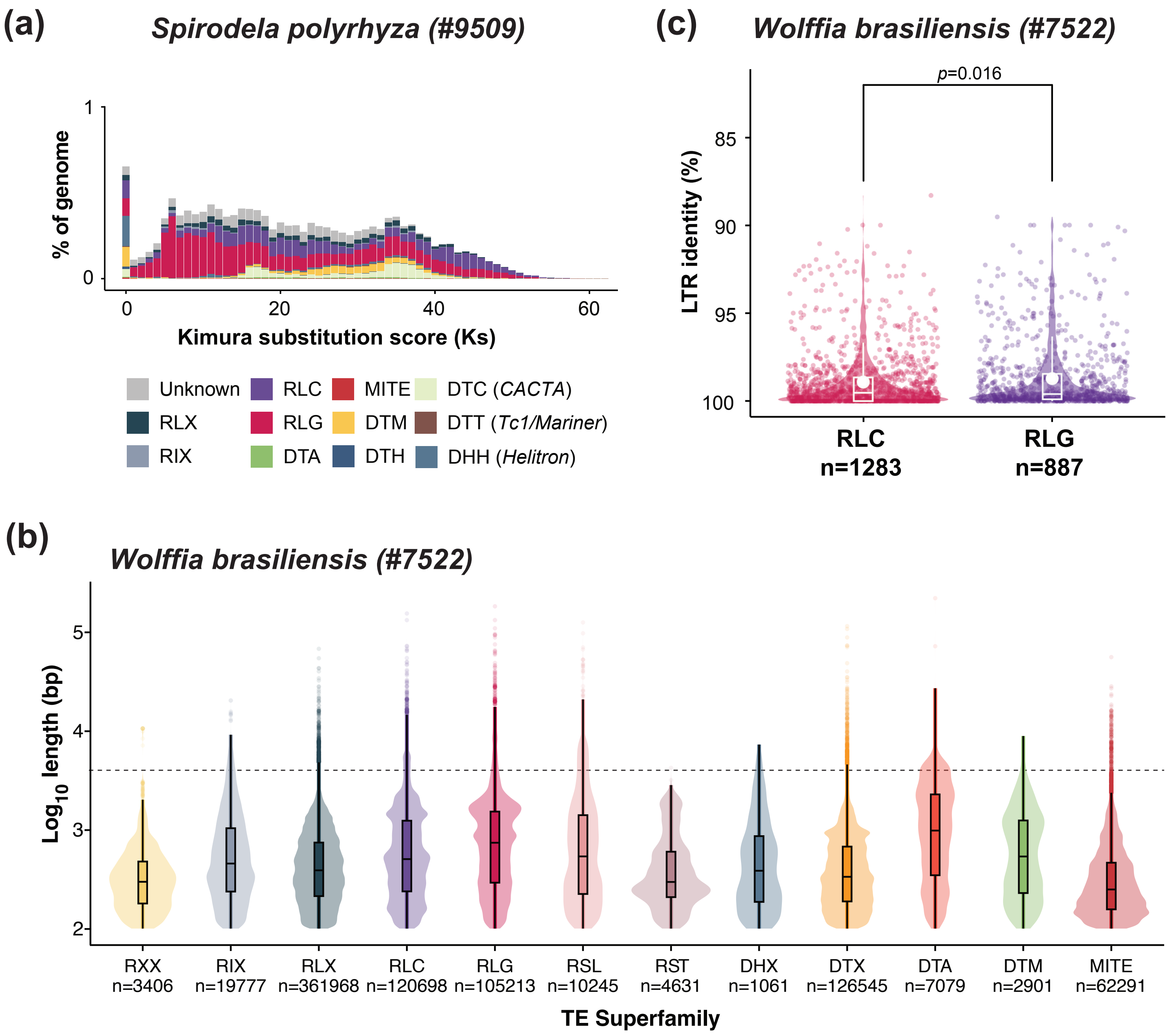


**Figure S4. Evaluation of TE relative age based on length and sequence divergence. (a)** Kimura plot showing the divergence distribution of single TE copies from their consensus sequence in *S. polyrhiza* #9509 and their genome occupancy. **(b)** Violin and boxplot analysis of the length distribution of TEs belonging to different TE superfamilies. Reference length of 4Kbp is represented as an horizontal dashed line. **(c)** LTR identity violin and boxplot (%) of "intact" TEs belonging to RLC or RLG TE superfamilies identified by EDTA in *W. brasiliensis.* In all boxplots the median is represented as a solid bar, with box upper and bottom limits representing the first and third quartiles. Whiskers range is 1.5 times the interquartile range. The white dot represents the distribution mean.

### SUPPLEMENTAL FIGURE S5

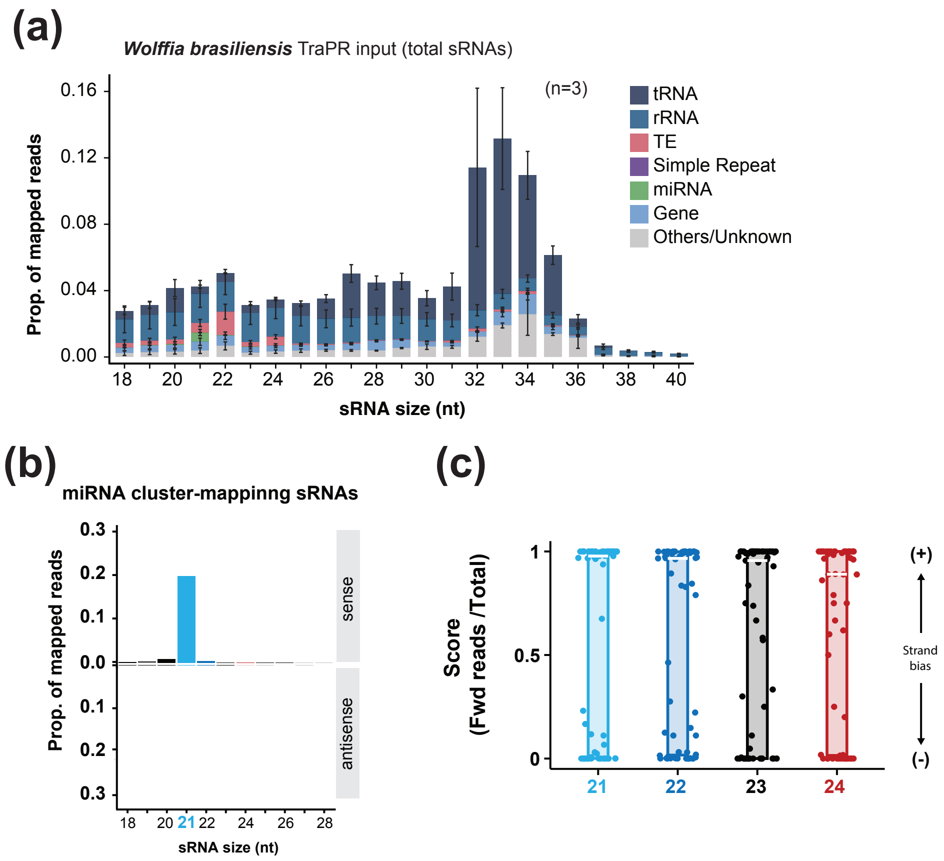


**Figure S5. Profiling of total and miRNA-mapping siRNAs in *W. brasiliensis*. (a)** Size distribution and genomic mapping of total RNA extracted small RNAs. Bar heights represent the proportion of the total mapped reads for each size, and the genomic features they were found mapping to. Bar heights represent the average of three replicates, and the error bars the SD from the average. **(b)** Proportion of miRNA-mapping RISC-loaded siRNAs distributed by size and relative mapping orientation to the miRNA loci. **(c)** Strand bias distribution of miRNAs. A strand-bias score (0-1) for each miRNA locus shown as the ratio of forward strand (+) mapping reads to total mapped reads independently of the orientation of the TE annotation. A score of 0 indicates exclusive mapping to the reverse strand (-), while a score of 1 indicates exclusive mapping to the forward (+) strand. **(d)** Stacked barplot showing the 5' nucleotide bias of miRNA-derived 21-24nt siRNAs.

### SUPPLEMENTAL FIGURE S6

**miRNA biogenesis**

| Gene | Arabidopsis | AGI | Spirodela | Spirodela Gene ID | Wolffia | Wolffia Gene ID |
|---|---|---|---|---|---|---|
| SE | AtSE | AT2G27100 | SpSE | Sp9509d012g007660 | WbSE | Wb7522d003g006430 |
| HYL1 | AtHYL1 | AT1G09700 | SpHYL1 | Sp9509d001g003860 | WbHYL1 | Wb7522d040g001880 |
| DRB2a | AtDRB2 | AT2G28380 | SpDRB2a | Sp9509d002g000320 | WbDRB2a | Wb7522d099g000820 |
| DRB2b | | | SpDRB2b | Sp9509d014g000150 | WbDRB2b | Wb7522d124g000210 |
| HEN1 | AtHEN1 | AT4G20910 | SpHEN1 | Sp9509d017g000460 | WbHEN1 | Wb7522d082g000650 |
| DCL1a | AtDCL1 | AT1G01040 | SpDCL1 | Sp9509d002g008800 | WbDCL1a | Wb7522d041g000520 |
| DCL1b | | | | | WbDCL1b | Wb7522d041g000580 |

**21–22-nt siRNA biogenesis**

| Gene | Arabidopsis | AGI | Spirodela | Spirodela Gene ID | Wolffia | Wolffia Gene ID |
|---|---|---|---|---|---|---|
| SGS3 | AtSGS3 | AT5G23570 | SpSGS3 | Sp9509d004g006000 | WbSGS3 | Wb7522d013g004650 |
| RDR1a | AtRDR1 | AT1G14790 | SpRDR1a | Sp9509d018g002090 | WbRDR1a | Wb7522d011g002820 |
| RDR1b | | | SpRDR1b | Sp9509d018g002070 | WbRDR1b | Wb7522d011g002900 |
| RDR6 | AtRDR6 | AT3G49500 | SpRDR6 | Sp9509d006g004830 | WbRDR6 | Wb7522d042g001320 |
| RDR3 | AtRDR3 | AT2G19910 | | | | |
| RDR4 | AtRDR4 | AT2G19920 | | | | |
| RDR5 | AtRDR5 | AT2G19930 | | | | |
| DRB4 | AtDRB4 | AT3G62800 | SpDRB4 | Sp9509d001g004790 | WbDRB4 | Wb7522d070g000480 |
| DRB7.1 | AtDRB7.1 | AT1G80650 | | | | |
| DRB7.2 | AtDRB7.2 | AT4G00420 | | | | |
| DCL4 | AtDCL4 | AT5G20320 | SpDCL4 | Sp9509d005g008560 | WbDCL4 | Wb7522d081g000400 |
| DCL2 | AtDCL2 | AT3G03300 | | | | |
| RTL1 | AtRTL1 | AT4G15417 | | | | |

**Argonautes**

| Gene | Arabidopsis | AGI | Spirodela | Spirodela Gene ID | Wolffia | Wolffia Gene ID |
|---|---|---|---|---|---|---|
| AGO1 | AtAGO1 | AT1G48410 | SpAGO1 | Sp9509d020g001530 | WbAGO1 | Wb7522d017g002510 |
| AGO10 | AtAGO10 | AT5G43810 | SpAGO10 | Sp9509d008g011640 | WbAGO10 | Wb7522d008g005230 |
| AGO5a | AtAGO5 | AT2G27880 | SpAGO5a | Sp9509d007g002750 | WbAGO5 | Wb7522d012g004110 |
| AGO5b | | | SpAGO5b | Sp9509d007g002760 | | |
| AGO5c | | | SpAGO5c | Sp9509d007g002770 | | |
| AGO5d | | | SpAGO5d | Sp9509d007g002790 | | |
| AGO5e | | | SpAGO5e | Sp9509d007g002800 | | |
| AGO5f | | | SpAGO5f | Sp9509d007g002810 | | |
| AGO5g | | | SpAGO5g | Sp9509d007g002820 | | |
| AGO5h | | | SpAGO5h | Sp9509d007g002830 | | |
| AGO2 | AtAGO2 | AT1G31280 | | | | |
| AGO3 | AtAGO3 | AT1G31290 | | | | |
| AGO7 | AtAGO7 | AT1G69440 | SpAGO7 | Sp9509d009g009840 | WbAGO7 | Wb7522d014g001860 |
| AGO4a | AtAGO4 | AT2G27040 | SpAGO4a | Sp9509d001g011250 | WbAGO4 | Wb7522d036g001130 |
| AGO4b | | | SpAGO4b | Sp9509d006g000640 | | |
| AGO8 | AtAGO8 | AT5G21030 | | | | |
| AGO9 | AtAGO9 | AT5G21150 | | | | |
| AGO6 | AtAGO6 | AT2G32290 | | | | |

**24-nt siRNA / RdDM**

| Gene | Arabidopsis | AGI | Spirodela | Spirodela Gene ID | Wolffia | Wolffia Gene ID |
|---|---|---|---|---|---|---|
| SHH1 | AtSHH1 | AT1G15215 | | | | |
| CLSY1 | AtCLSY1 | AT3G42670 | | | | |
| CLSY2 | AtCLSY2 | AT5G20420 | | | | |
| CLSY3 | AtCLSY3 | AT1G05490 | SpCLSY3 | Sp9509d011g003360 | WbCLSY3 | Wb7522d058g001390 |
| CLSY4 | AtCLSY4 | AT3G24340 | | | | |
| NRPD1 | AtNRPD1 | AT1G63020 | SpNRPD1 | Sp9509d0013g000080 | WbNRPD1 | Wb7522d059g000360 |
| RDR2 | AtRDR2 | AT4G11130 | SpRDR2 | Sp9509d002g008450 | WbRDR2 | Wb7522d119g000030 |
| DCL3 | AtDCL3 | AT3G43920 | SpDCL3 | Sp9509d005g001240 | WbDCL3 | Wb7522d021g002440 |
| SUVH2 | AtSUVH2 | AT2G33290 | SpSUVH2 | Sp9509d020g001160 | WbSUVH2 | Wb7522d034g002220 |
| SUVH9 | AtSUVH9 | AT4G13460 | | | | |
| DRD1 | AtDRD1 | AT2G16390 | SpDRD1 | Sp9509d007g000370 | WbDRD1 | Wb7522d012g000500 |
| NRPE1 | AtNRPE1 | AT2G40030 | SpNRPE1 | Sp9509d003g010520 | WbNRPE1 | Wb7522d009g001390 |
| SPT5L | AtSPT5L | AT5G04290 | SpSPT5L | Sp9509d001g003550 | WbSPT5L | Wb7522d118g000060+070 |
| DRM2 | AtDRM2 | AT2G33830 | SpDRM2 | Sp9509d008g003000 | WbDRM2 | Wb7522d102g000510 |
| DRM1 | AtDRM1 | AT1G28330 | | | | |
| DRM3 | AtDRM3 | AT3G17310 | | | | |

**DNA methylation maintenance**

| Gene | Arabidopsis | AGI | Spirodela | Spirodela Gene ID | Wolffia | Wolffia Gene ID |
|---|---|---|---|---|---|---|
| DDM1 | AtDDM1 | AT5G66750 | SpDDM1 | Sp9509d003g002770 | WbDDM1 | Wb7522d006g002490 |
| MET1 | AtMET1 | AT5G49160 | SpMET1 | Sp9509d003g004910 | WbMET1 | Wb7522d006g004820 |
| BRM | | | | | WbBRM | Wb7522d012g000360 |
| VIM1 | AtVIM1 | AT1G57820 | SpVIM1 | Sp9509d004g007270 | WbVIM1 | Wb7522d013g002530 |
| VIM2 | AtVIM2 | AT1G66050 | | | | |
| VIM3 | AtVIM3 | AT5G39550 | | | | |
| VIM4 | AtVIM4 | AT1G66040 | | | | |
| VIM5 | AtVIM5 | AT1G57800 | | | | |
| CMT3 | AtCMT3 | AT1G69770 | SpZMET | Sp9509d003g011020 | WbZMET | Wb7522d009g001280 |
| CMT2 | AtCMT2 | AT4G19020 | | | | |
| SUVH4 | AtSUVH4 | AT5G13960 | SpSUVH4 | Sp9509d004g011450 | WbSUVH4 | Wb7522d016g001290 |
| SUVH5 | AtSUVH5 | AT2G35160 | SpSUVH5 | Sp9509d008g010000 | WbSUVH5 | Wb7522d008g003350 |
| SUVH6 | AtSUVH6 | AT2G22740 | | | | |

**Other**

| Gene | Arabidopsis | AGI | Spirodela | Spirodela Gene ID | Wolffia | Wolffia Gene ID |
|---|---|---|---|---|---|---|
| SHH2a | AtSHH2 | AT3G18380 | SpSHH2a | Sp9509d020g005890 | WbSHH2a | Wb7522d032g001970 |
| SHH2b | | | SpSHH2b | Sp9509d009g007120 | | |
| DRB3 | AtDRB3 | AT3G26932 | | | | |
| DRB5 | AtDRB5 | AT5G41070 | | | | |
| DRB6 | | | SpDRB6 | Sp9509d019g004650 | WbDRB6 | Wb7522d005g006160 |
| ASI1 | AtASI1 | AT5G11470 | SpASI1 | Sp9509d005g000050 | WbASI1 | Wb7522d021g003330 |

**Figure S6. List of gene IDs.** List of Gene IDs of silencing orthologues in *W. brasiliensis, S. polyrhiza* and *A. thaliana.*

### SUPPLEMENTAL FIGURE S7

| | | 1 | 2 | 3 | 4 | 5 | 6 | 7 |
|---|---|---|---|---|---|---|---|---|
| Wb7522d041g000520 **(DCL1a)** | 1 | | 86.33 | 68.69 | 25.48 | 26.57 | 19.53 | 16.12 |
| Wb7522d041g000580 **(DCL1b)** | 2 | 1591 | | 64.21 | 22.99 | 24.32 | 16.71 | 13.22 |
| **SpDCL1** | 3 | 1367 | 1274 | | 23.59 | 24.86 | 16.59 | 13.41 |
| Wb7522d081g000400 **(DCL4)** | 4 | 504 | 460 | 505 | | 60.20 | 20.53 | 16.63 |
| **SpDCL4** | 5 | 512 | 473 | 520 | 983 | | 19.45 | 16.78 |
| **SpDCL3** | 6 | 410 | 359 | 381 | 390 | 363 | | 35.69 |
| Wb7522d021g002440 **(DCL3)** | 7 | 303 | 254 | 279 | 284 | 280 | 555 | |

Upper triangle: **protein similarity %**; lower triangle: **number of identities (aa)**

**Figure S7. Pairwise comparison of DCL proteins identified in *W. brasiliensis.*** Pairwise aminoacid comparison between the *W. brasiliensis* and *S. polyrhiza* DCL proteins. Table shows number of aminoacids that are identical, and global protein similarity (%).

#### SUPPLEMENTAL FIGURE S8

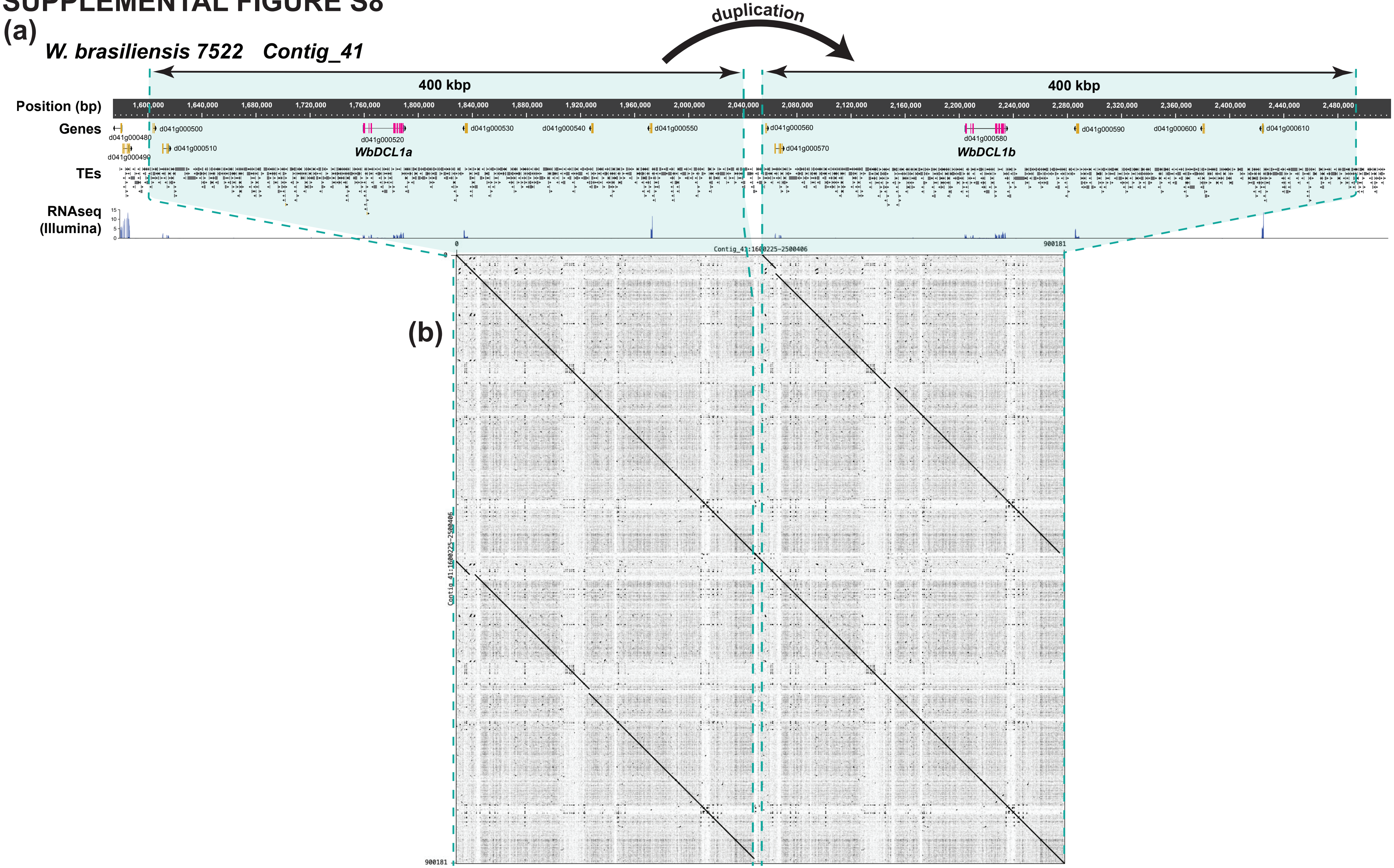


**Figure S8. Genome duplication of DCL1 in *W. brasiliensis*. (a)** Genome screenshot illustrating the WbDCL1 genomic loci. Genomic coordinates, gene and TE annotations and Illumina RNA-seq read coverage are shown as tracks. WbDCL1a (Wb7522d041g000520) and WbDCL1b (Wb7522d041g000580) are highlighted in pink. Color shaded areas delimited by dashed lines indicate the duplicated regions. **(b)** Self dot plot of the genomic sequence highlighted in a) with 100 bp window was used as word size. Main diagonal (top left to bottom right) indicates perfect sequence identity with itself. Parallel lines to the diagonal indicates the presence of duplicated segments within the sequence (delimited with blue dashed lines).

### SUPPLEMENTAL FIGURE S9

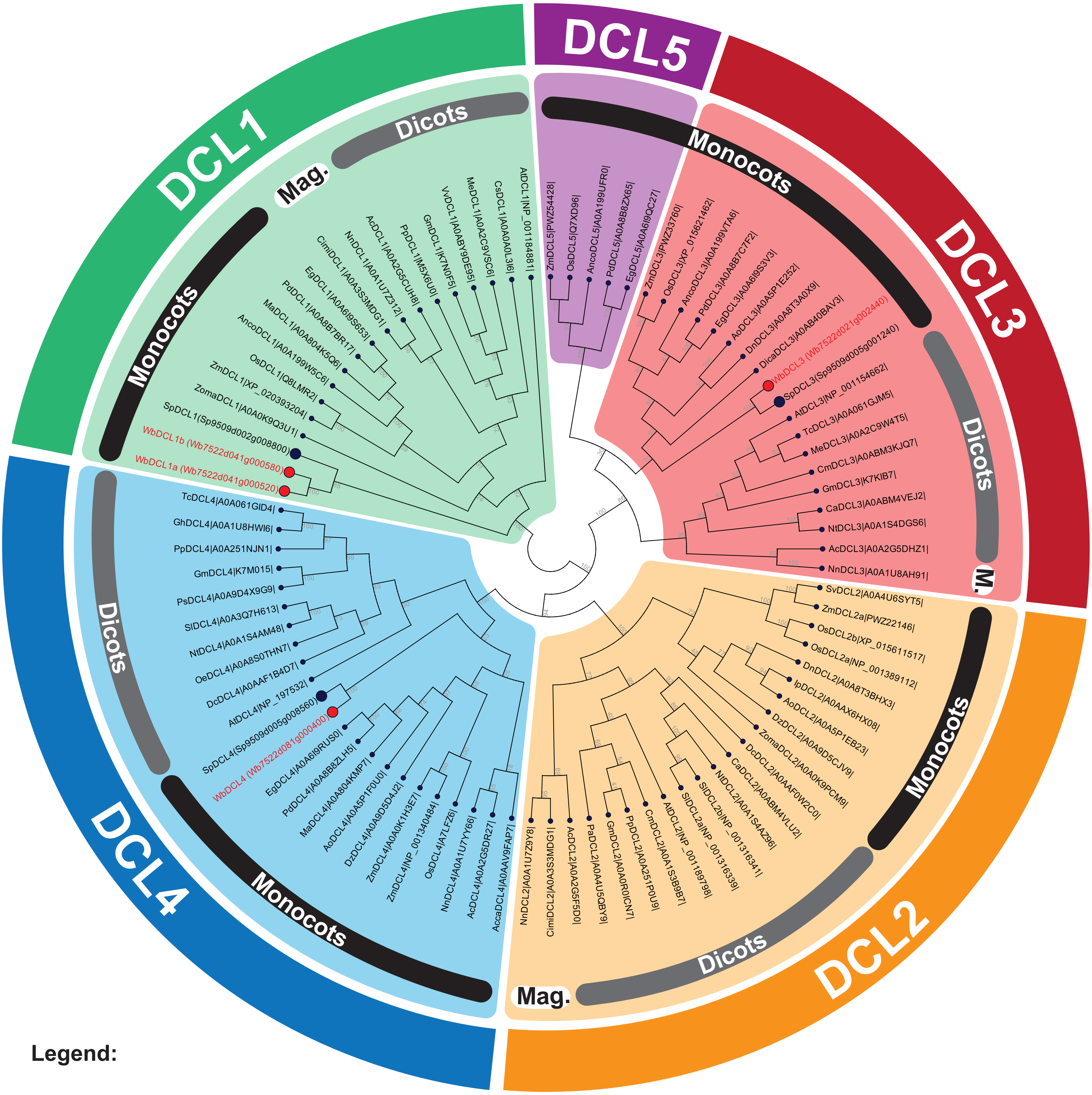


#### Legend:

**Clade *Magnoliids***

Cimi: *Cinnamomum micranthum*
Nn: *Nelumbo nucifera*

**Clade *Dicots***

Ac: *Aquilegia coerulea*
At: *Arabidopsis thaliana*
Ca: *Coffea arabica*
Cs: *Cucumis sativus*
Dc: *Daucus carota*
Gh: *Gossypium hirsutum*
Gm: *Glycine max*
Me: *Manihot esculenta*
Nt: *Nicotiana tabacum*
Oe: *Olea europea*
Pa: *Populus alba*
Pp: *Prunus persica*
Ps: *Pisum sativum*
Sl: *Solanum lycopersicum*
Tc: *Theobroma cacao*

**Clade *Monocots***

Acca: *Acorus calamus*
Anco: *Ananas comosus*
Ao: *Asparagus officinalis*
Dica: *Discorea cayennensis*
Dn: *Dendrobium cayennensis*
Dz: *Discorea zingiberensis*
Eg: *Elaeis guineensis*
Ma: *Musa acuminata*
Os: *Oryza sativa*
Pd: *Phoenix dactilifera*
Sp: *Spirodela polyrhiza*
Sv: *Setaria viridis*
Wb: *Wolffia brasiliensis*
Zoma: *Zostera marina*
Zm: *Zea mays*

**Figure S9. Phylogenetic analysis of DCL proteins in *W. brasiliensis*.** Unrooted maximum likelihood phylogenetic tree of DCL proteins from several diverse angiosperm species. The DCL proteins identified in the *W. brasiliensis* genome were also included. The tree was constructed with 1000 bootstrap replicates to assess node support. Distinct protein clades are demarcated by color-coded background shading. *W. brasiliensis* proteins are highlighted in bold red and were annotated based on their phylogenetic clustering. For those, and for DCL proteins in *S. polyrhiza,* gene model IDs are provided, while UniProt accessions are listed for all other taxa. Comprehensive data, including the underlying amino acid sequences, multiple sequence alignments, and source tree files, are accessible in the supplementary raw data files.

### SUPPLEMENTAL FIGURE S10

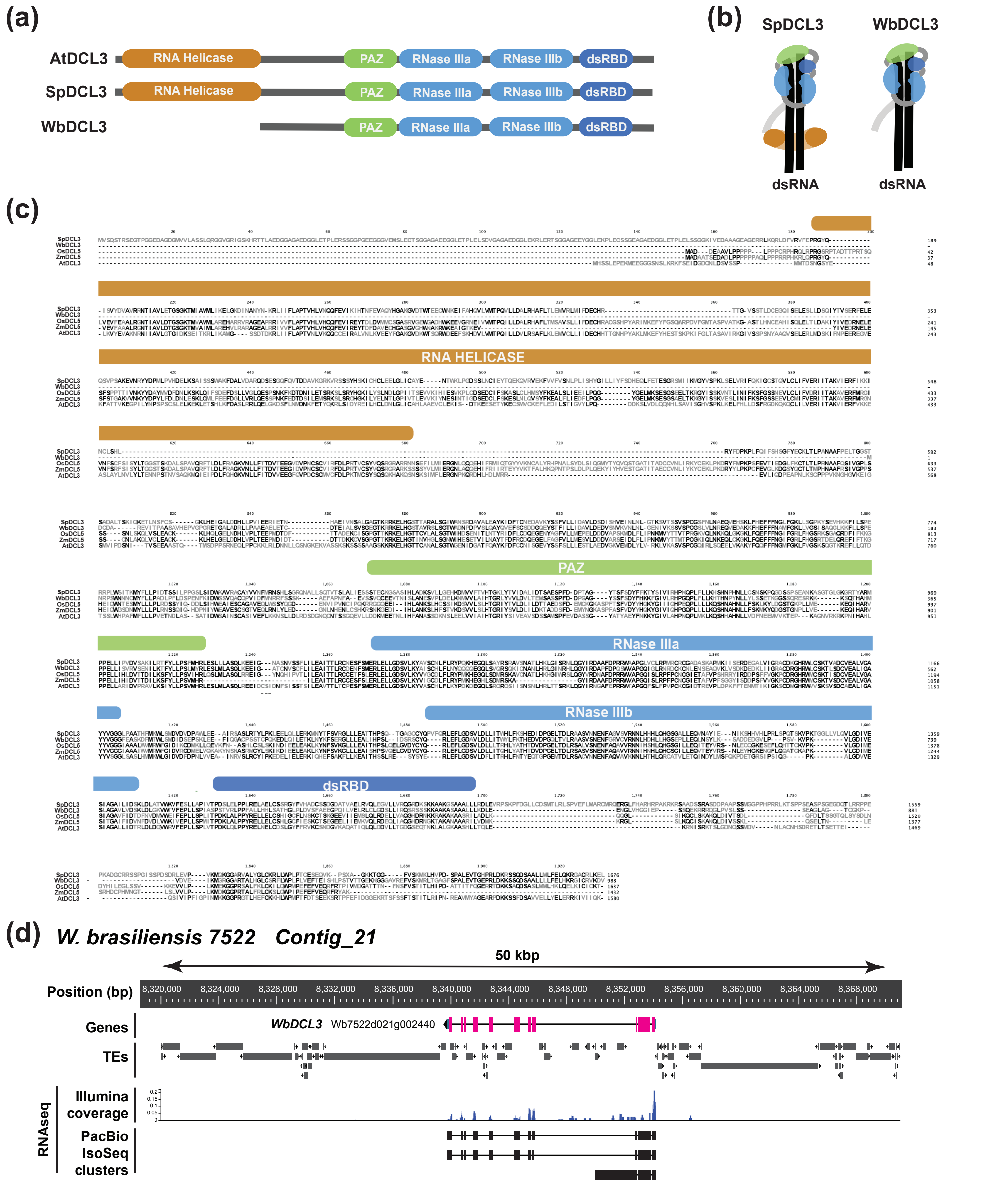


**Figure S10. Structural analysis of WbDCL3. (a)** Schematic representation of the domain organization of WbDCL3 (Wb7522d021g002440) and its comparison with *S. polyrhiza* #9509 SpDCL3 and *A. thaliana* Col-0 AtDCL3. **(b)** Comparative Illustration of SpDCL3 and WbDCL3 bound to dsRNA, suggesting WbDCL3 may still be completely functional despite the lack of the helicase domain. **c)** Protein allignment of WbDCL3 and other angiosperms illustrates the level of conservation at the identified domains. Proteins used were SpDCL3, *Oryza sativa* DCL5, *Zea mays* DCL5 and AtDCL3. DCL5 proteins from maize and rice were used due to their closest phylogenetic relationship with DCL3 than other DCL proteins identified on those species. Aminoacid conservation was highlighted in black, while non-conserved aminoacids were colored in grey. Solid colored bars indicate and label the conserved domains. **(d)** Genome snapshot of the WbDCL3 *locus* (Wb7522d021g002440), in pink. The lack of helicase domain is not due to a misannotation of the WbDCL3, as confirmed by Illumina RNA-seq and PacBio Isoseq reads. Genomic coordinates, gene and TE annotations, Illumina RNA-seq coverage and PacBio Isoseq read clusters are shown as tracks.

### SUPPLEMENTAL FIGURE S11

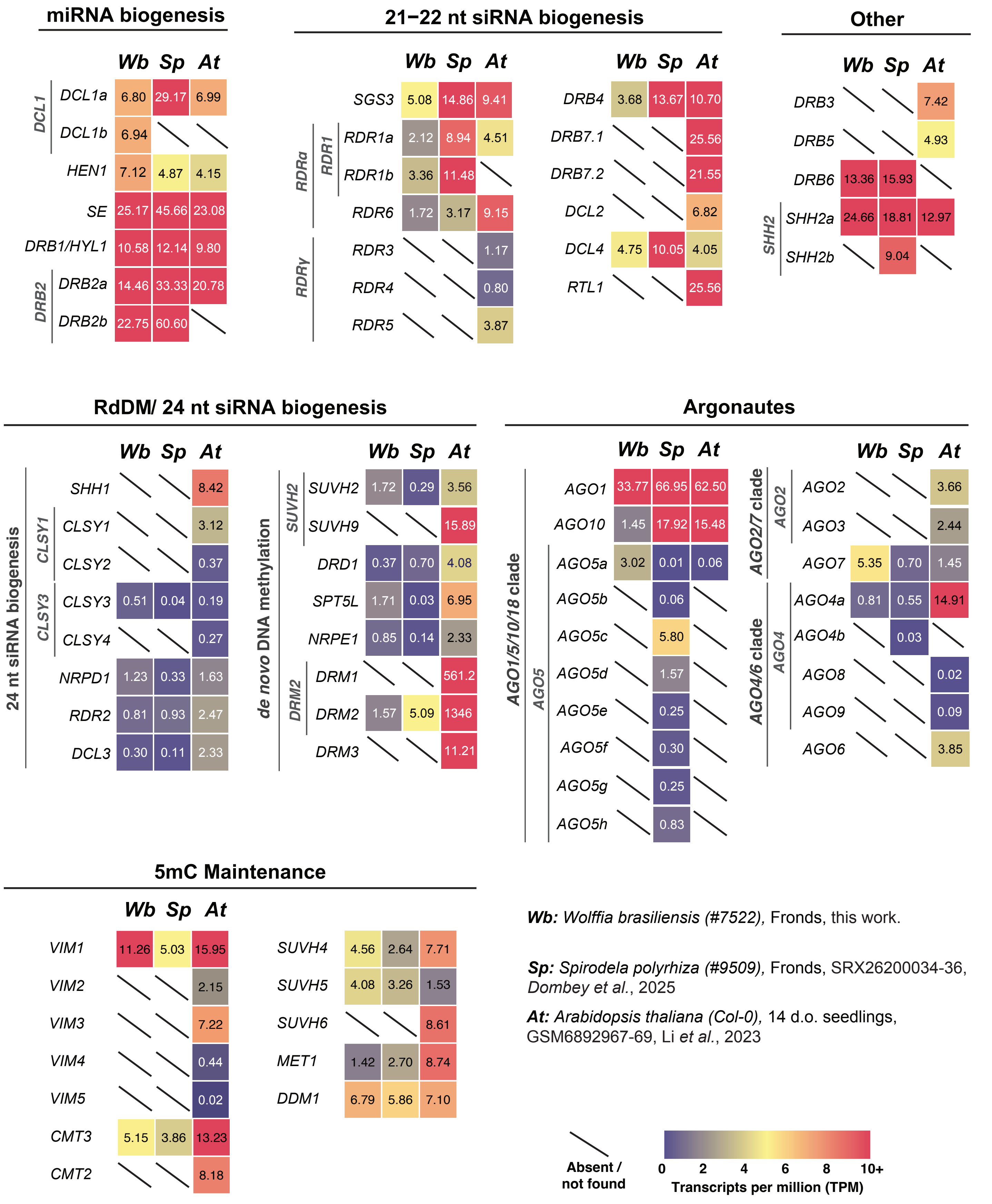


**Figure S11. Gene expression profile of gene silencing components in *W. brasiliensis* and its comparison with *S. polyrhiza* and *A. thaliana*.** Heatmap shows the expression levels (Transcripts per million; TPM) of key ortholog genes. Gene names are indicated next to the expression values. Color gradient represents level of expression, and a diagonal line indicates the absence of the gene in the corresponding genome. Sp= *Spirodela polyrhiza* #9509, Wb=*Wolffia brasiliensis* #7522, At= *Arabidopsis thaliana* (Col-0).

### SUPPLEMENTAL FIGURE S12

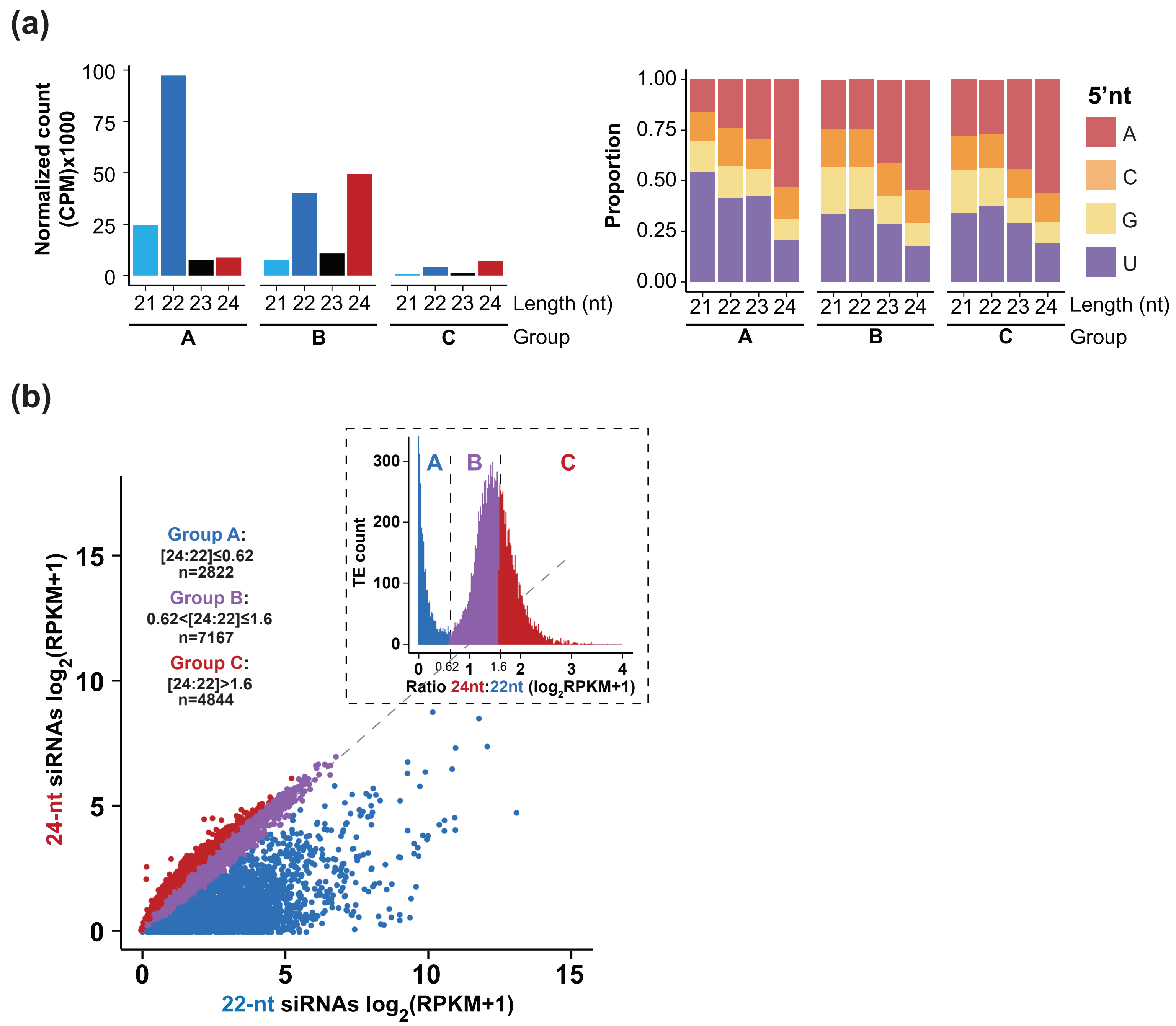


**Figure S12. Profile of siRNAs from siRNA producing TEs in *W. brasiliensis*. (a)** Left: Barplot illustrating the normalized abundance of reads from different siRNA sizes for each subpopulation of siRNA producing TEs described in Fig 3f. Right: Stacked barplot showing the 5' nucleotide bias of TE-derived 21-24nt siRNAs on each siRNA producing TE category. **(b)** siRNA-producing TE subpopulations based on the relative abundance of uniquely mapped 22 and 24nt siRNAs. Inset plot shows a histogram of the ratio of 24/22nt siRNA levels categorized as described in the Material and Methods section. TE categories are A: 22nt siRNA producing bias; B: No 22 or 24nt siRNA producing bias. C: 24nt siRNA producing bias. Left and Right thresholds are indicated below the category labels, alongside with the number of TEs corresponding to each category. Diagonal dashed line represents the 1:1 ratio.

#### SUPPLEMENTAL FIGURE S13

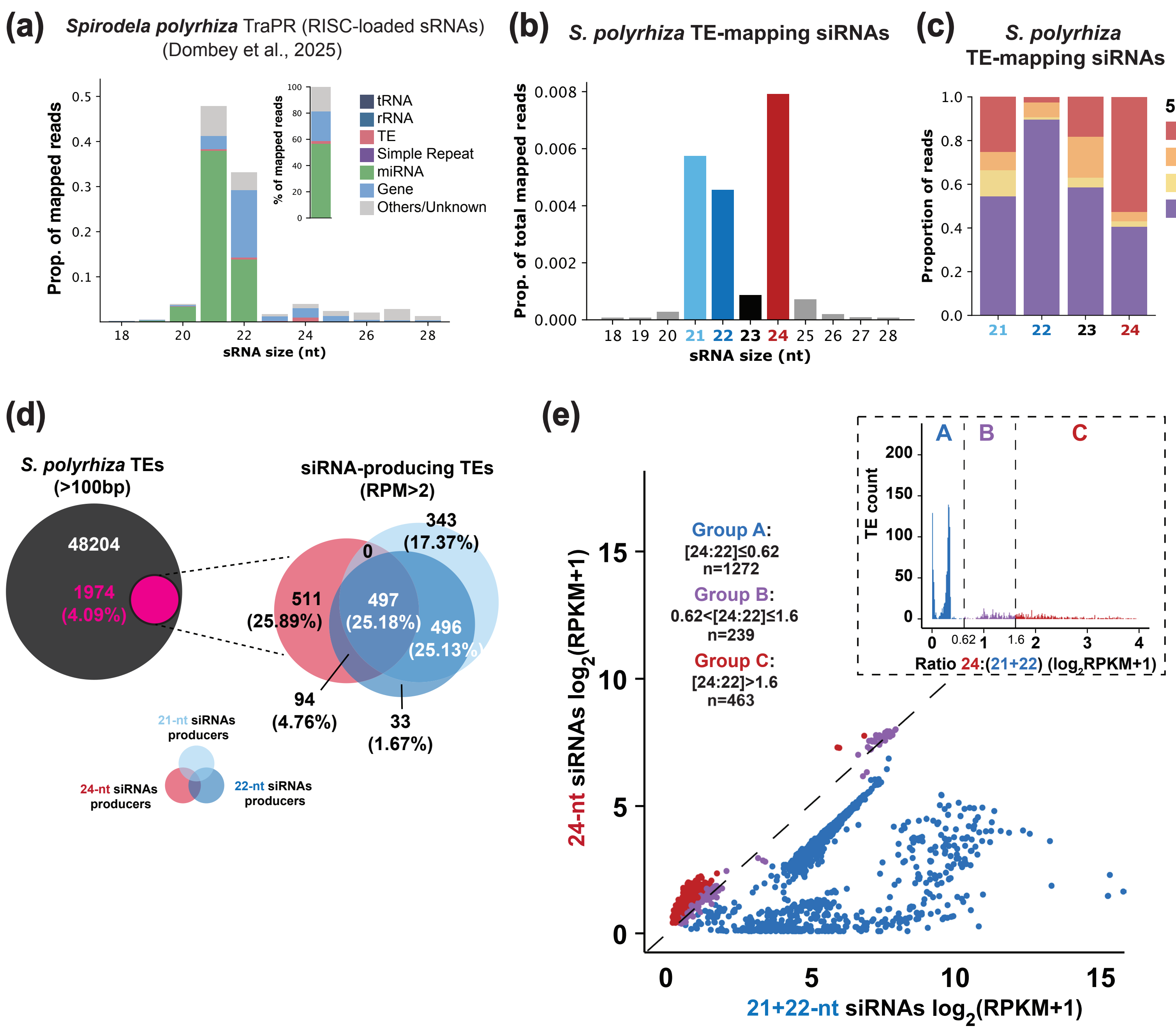


**Figure S13. Analysis of TE-derived small RNA in *S. polyrhiza*. (a)** Size distribution and genomic mapping of TraPR-purified small RNAs. Inset shows percentage (%) of all mapped siRNA per feature. **(b)** Proportion of TE-mapping siRNAs distributed by size. **(c)** 5'nt bias distribution of TE-mapping siRNAs. **(d)** Venn diagrams of siRNA-producing (>2 reads per million; RPM) over total TE annotations and overlap of 22- and 24-nt siRNA-producing TEs. **(e)** Abundance (in reads per kb per million reads; RPKM+1) of 21-22- and 24-nt siRNAs for each siRNA-producing TE colored by groups determined by their 24-nt:21-22-nt ratio as defined in inlet. Diagonal line represents the 1:1 ratio.

### SUPPLEMENTAL FIGURE S14

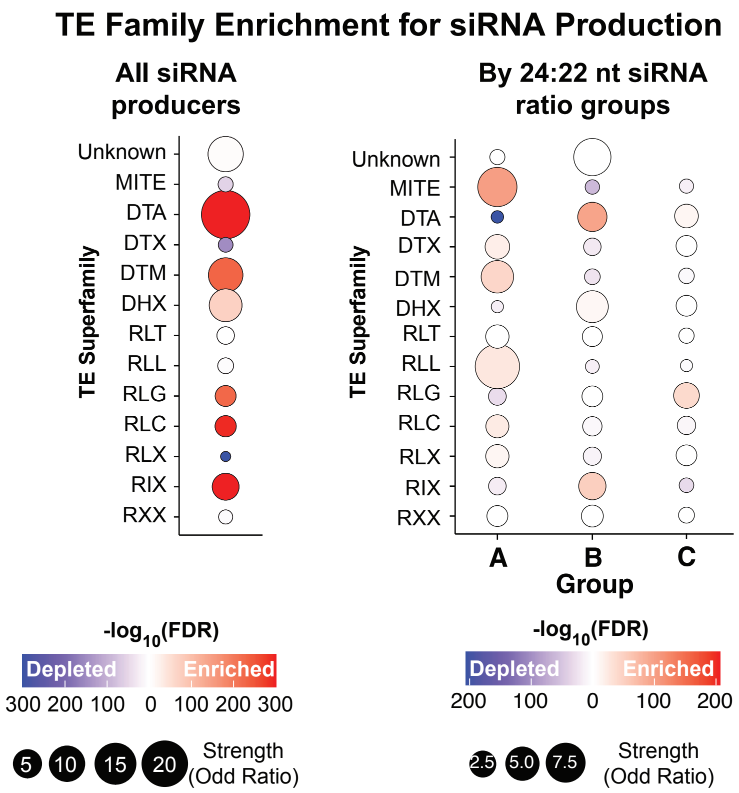


**Figure S14. TE Family siRNA enrichment analysis in *W. brasiliensis*.** Bubble heatmap showing the TE Family association with siRNA production. Enrichment/depletion of siRNA in different TE families was determined using two-sided Fisher's exact tests for each family applying the Benjamini-Hochberg (BH) correction. The heatmap represents the results of Fisher's exact tests, showing the significance and direction of siRNA association for each family. Size of the bubble represents the magnitude of the enrichment/depletion, and color, indicated by the Odd Ratio (OR) shows the direction. OR >1 was considered as "enriched" and colored in red. OR <1 was considered as "depleted" in siRNA and colored in blue. Gradient color intensity illustrates the significance of the enrichment/depletion and corresponds to the -log10 of the adjusted p-value (FDR).

### SUPPLEMENTAL FIGURE S15

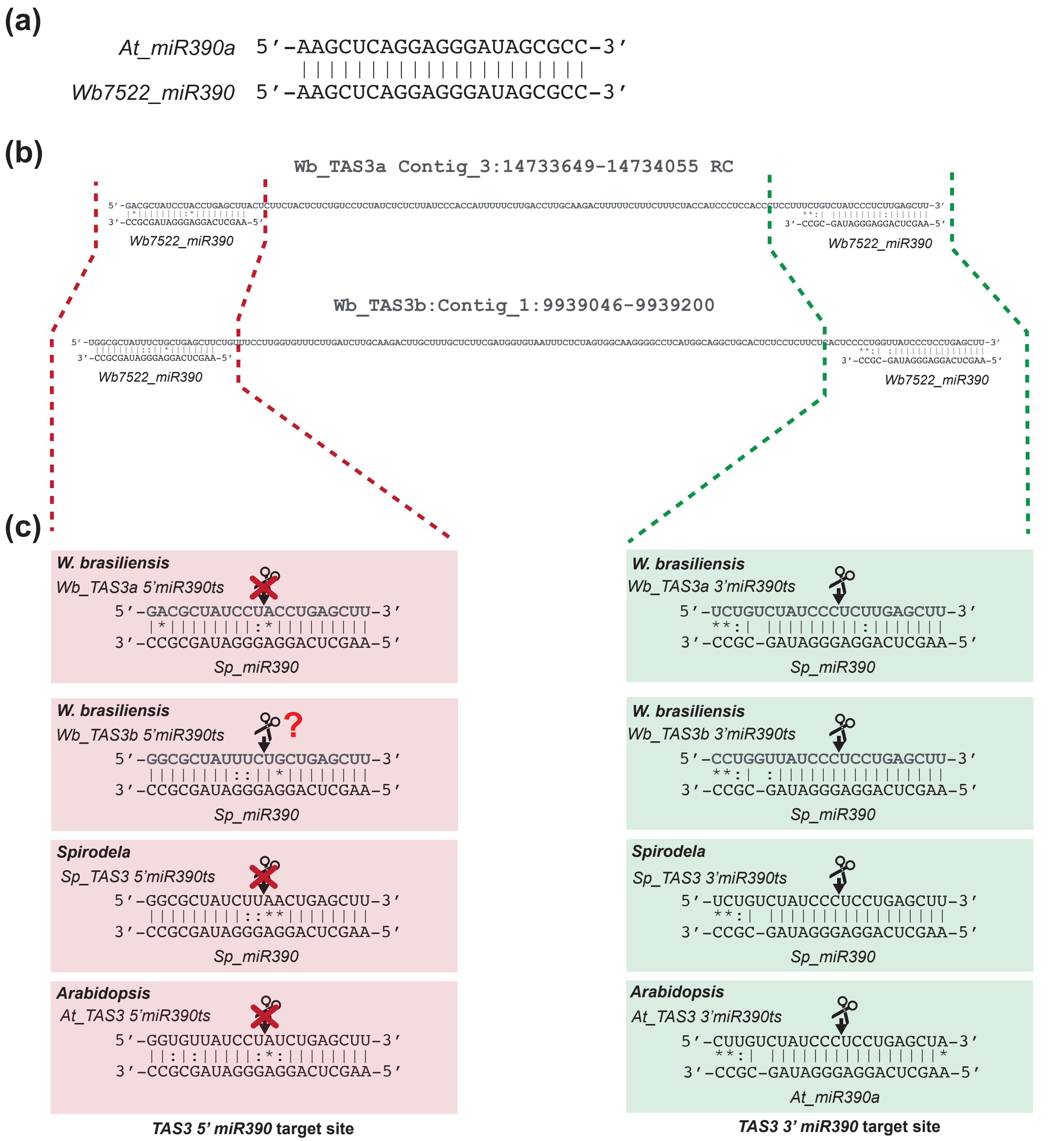


**Figure S15. TAS3 loci identified in *W. brasiliensis*. (a)** Sequence alignment of miR390 from *A. thaliana* (At_miR390a) and *W. brasiliensis* (Wb7522_miR390). Perfect alignment is observed between the two miRNAs. **(b)** *W. brasiliensis* identified TAS3a and TAS3b genomic sequences. Dashed lines delimit miR390 target sites, and the alignment with the miR390 sequence is shown. Vertical lines indicate perfect match. An asterisk "*" indicates mismatch and two dots ":" indicate semi-conserved substitutions. **(c)** Target site analysis of WbTAS3a and WbTAS3b, and its comparison with the corresponding *S. polyrhiza* and *A. thaliana* TAS3 miR390 target sites. Scissors indicate miR390-RISC induced cleavable sites.

### SUPPLEMENTAL FIGURE S16

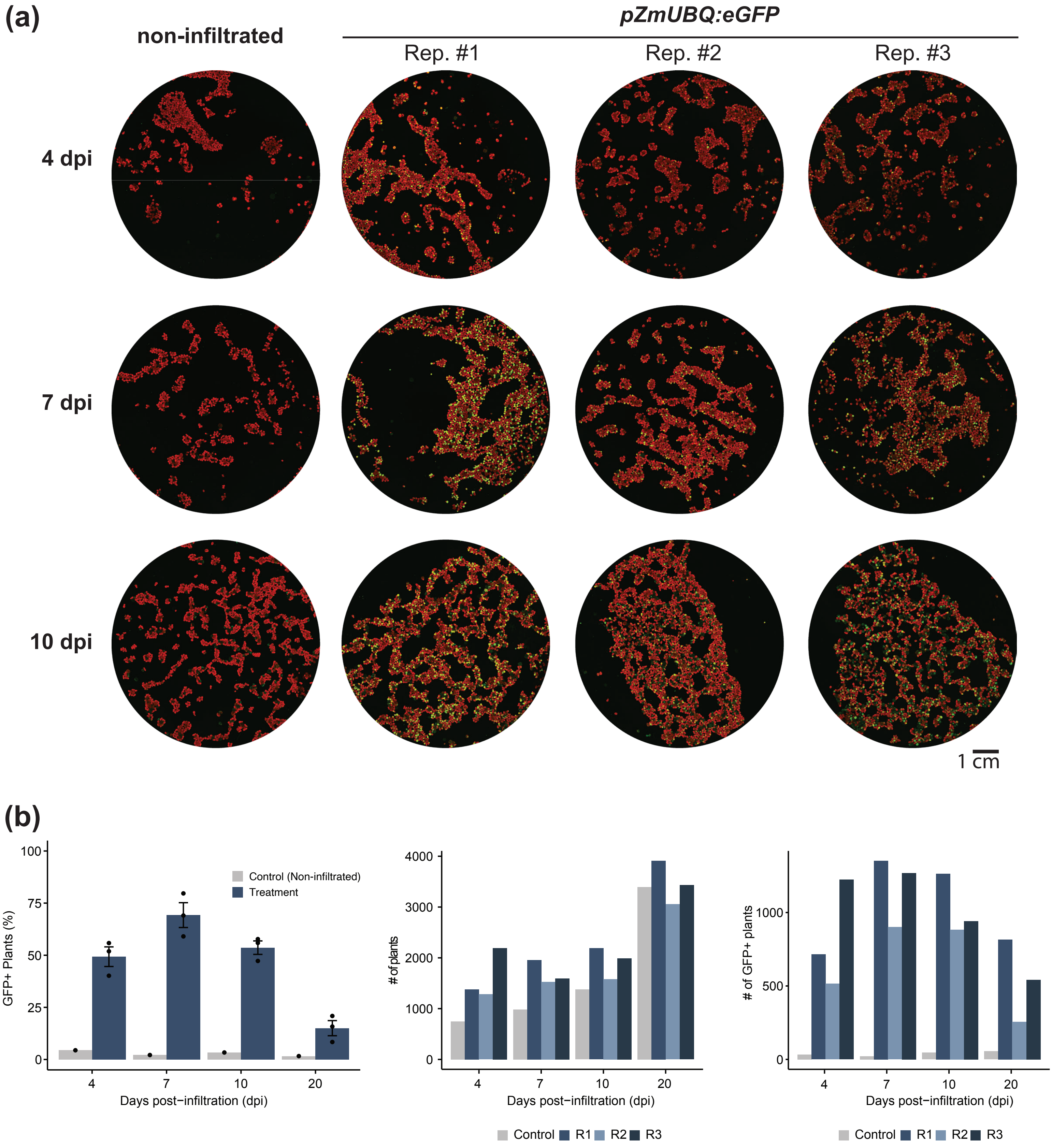


**Figure S16.** ***Agrobacterium*****-mediated transient expression on *W. brasiliensis* fronds. (a)** Representative fluorescence images of non-infiltrated (neg. Control) and pZmUBQ:eGFP transformed fronds from the three biological replicates, across different days after infiltration (dpi). Images were taken as described in the Materials and Methods section. **(b)** Barplot showing fluorescence quantification data from the transformed and control plates at different time points after infiltration. Left: Control bar height represents a single plate, while treatment bar represents the average of the three biological replicates. Error bars represent the SD from the mean. Center and Right: Absolute number of GFP+ fronds and total amount of fronds quantified during the experiment time, respectively. R1-3=Replicates 1-3.

### SUPPLEMENTAL FIGURE S17

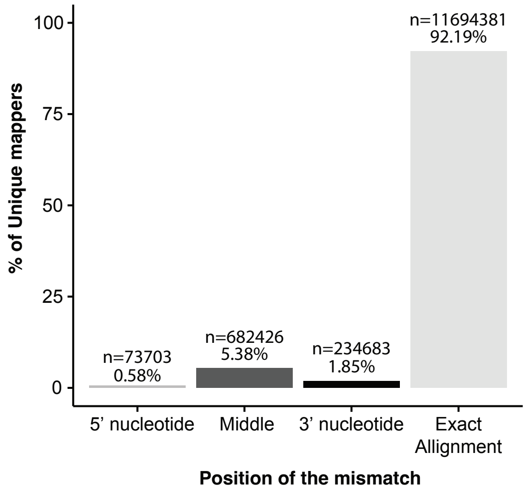


**Figure S17. Analysis of non-templated nucleotides in uniquely mapped siRNA in Wolffia.** Bar-plot representing the percentage of unique mappers that show a mismatch in their alignment to the genome at different positions of the read. Percentage of reads with a perfect alignment are also shown.

#### SUPPLEMENTAL FIGURE S18

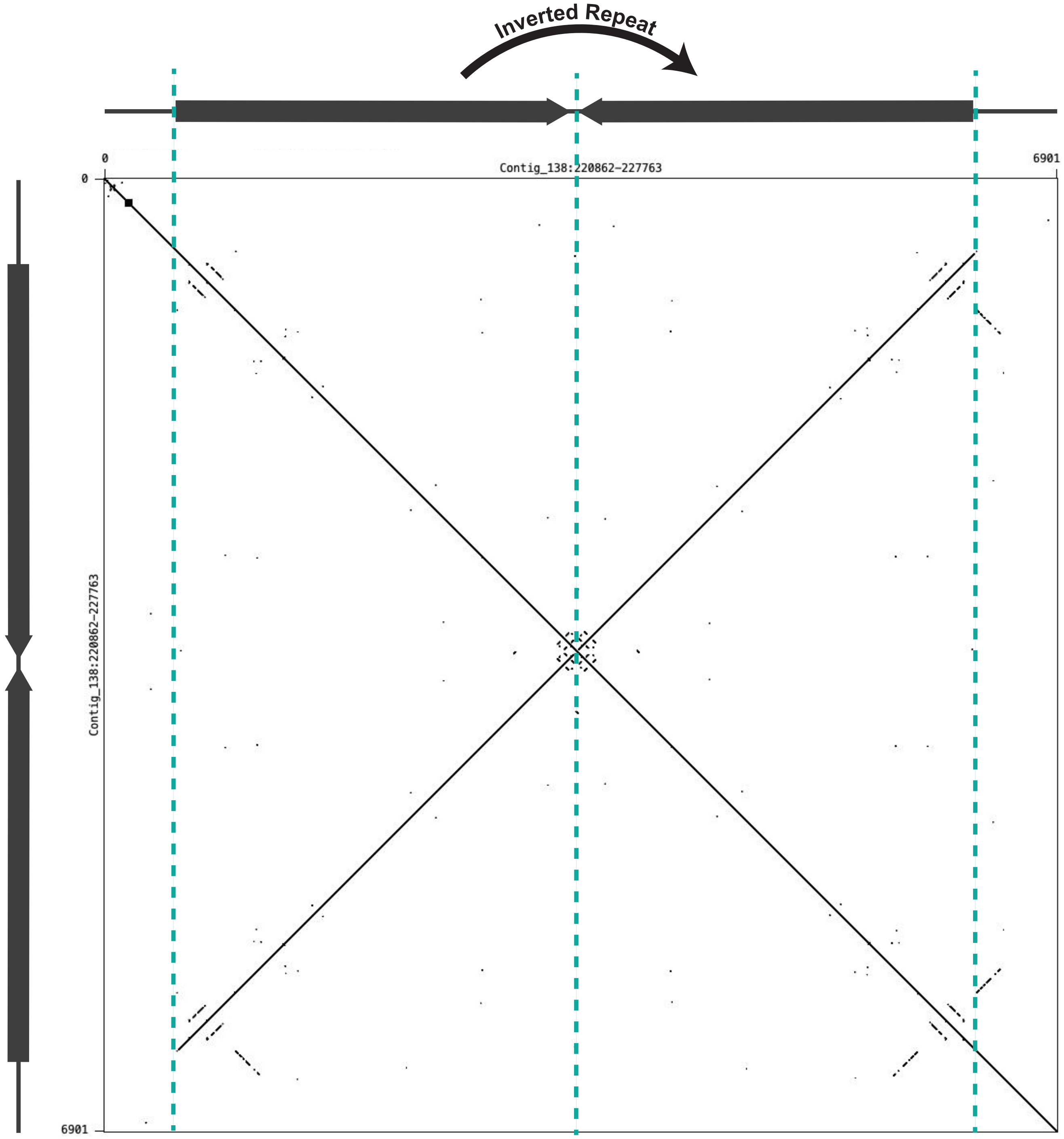


**Figure S18. Example of an Inverted Repeat in Wolffia.** Self dot-plot analysis of an Inverted repeat sequence. A 100 bp window parameter was used as word size. Main diagonal (top left to bottom right) indicates perfect sequence identity with itself. Perpendicular (bottom left to top-right) diagonal indicates the presence of an inverted repeat covering most of the identified sequence (delimited with blue dashed-lines).

### SUPPLEMENTAL FIGURE S19

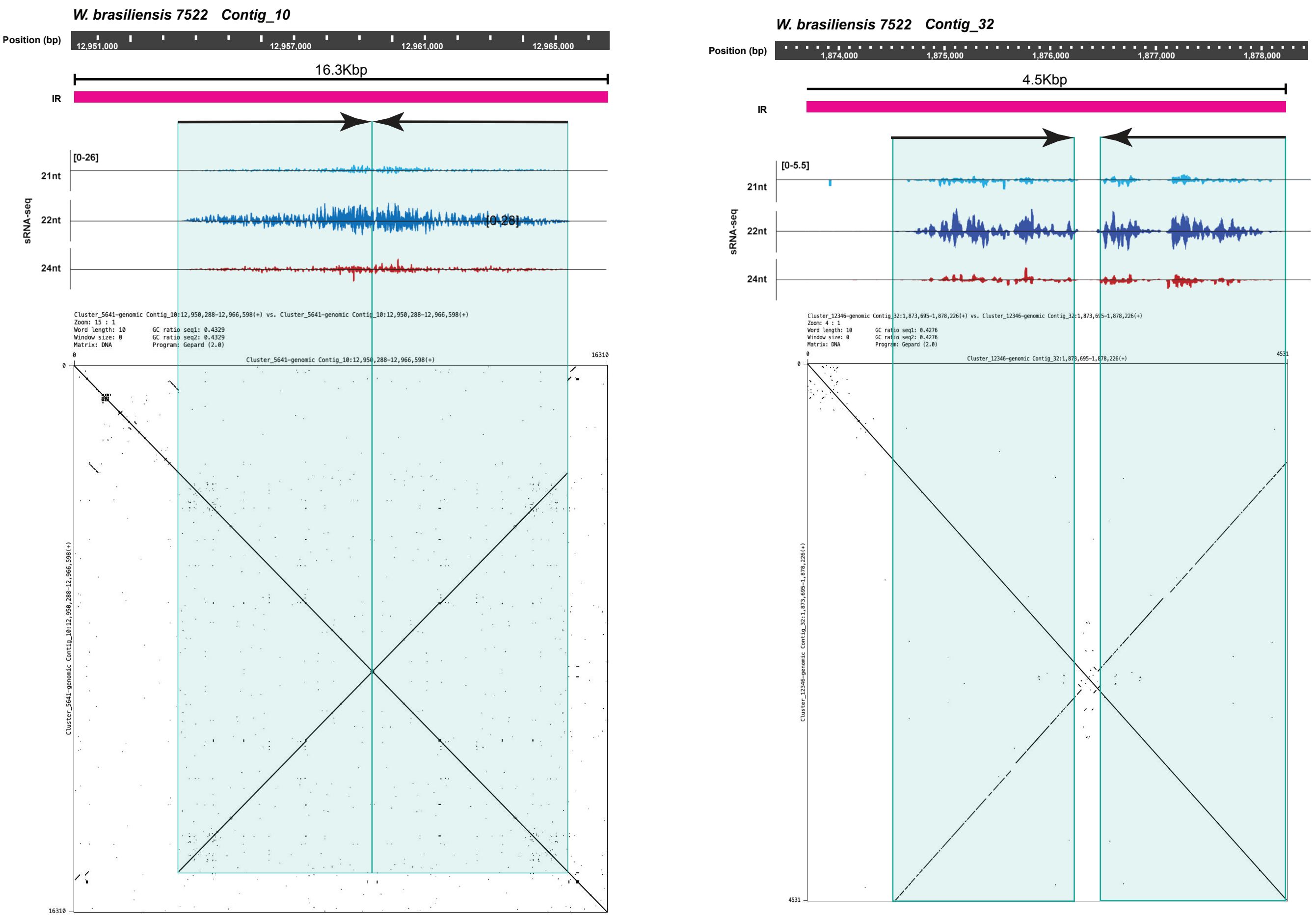


**Figure S19. Example of additional siRNA-producing IR+ clusters in Wolffia.** Genome browser snapshots of two Inverted repeat (IR) containing siRNA clusters. Genomic coordinates, IR-containing siRNA cluster annotation and siRNA-seq read coverage are presented as tracks. Coverage values are expressed in reads per million (RPM)**.** Inverted repeat stems are annotated with black arrows and color-shaded background. Bottom panel depicts a self dot-plot analysis. A 100 bp window parameter was used as word size. Main diagonal (top left to bottom right) indicates perfect sequence identity with itself. Perpendicular (bottom left to top-right) diagonal indicates the presence of the inverted repeat covering most of the identified sequence (delimited with blue dashed lines). Both examples illustrate how siRNA production from the clusters is almost exclusively restricted to the inverted repeat.

### SUPPLEMENTAL FIGURE S20

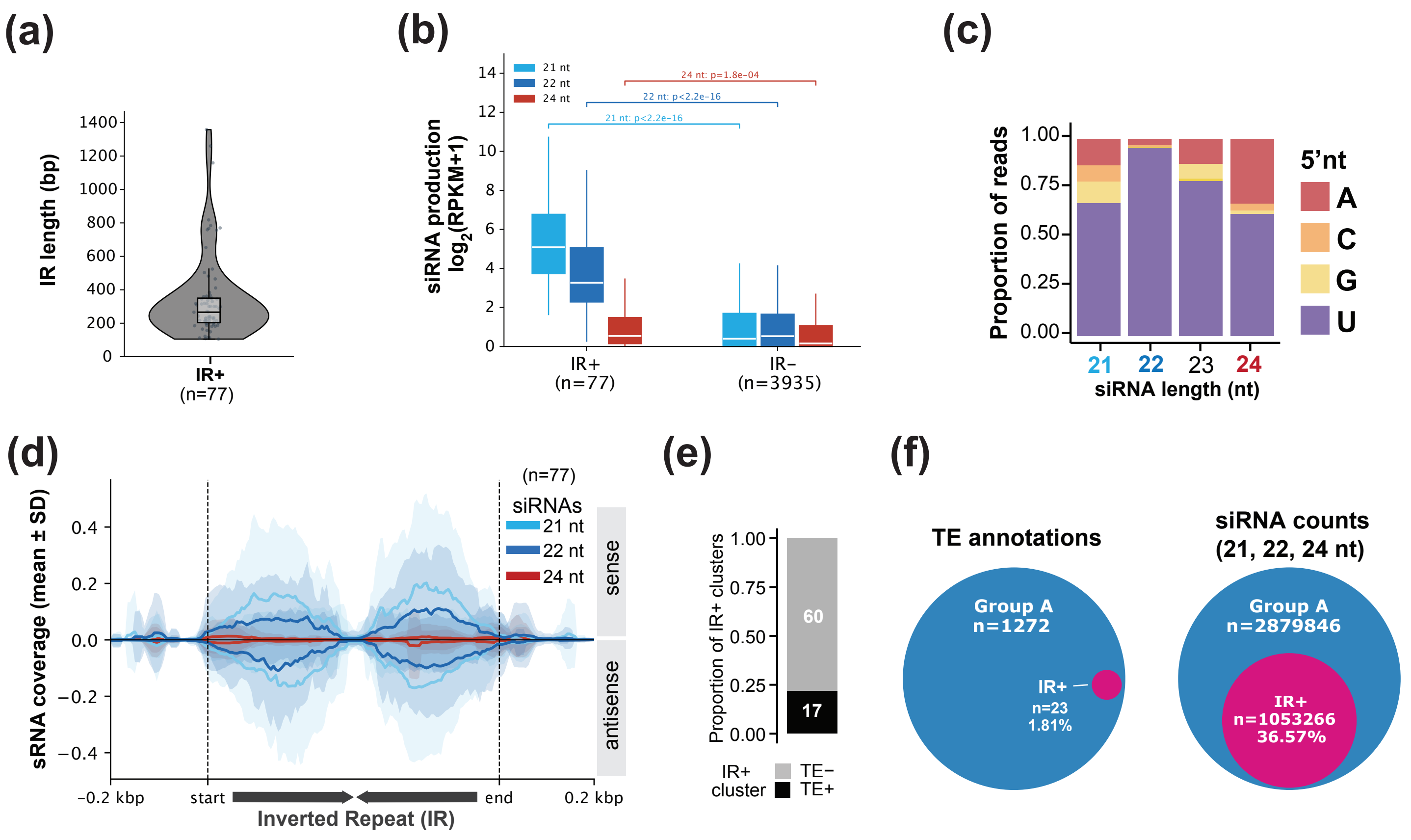


**Figure S20. Identification and characterization of endogenous inverted repeats in *S. polyrhiza*. (a)** Length distribution in bp of identified IRs using the strategy described in FIgure 5c. **(b)** 21-, 22-and 24-nt siRNA levels (RPKM+1) of IR+ and IR- siRNA clusters. P-values determined by Wilcoxon rank sum test. **(c)** 5'nt bias of IR+ cluster-derived siRNAs. **(d)** siRNA distribution across identified IRs and their flanking regions. Solid line and shaded area represent the mean normalized coverage and the SD respectively. **(e)** Proportion of IR+ siRNA clusters overlapping with TE annotations. **(f)** Venn diagrams of Group A TEs overlapping with IR+ clusters and their contribution to Group A-mapping siRNAs.

### SUPPLEMENTAL FIGURE S21

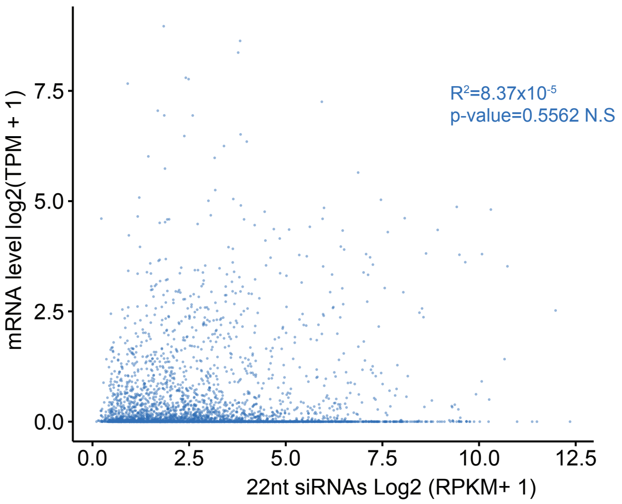


**Figure S21. Correlation between 22nt-siRNA production and expression on TEs.** Dot plot illustrating the relationship between 22nt-siRNA levels and RNA expression (Transcripts per million; TPM). Pearson's correlation coefficient and the associated p- value were calculated using log-transformed coordinates.

### SUPPLEMENTAL FIGURE S22

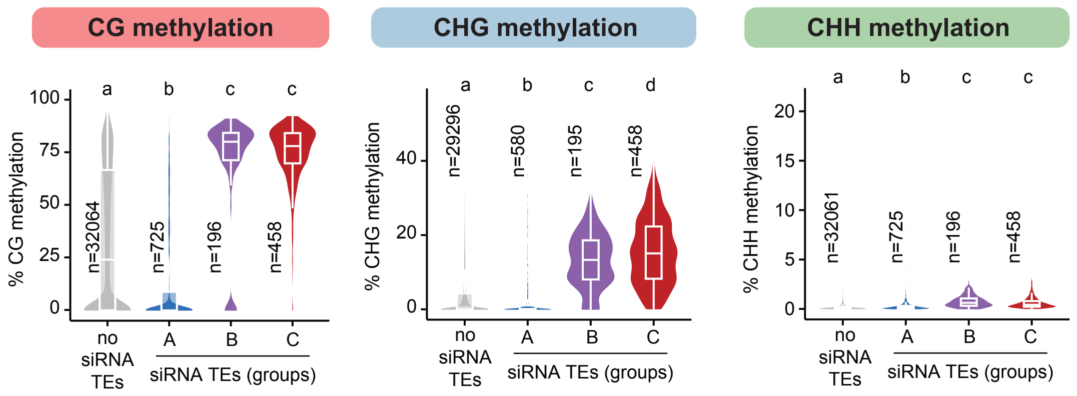


**Figure S22. Distribution of DNA methylation in siRNA TE-groups in *S. polyrhiza*.** Violin and boxplot showing DNA methylation levels of TEs belonging to the different TE siRNA producing categories in *S. polyrhiza*. In all boxplots the median is represented as a solid bar, with box upper and bottom limits representing the first and third quartiles. Whiskers range is 1.5 times the interquartile range. Statistical significance indicated by letters was calculated using a Kruskal-Wallis test followed by a *post-hoc* Dunn's test. Test statistical P-values can be found in Table S3

### SUPPLEMENTAL FIGURE S23

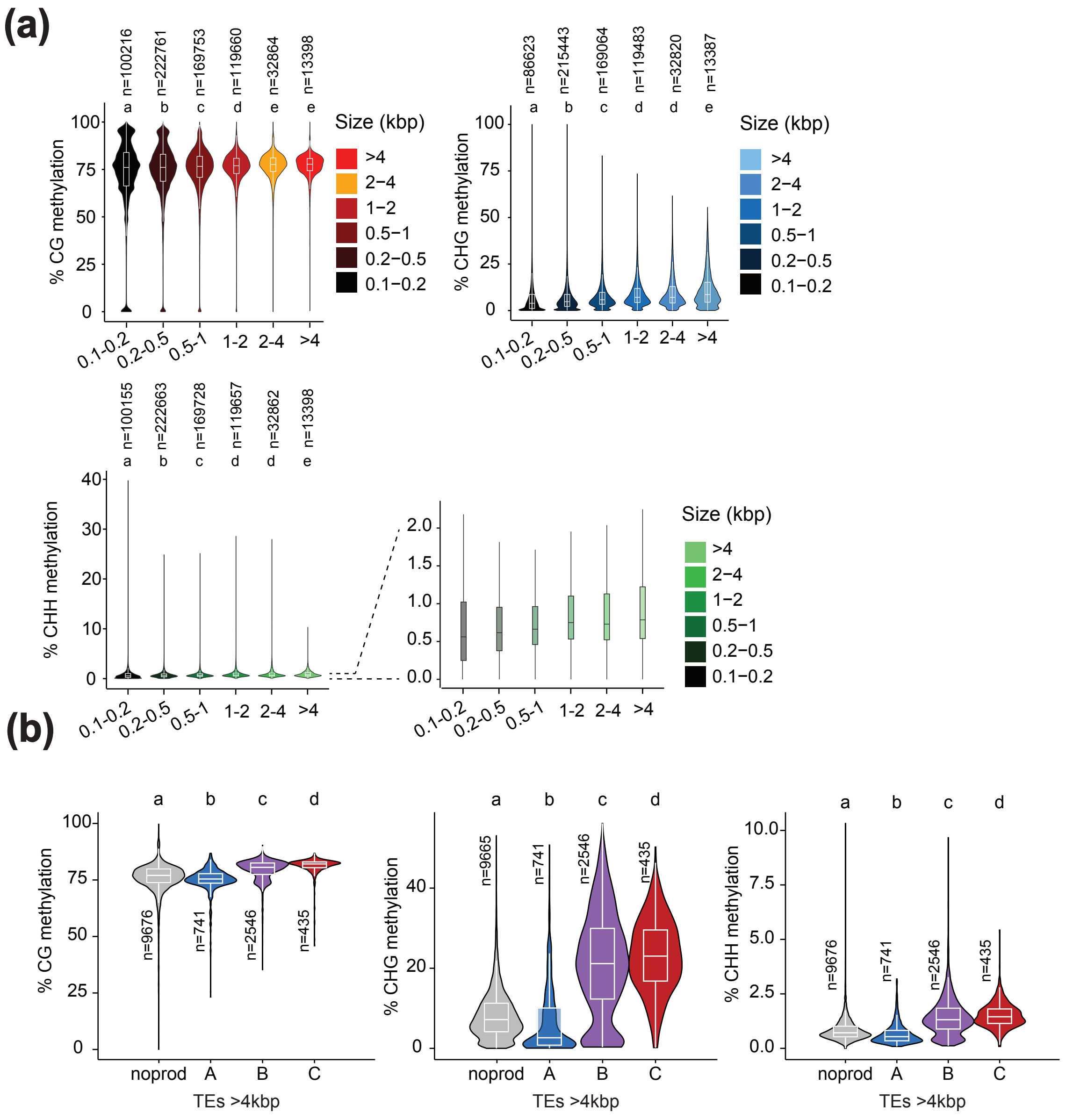


**Figure S23. Effect of TE size on DNA methylation. (a)** Violin and boxplot illustrate the levels of DNA methylation on the CG CHG and CHH contexts across different TE size intervals. For the CHH context, a zoomed-in inset panel of the low methylation range is shown. **(b)** Violin and boxplot showing DNA methylation levels of TEs >4kbp belonging to the different TE siRNA producing categories described in Figure 3f. In all boxplots the median is represented as a solid bar, with box upper and bottom limits representing the first and third quartiles. Whiskers range is 1.5 times the interquartile range. Statistical significance indicated by letters was calculated using a Kruskal-Wallis test followed by a *post-hoc* Dunn's test. Test statistical P-values can be found in Table S3

### SUPPLEMENTAL FIGURE S24

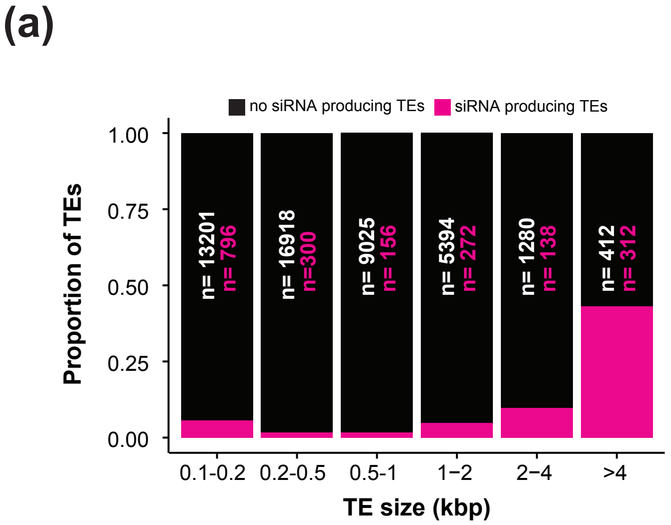


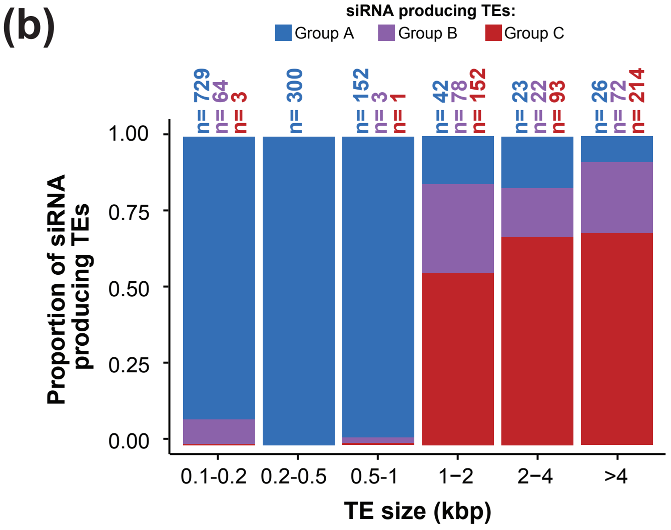


**Figure S24. Impact of TE size on siRNA production in *S. polyrhiza*. (a-b)** Proportion of siRNA-producing TEs (a) and of TE Groups by 24-nt:21-22-nt ratios (b), per size category.

### SUPPLEMENTAL FIGURE S25

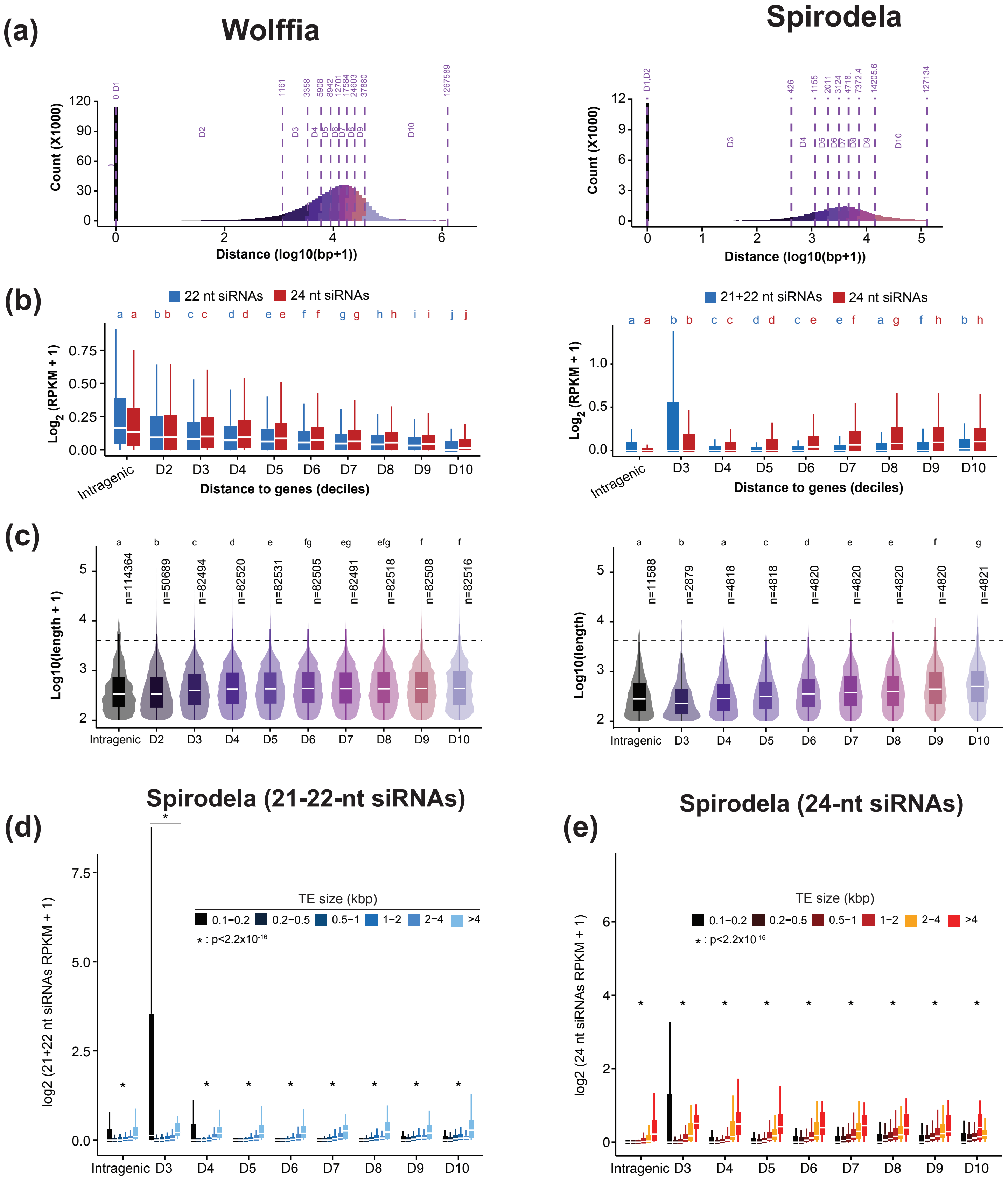


**Figure S25. TE siRNA production correlates with gene distance. (a)** Histogram Illustrating the distribution of the number of TEs according to the genomic distance to the nearest gene, with bars colored by distance deciles (Intragenic, D2-D10). Distances above dashed lines delimiting the different deciles are shown. **(b)** siRNA production (Reads per kilobase pair per million; RPKM) of TE annotations based on their genomic distance to the closest gene deciles (Intragenic, D2-D10). **(c)** TE size distribution in each distance decile in each species. **(d-e)** TE-21+22-nt (d) and 24-nt (e) siRNA levels (RPKM+1) grouped by distance to genes and TE sizes in *S. polyrhiza*. Boxplots represent the median as a solid bar, with box upper and bottom limits representing the first and third quartiles. Whiskers range is 1.5 times the interquartile range. Statistical significance indicated by letters was calculated using a Kruskal-Wallis test followed by a *post-hoc* Dunn's test. All test statistical P-values can be found in Table S4, S5.

### SUPPLEMENTAL FIGURE S26

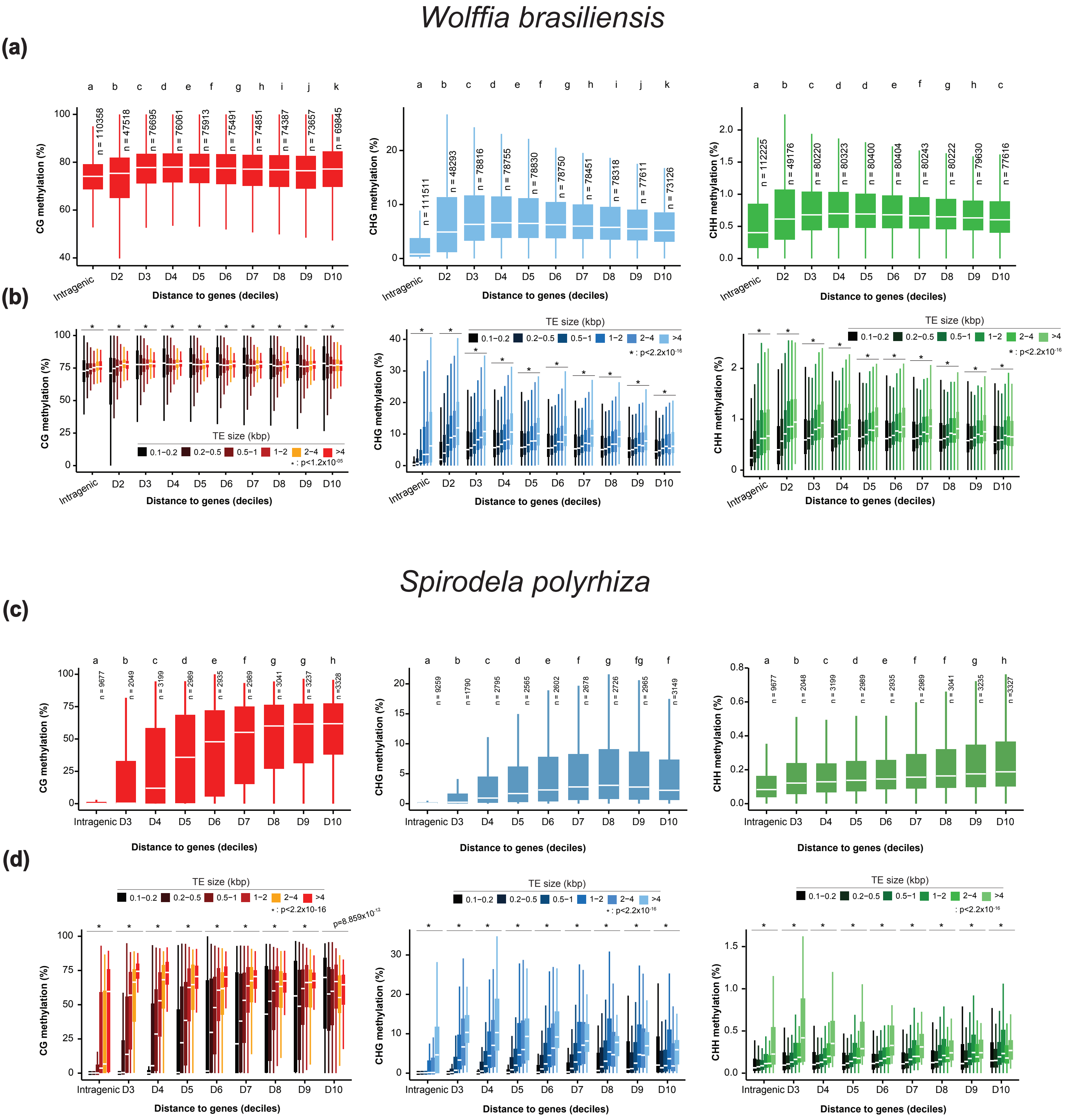


**Figure S26. Analysis of DNA methylation by distance to genes and TE size in Wolffia and Spirodela. (a,b)** Distributions of CG (left), CHG (center) and CHH (right) methylation levels (%) at *W. brasiliensis* TEs, grouped by distance to genes (a) and stratified by TE size (b). The number of TEs per distance class is indicated above each column. **(c,d)** Same analysis as in a-b but in *S. polyrhiza*. (*) P-values<2.2 x 10-16 (Kruskal-Wallis test) between size categories within each decile and intragenic groups. All P-values and n-values can be found in Table S6.

### SUPPLEMENTAL FIGURE S27

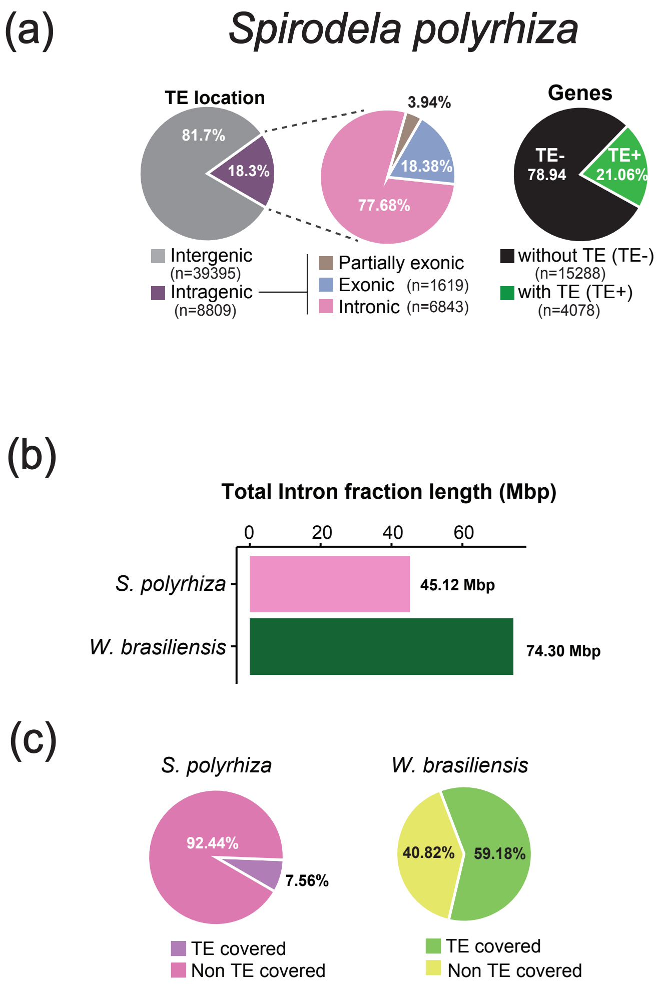


**Figure S27. Intragenic TE distribution in Spirodela and Wolffia. (a)** Percentage of intragenic TEs and their distribution within genes. **(b)** Proportion of genes overlapping with at least one TEs annotation. **(b)** Total intronic sequences in Mbp in each genome. **(c)** Relative (%) of intronic sequences occupied by TE annotations in each species.

### SUPPLEMENTAL FIGURE S28

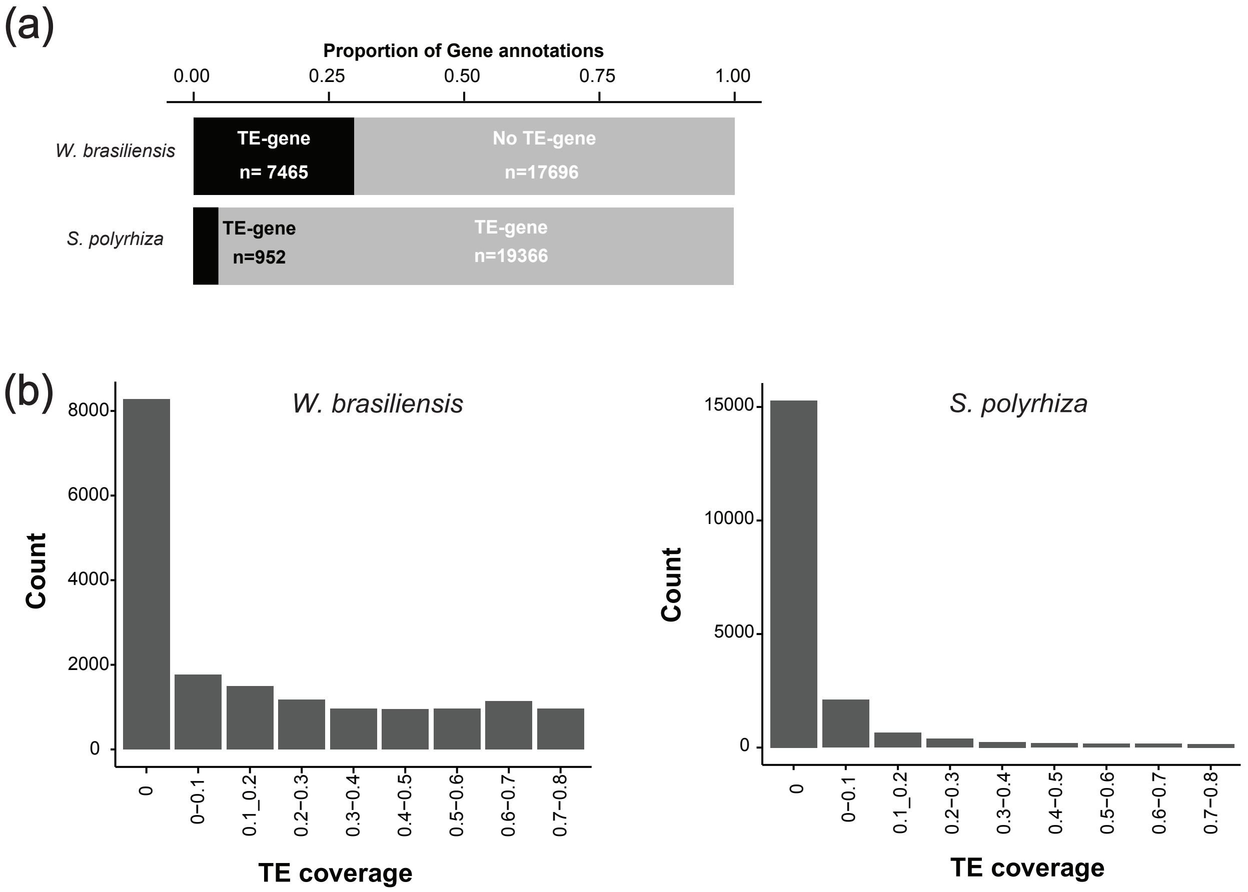


**Figure S28. TE-Genes and TE coverage distribution across gene annotations. (a)** Stacked barplot illustrating the proportion of TE-genes across all gene annotations in Spirodela and Wolffia. TE-genes were defined as gene models in which >80% of the gene-body length overlaps annotated TE sequences lacking canonical TE protein domains for their classification and are therefore retained in the gene set. (**b)** Distribution of genes based on their TE coverage in Spirodela and Wolffia.

### SUPPLEMENTAL FIGURE S29

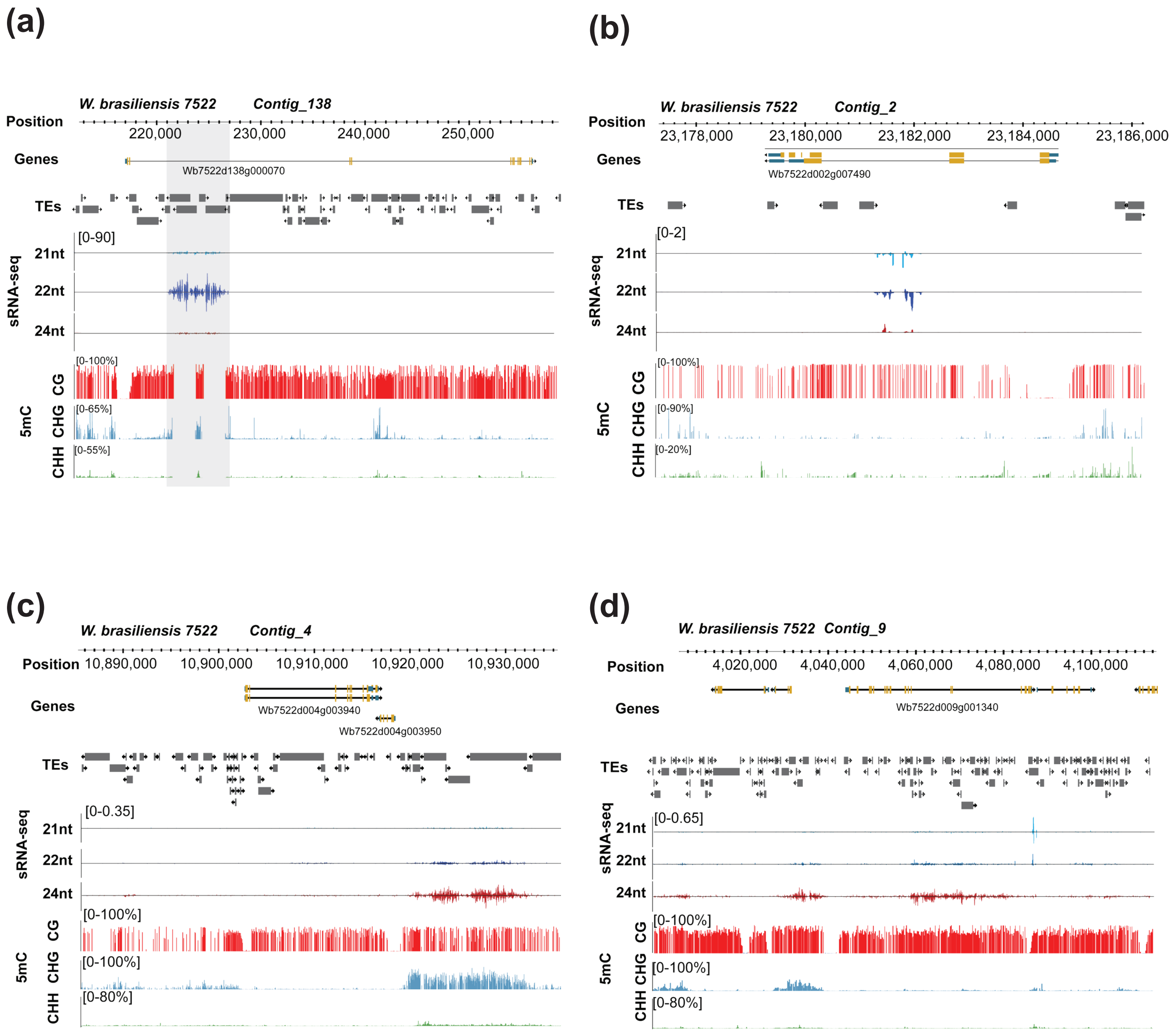


**Figure S29. Examples of siRNA-producing Intergenic and Intronic regions in Wolffia. (a-b)** Genome browser screenshots of intronic IR+ clusters producing sRNAs. Genomic coordinates, Gene and TE annotations, siRNA seq (21,22 and 24nt) coverage and DNA methylation levels for the three 5mC contexts (CG, CHH and CHH) are presented as tracks. Coverage values are expressed in reads per million (RPM). **(c-d)** Genome browser snapshots of c) intergenic and d) intronic siRNA-producing TEs. Genomic coordinates, Gene and TE annotations, siRNA seq (21,22 and 24nt) coverage and DNA methylation levels for the three 5mC contexts (CG, CHH and CHH) are presented as tracks. Coverage values are expressed in reads per million (RPM).

### SUPPLEMENTAL FIGURE S30

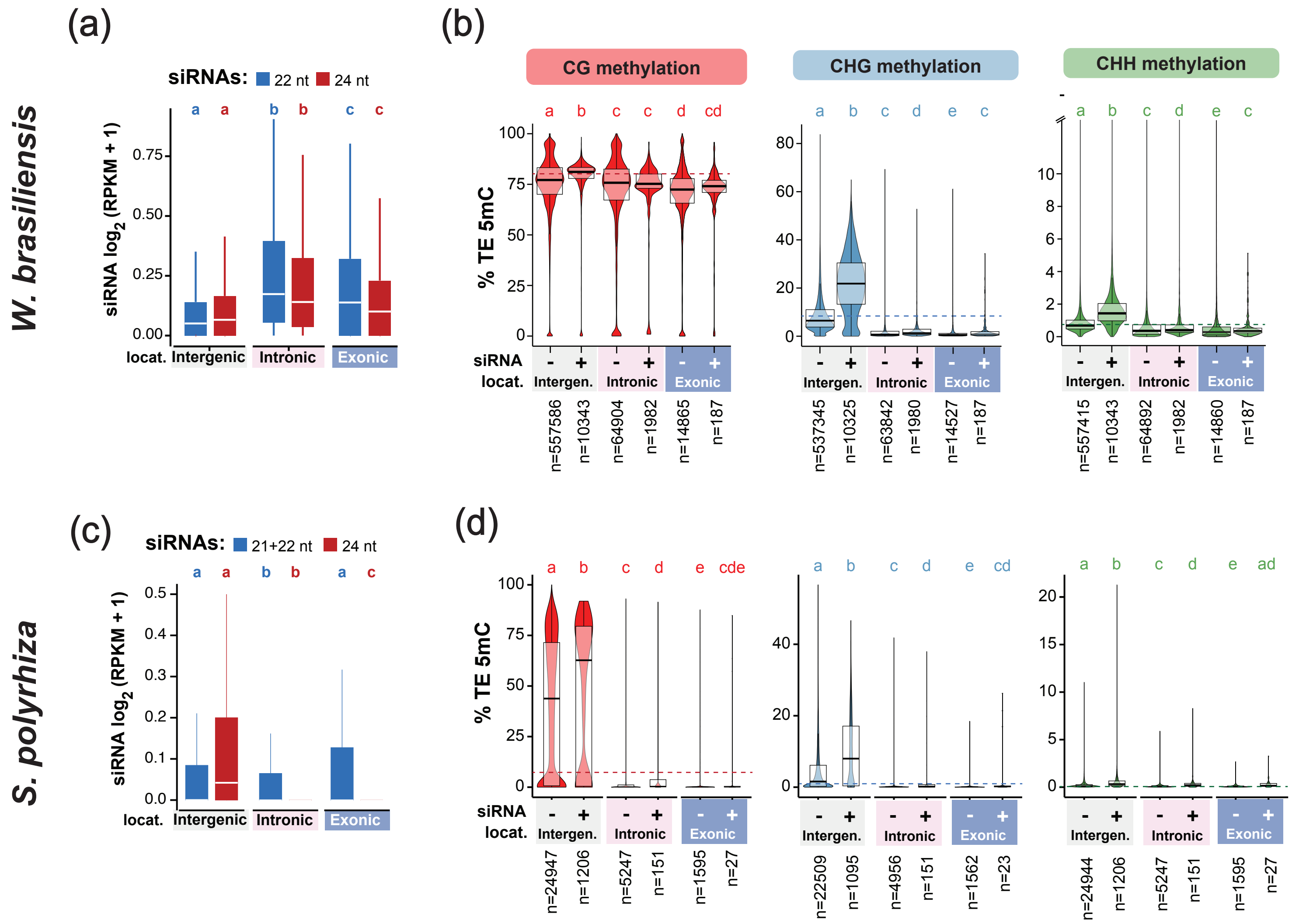


**Figure S30. TE siRNAs and DNA methylation by genomic compartment in Wolffia and Spirodela.** **(a)** siRNA levels (RPKM+1) levels of intergenic, intronic and exonic TEs in Wolffia. **(b)** 5mC levels in all three contexts for siRNA- and non-siRNA-producing TEs in intragenic, intronic and exonic locations in Wolffia. Dashed line indicates average clobal methylation levels in each context. **(c-d)** Same as (a-b) but in Spirodela. In (b-d) different letters indicate significant differences between groups (Kruskal-Wallis followed by Dunn's test $p<0.05$). All P-values can be found in Tables S7.

### SUPPLEMENTAL FIGURE S31

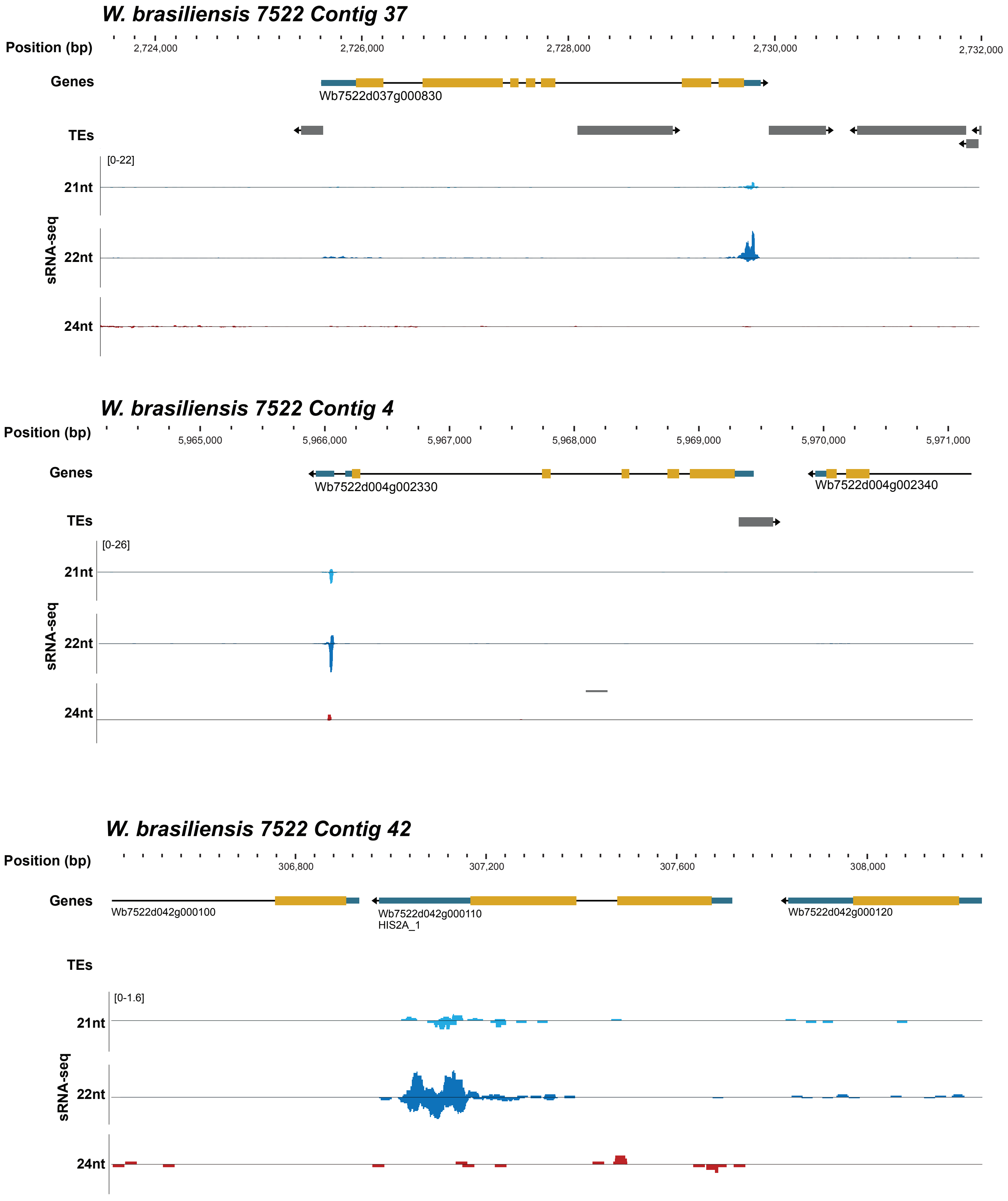


**Figure S31. Examples of siRNA producing 3'UTRs in Wolffia.** Genome browser screenshots from three 3'UTR siRNA producing example sequences. Genomic coordinates, Gene and TE annotations and siRNA seq (21,22 and 24nt) normalized coverage (RPM) are presented as tracks. Bottom panel depicts siRNAs mapping to the 3'UTR of a histone H2A gene (Wb7522d042g000110).

### SUPPLEMENTAL FIGURE S32

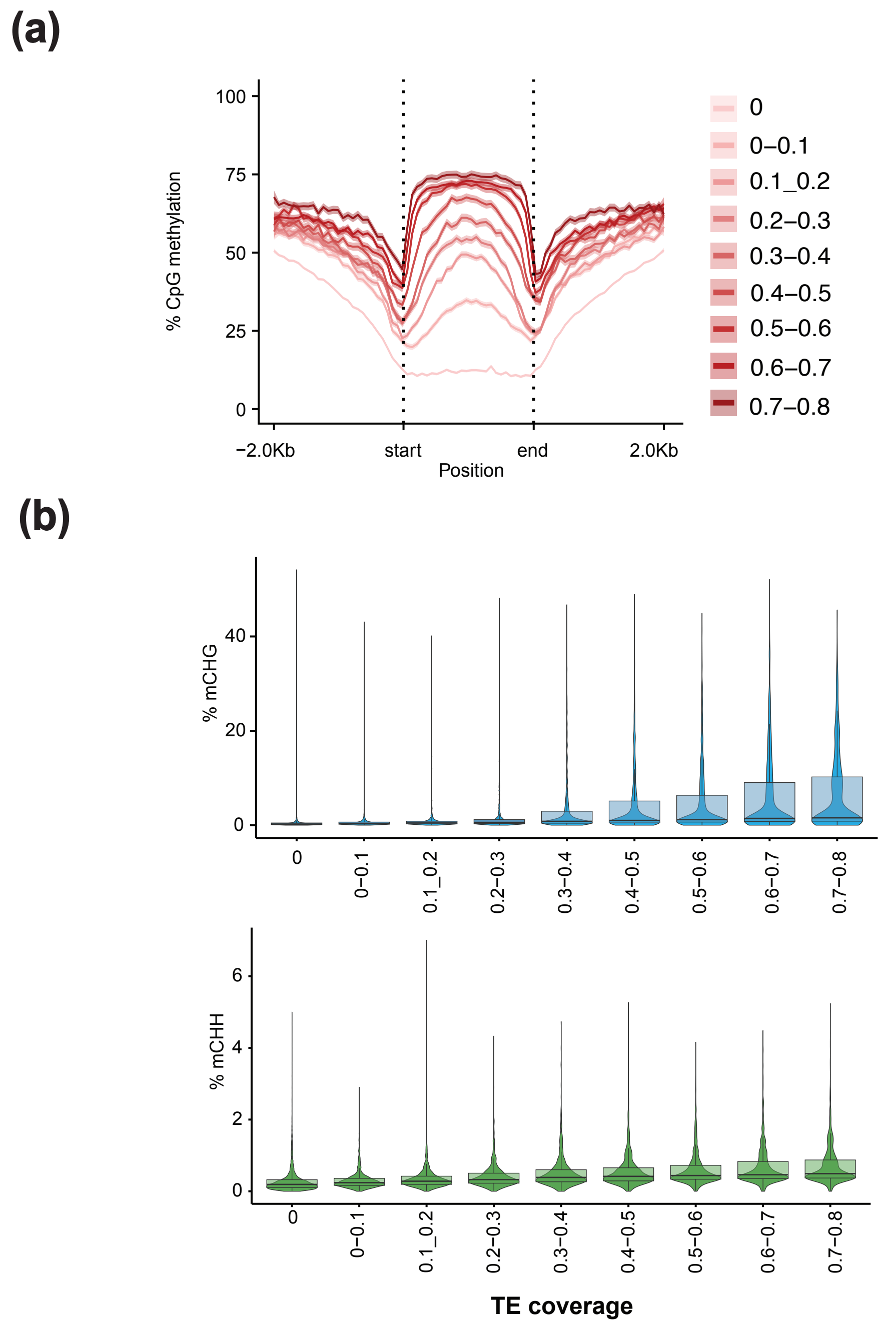


**Figure S32. DNA methylation levels on gene annotations correlate with gene TE coverage in Wolffia. (a)** Metaplots showing the levels of CG methylation over gene annotations categorized according to their relative TE content [0-100%]. Genes with a TE coverage higher than 80% were considered as TE-genes and excluded from this analysis. **(b)** Stand-alone violin and boxplots from Figure 8g for CHG and CHH methylation contexts. Boxplots represent the median as a solid bar, with box upper and bottom limits representing the first and third quartiles. Whiskers range is 1.5 times the interquartile range. Statistically significant differences between groups are shown in Fig 8g.

### SUPPLEMENTAL FIGURE S33

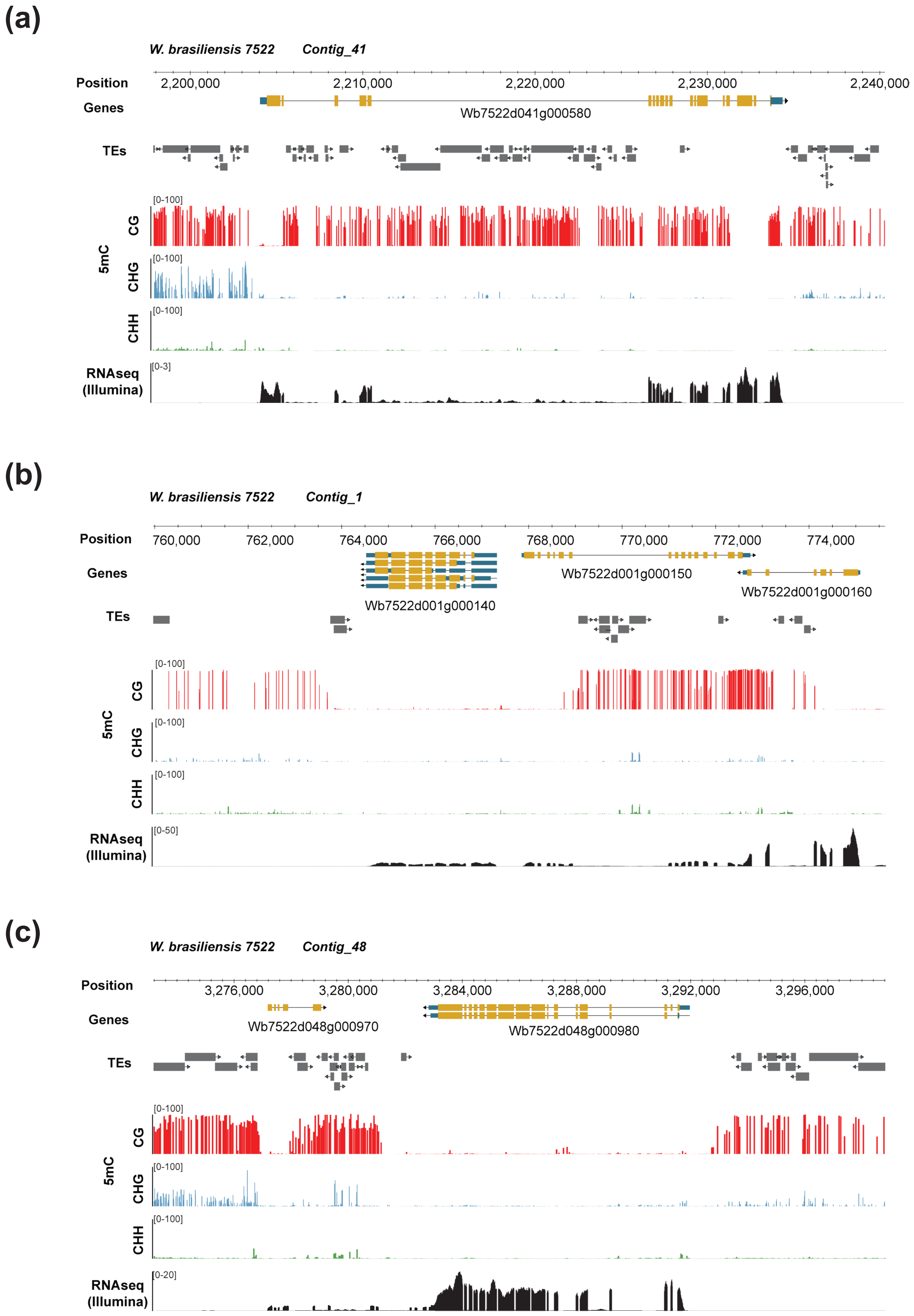


**Figure S33. Examples of gene body methylation on genes containing or not TEs in Wolffia.** Genome browser snapshots of genes with or without TEs and their respective gene-body methylation. **(a)** Example of a gene overlapping with TEs (coverage>0). **(b)** Genomic region with three genes, showing distinct TE coverage and its correlation with DNA methylation. **(c)** Example of a gene without any TE. Genomic coordinates, Gene and TE annotations and DNA methylation levels [0-100%] for the three 5mC contexts (CG, CHH and CHH) are presented as tracks.

### SUPPLEMENTAL FIGURE S34

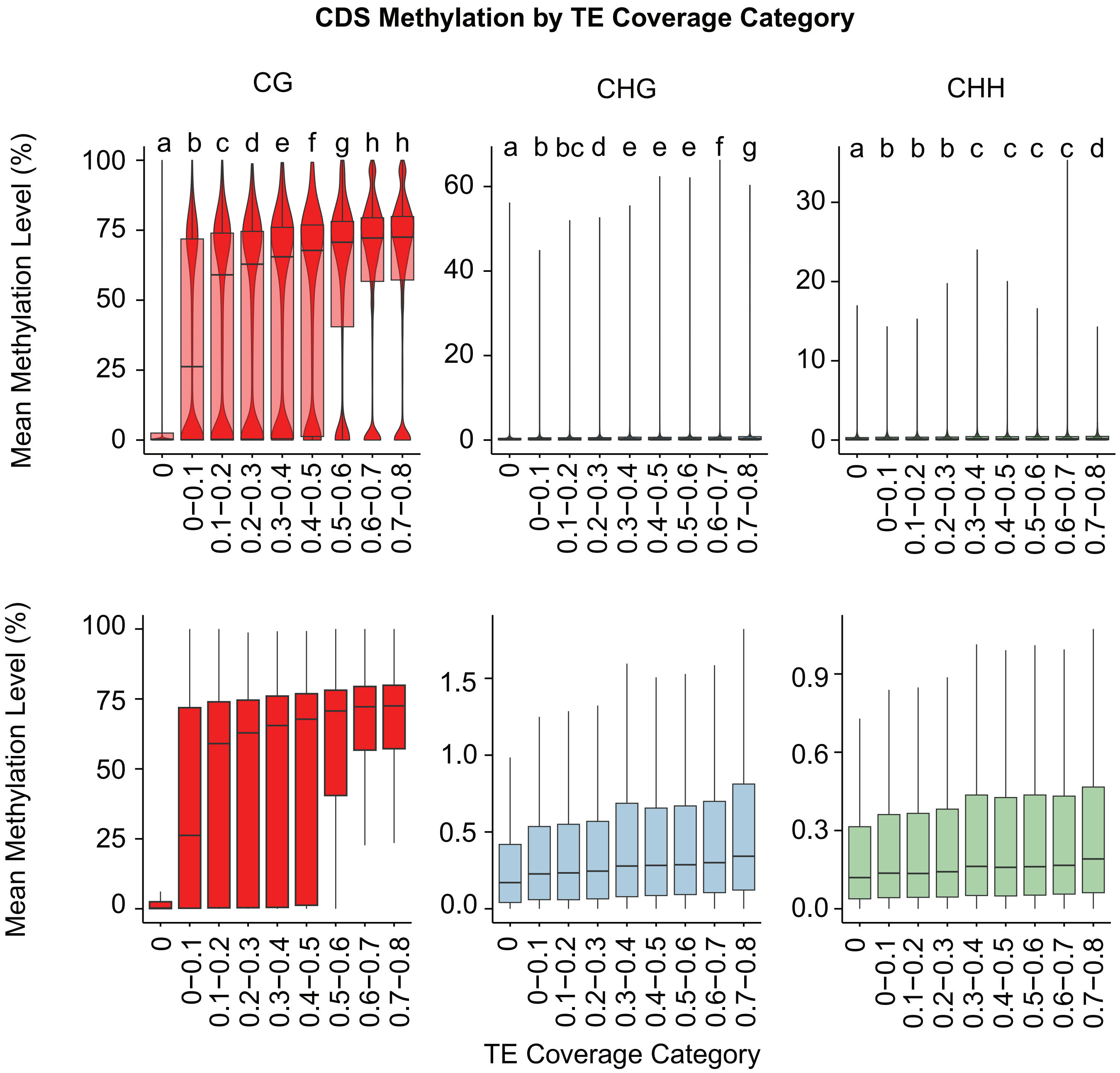


**Figure S34. DNA Methylation analysis on CDS sequences in *W. brasiliensis*.** Top: Violin and boxplots showing the DNA methylation level distribution on the three 5mC methylation contexts for CDS genic regions, according to their respective gene TE coverage [0-80%]. Genes with a TE coverage higher than 80% were considered as TE-genes and their CDS sequences excluded from this analysis. Bottom: Boxplots (from top plots) without the violin distribution plots, depicting the levels of DNA methylation for better visualization. In all boxplots the median is represented as a solid bar, with box upper and bottom limits representing the first and third quartiles. Whiskers range is 1.5 times the interquartile range. Differences and statistical significance indicated by letters was calculated using a Kruskal-Wallis test followed by a *post-hoc* Dunn's test. All test statistical P-values can be found in Table S9.

### SUPPLEMENTAL FIGURE S35

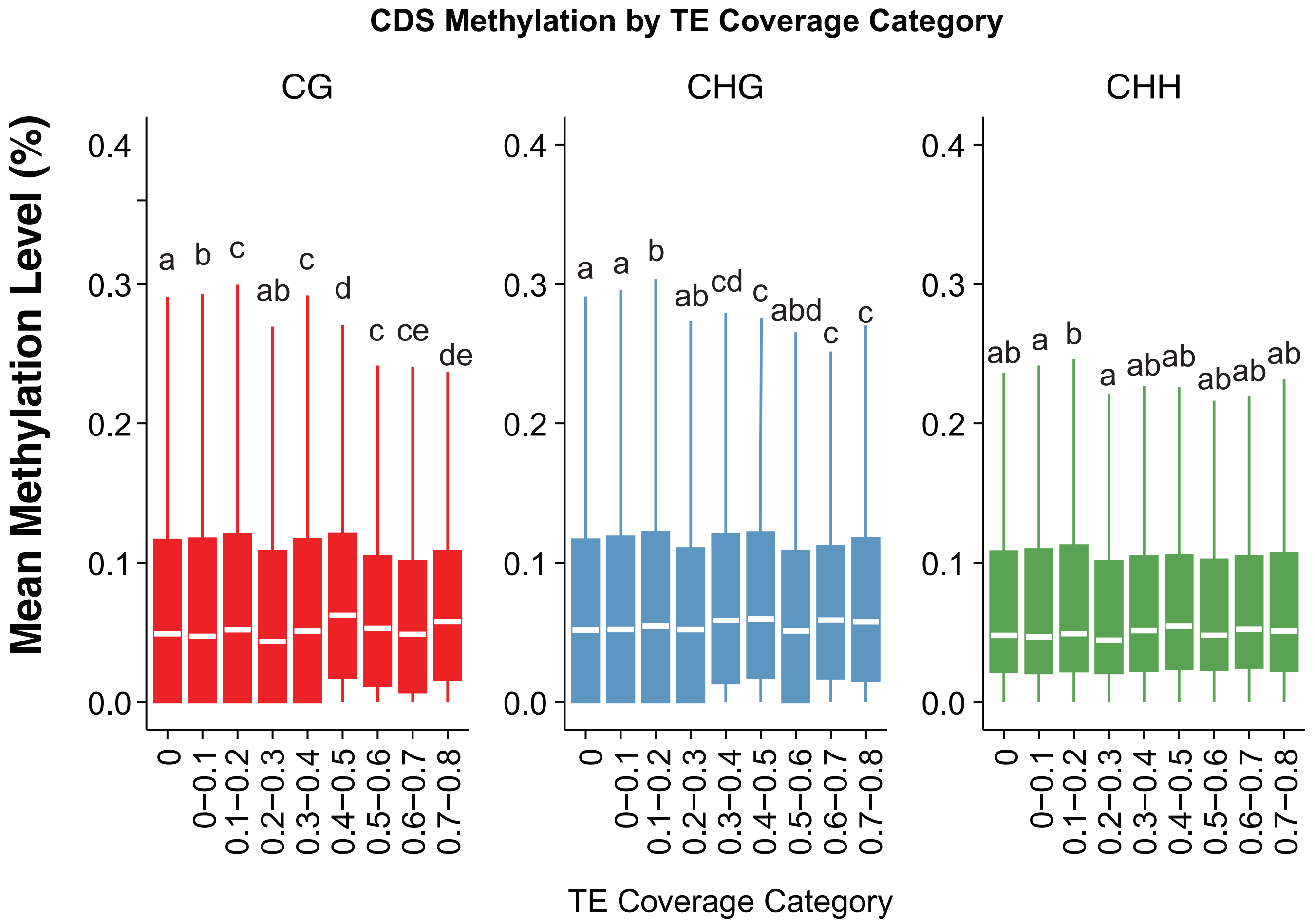


**Figure S35. DNA Methylation analysis on CDS sequences in *S. polyrhiza*.** Boxplots showing the DNA methylation level distribution on the three 5mC methylation contexts for CDS genic regions, according to their respective gene TE coverage [0-80%]. Genes with a TE coverage higher than 80% were considered as TE-genes and their CDS sequences excluded from this analysis. In all boxplots the median is represented as a solid bar, with box upper and bottom limits representing the first and third quartiles. Whiskers range is 1.5 times the interquartile range. Differences and statistical significance indicated by letters was calculated using a Kruskal-Wallis test followed by a *post-hoc* Dunn's test. All test statistical P-values can be found in Table S9.

#### EXTENDED MATERIALS AND METHODS

##### Plant material

All duckweed accessions from *Spirodela polyrhiza* (#5676), *Landoltia punctata* (#7487), *Lemna minor* (#8623), *Wolffiella gladiata* (#8768), *Wolffia brasiliensis* (#7500, #7522, #7925, #8743, #9134, #9178, #9273, #9380, #9386, #9390, #9403, #9447, #9597 and #9656), and *Wolffia arrhiza* (#7158, #7193, #7196, #7215, #7246, #7265, #7347, #7421, #7678a, #7699, #7736, #8272, #8450,#8618, #8649, #8953, #9113, #9412, #9492 and #9615) were obtained from the Landolt collection available at the CNR-IBBA-MIDW collection (Milan, Italy; https://biomemory.cnr.it/collections/CNR-IBBA-MIDW).

Axenic cultures from each duckweed accession were obtained following the protocol described in (Dombey *et al.*, 2025) with species-specific modifications. Briefly, Spirodela, Landoltia and Lemna accessions were sterilized in 10 mL of 1% DanKlorix® solution (Colgate-Palmolive) with mild agitation for 1 minute, washed 3 times with sterile water and transferred to 0.5X Schenk and Hildebrandt (SH, Duchefa Biochemie, #S0225) media. Wolffiella and Wolffia accessions were both cultured in N media (Appenroth *et al.*, 1996) after sterilization in 1% DanKlorix® for 1 and 2 minutes respectively and subsequent washing.

Axenic cultures were long-term kept on solid (0.5X SH or N, 0.4% gelrite) media in MagentaTM GA-7 containers, in a plant growth cabinet (SEQI-P4, Percival) at 15°C with 16h light /8h dark light cycle, low light intensity (19.5 µmol $m^{-2}$ $s^{-1}$).
To obtain plant material, inoculums from the long-term storage cultures were grown on 0.5X SH or N liquid media at 21°C with a light intensity of 85 µmol $m^{-2}$ $s^{-1}$ in MagentaTM GA-7 containers sealed with Leucopore tape (Duchefa Biochemie #L3301), under long-day conditions (16h light /8h dark). To prevent confluency, all accessions were sub-cultured every 1.5~2 weeks.

##### Plant imaging:

Pictures of fronds were taken using a Keyence VHX-7000 (Keyence, Osaka, Japan) and an Andonstar AD409 digital microscope. Fluorescence images of single *W. brasiliensis* transformed fronds were obtained using a stereo zoom microscope Zeiss Axio Zoom V16 (chlorophyll autofluorescence: excitation 655 nm, detection 667 nm; eGFP: excitation 488 nm, detection 509 nm) or a Sapphire FL Biomolecular scanner

(Azure biosystems) (chlorophyll autofluorescence: excitation 658 nm, detection 710 nm; eGFP: excitation 488 nm, detection 518 nm).

**Tubulin-based polymorphism (TBP) analysis**

Genotyping of the analyzed accessions was performed by comparison of the amplification profiles of intron 1 of the ß- tubulin genes, according to a standard TBP amplification protocol, from 30 ng of total genomic DNA. Polymorphisms in amplicon length were resolved by capillary electrophoresis. DNA samples were independently analyzed twice. Data analysis was performed as in (Braglia *et al.*, 2023)

**Chromosome spreads**

Healthy fronds were incubated in 2mM 8-hydroxyquinoline (Merck, #252565) at 37°C for 2h, and fixed in Carnoy's solution (6:3:1 Ethanol:Chloroform:Glacial Acetic Acid) for 24h, and then transferred into a 3:1 Ethanol:Glacial Acetic Acid solution for at least 48h.

Chromosome spreads were then performed according to the protocols described in (Cao *et al.*, 2015; Hoang *et al.*, 2019) with the following modifications. Briefly, fronds were rinsed three times for 5 minutes each in dH2O, followed by another three 5 minutes washes in sodium-citrate buffer (10 mM, pH 4.6). Enzymatic maceration was performed by incubating samples in 2 mL of a 0.4% enzymatic mixture containing cellulase (Sigma-Aldrich, #C1184), pectinase and pectolyase (Sigma, #P3026) in sodium citrate buffer for 30 minutes at 37°C. Samples were then washed twice for 5 minutes in sodium citrate buffer and twice again for 5 min in dH2O.

Samples were placed in the center of a microscope slide with a drop of 60% acetic acid. Two coverslips (24 × 24 mm) were positioned—one to the side and one on top of the sample—and the tissue was mechanically disrupted by repeated tapping with a thin instrument. The side coverslip was removed, and the top coverslip was pressed firmly with thumb pressure to squash the sample. Preparations were then frozen in liquid nitrogen, the coverslip was removed, and the slide was briefly rinsed in 100% ethanol and air-dried.

The slides were then submerged in a pepsin enzymatic solution (50 µg/mL 0.01 N HCl; Sigma-Aldrich, #P7000) for 3 min at 37 °C, washed twice for 5 minutes in 2x SSC buffer and fixed in 4% formaldehyde in 2x SSC for 10 minutes. Preparations were then rinsed twice for 5 min in 2x SSC, and dehydrated through an ethanol series (70%,

96%, 100%) for 2 min each, air-dried, and mounted with 10 µL of DAPI-containing Vectashield Antifade Mounting Medium (Vector Laboratories, #H-1200). Images of the DAPI-stained chromosomes were taken using a Zeiss Axio Imager.Z2 with a sCMOS camera.

**Genome sequencing and gene annotation**

*Wolffia brasiliensis* nuclei were isolated according to the Protocol C described in (Lutz *et al.*, 2011) with the following modifications. 10 g of tissue were freeze-ground to fine powder with mortar and pestle. The powder was then resuspended in 200 mL MEB buffer (MES 10 mM; MgCl2 10 mM; PVP 2; sodium metabisulfate 10 mM; Hexylene Glycol 1M; 2-mercaptoethanol 5 mM; sodium diethyldithiocarbamate 0.5%; EGTA 6 mM; L-lysine-HCl 200 mM, pH 5), filtered through 2 layers of Miracloth (Millipore, #475855) and incubated for 15 minutes on ice after the addition of Triton-X-100 to a concentration of 0.5%. Nuclei were then pelleted (800 g for 20 minutes at 4°C) and washed with MPDB buffer (MES 10 mM; MgCl2 10 mM; sodium metabisulfate 10 mM; Hexylene Glycol 500 mM; 2-mercaptoethanol 5 mM; Triton-X-100 0.5%) until the pellet was white and supernatant clear. The nuclei pellet was then purified through a 37.5% Percoll® (Merck, #P1644) cushion.

HMW DNA extraction from purified nuclei was performed following the protocol described in (Rabanal *et al.*, 2022). In summary, the pellet was resuspended in G2 lysis buffer (Qiagen, #19060) with 50 µg RNAse A and incubated for 30 minutes at 37°C. Proteinase K (ThermoFisherscientific, #EO0491) was added to a final concentration of 200µg/mL and incubated 3h at 50°C, and then centrifuged (10500g for 15 minutes at 4°C). HMW DNA was then purified using a Qiagen genomic top 100G (Qiagen, #10243) previously equilibrated according to manufacturer's instructions. DNA was then precipitated with 0.7 volumes of isopropanol and fished with a glass rod and transferred to 300 mL TE buffer (10 mM Tris-HCl pH 8.0 and 1 mM EDTA). Library preparation was carried out using SMRTBell prep kit v2 (Pacific Biosciences, PN 100-938-900) following manufacturer's instructions. Sequencing was performed on a PacBio® Sequel® II.

HiFi reads were obtained using PacBio ccs package (v 6.4.0). Reads were assembled with HiFiasm (v 0.16.1; -D 10) and those belonging to contigs with >80% length and >95% identity with chloroplast sequence were filtered out before a new assembly was performed using the same parameters. Contig redundancy was assessed using BLAST+ (v 2.8.1 default parameters) and those with high overlap (>95% identity) to a bigger contig were removed.

Gene annotation was performed with Funannotate (v1.8.13). First, a round of Training (Funannotate train --pacbio_isoseq --max_intronlen 40000) and gene prediction (Funannotate predict -w pasa:10 CodingQuarry:0 --busco_seed_species arabidopsis --busco_db embryophyta --transcript_evidence --repeats2evm --optimize_augustus --genemark_mode ES) using PacBio Isoseq reads was performed. A final update with Illumina RNAseq short reads was performed (Funannotate update -l -r), and protein domain identification with Interproscan (v5.64-96.0; -appl CDD,PANTHER,Pfam -goterms) before the final annotation with Funannotate default parameters.

**Small RNA seq**

- **Extraction**

Purification of small RNAs from TraPR (Argonaute-loaded) total RNA was performed according to the protocol described in (Grentzinger *et al.*, 2020; Dombey *et al.*, 2025). In brief, 100mg of flash-frozen plant tissue was mechanically disrupted with 1mm ø glass beads (Carl Roth, #A554.1) using a Silamat S6 (Ivoclar Vivadent). The resulting powder was homogenized in TraPR lysis buffer and clarified with a brief centrifugation (10000g, 5 minutes, 4°C) before being applied to TraPR columns (Lexogen, #128.24). Small RNAs were extracted from the TraPR eluate fractions using 1 volume of acidic Phenol:Chloroform:Isoamyl alcohol (PCI 50:49:1, pH 4.5-5) (Carl Roth, #X985.1). After phase separation by centrifugation (12000g for 10 min at 4°C), RNA was precipitated from the aqueous phase with isopropanol and sodium acetate (3M pH 5.2, 10% v/v) overnight at -20°C. Pellets were recovered (12000g for 30 min at 4°C), rinsed with 75% ethanol, and resuspended in 10–20 µL of nuclease-free water.

For total small RNA libraries, an aliquot of the TraPR lysate was set aside before loading onto the columns and processed by the same phenol-chloroform procedure described for the TraPR eluates. The resulting RNA was loaded and separated in a 15% denaturing polyacrylamide-urea gel. A microRNA marker (20 µL; New England Biolabs, #N2102S) was included as a size reference. The 17-25-nt fraction was excised and purified using the ZR small-RNA PAGE Recovery Kit (Zymo Research, #R1070).

- **Small RNA library preparation**

Small RNA libraries were generated as previously described (Hayashi *et al.*, 2016; Jayaprakash *et al.*, 2011). In summary, small RNAs were first ligated to 3′ barcoded DNA adapters using truncated T4 RNA ligase 2 (New England Biolabs, #M0373). Ligation products were resolved in a 12% denaturing polyacrylamide-urea gel. The appropriate size range was excised and recovered using ZR small-RNA PAGE Recovery Kit (Zymo Research, # R1070). The purified products were subsequently ligated to 5′ barcoded RNA adapters using T4 RNA ligase 1 (New England Biolabs, #M0204). Both 5' and 3' adapters carried 4 randomized nucleotides at their termini to mitigate ligation bias. Adapter-ligated small RNAs were reverse-transcribed and PCR-amplified using Illumina-compatible primers. Library size profiles were evaluated on a 5200 Fragment Analyzer System (Agilent, #M5310AA).

- **Small RNA sequencing and bioinformatic analysis**

For small RNA profiling of multiple accessions, sequencing of two biological replicates per accession was performed on an Illumina Novaseq X to generate 150-bp paired-end reads. Raw reads were adapter-trimmed and quality-filtered with Trimmomatic (Bolger *et al.*, 2014) (v0.39). Trimmed reads from both pair-end were merged using Pandaseq v2.11 (Masella *et al.*, 2012). A custom-made script was used to categorize and summarize the reads in different sizes for visualization.

For *Wolffia brasiliensis* hpScarlet transformation experiments, two independent biological replicates were sequenced on the same instrument and processed as mentioned above. Subsequent analysis was performed as described in (Dombey *et al.*, 2025). Processed reads were aligned specifically to the hpScarlet hairpin sequence using Bowtie (Langmead and Salzberg, 2012) (v1.2.2; -e 50 -a -v 0 --best --strata --nomaqround -y --phred33-quals --no-unal --sam). Mapped reads were then normalized using bamCoverage (Ramírez *et al.*, 2016) (Deeptools v3.3.1 --normalizeUsing CPM --binSize 1).

For *Wolffia brasiliensis* genome-wide small RNA profiling and analysis, 3 replicates were sequenced on an Illumina NextSeq 550 to generate 75-bp single-end reads. Raw reads were adapter-trimmed and quality-filtered with Trimmomatic. Processed reads were merged and aligned to the *Wolffia brasiliensis* generated genome using Bowtie (-v 1). Small RNA-producing loci were identified and annotated using ShortStack

((Johnson *et al.*, 2016) v 3.8.5 --foldsize 1000 --mincov 0.6rpm). Mapped reads were then normalized using bamCoverage as above mentioned.

- **Small RNA Categories definition**

The empirical probability density function of the filtered ratio (values was estimated using kernel density estimation (KDE). The "nrd0" bandwidth selection method was chosen for its robustness to sparse or discrete data.

Local minima (valleys) in the estimated density curve were identified by detecting points where the density function slope changed from negative to positive. Among the identified valleys, the one closest to 1 (representing a neutral 22-nt:24-nt ratio) was selected as the primary separation point. Based on the identified valley two symmetric thresholds were established: Left threshold (LT), representing the primary separation point, and Right Threshold as 1/LT. Each locus was then classified into one of the three intervals defined (x<LT, LT<x<RT, x>RT).

**Flow cytometry**

Genome size estimation for different *Wolffia* accessions was performed by flow cytometry following the protocol described in (Temsch *et al.*, 2010; Barragán-Borrero *et al.*, 2026). *Solanum pseudocapsicum* (1C = 1.295 pg) and *Pisum sativum* 'Kleine Rheinländerin' (1C= 4.42 pg) (Greilhuber and Ebert, 1994) served as the internal standards. Fresh fronds (25 mg) of *Wolffia* samples were mechanically disrupted by razor blade sectioning (Astra superior platinum; Gillette) in a 94 mm petri dish with 100 µL of nuclei isolation buffer (NIB: 0.1 mM citric acid, 0.5% v/v Triton X-100, pH 1.5). The homogenate was diluted to a final volume of 550 µL with additional NIB and filtered through a 35 µm nylon cell strainer (Corning, #352235). RNase A (ThermofisherScientific, #EN0531) was added to a final concentration of 0.15 mg/mL and the preparation was incubated at 37°C for 30 min. Nuclei were labeled with propidium iodide (6 mg/mL, pH 9.5; Merck, #P4170-25MG) and held at 4°C in the dark for 16 h. Flow cytometry was performed on a FACSCanto (BD Biosciences) equipped with a blue laser (488 nm, 20 mW) and a red laser (633 nm, 17 mW). Propidium iodide stained-nuclei were detected with a bandpass filter of 585/42. Data acquisition was conducted using the BD FACS DiVa software. Analysis of the flow cytometry data was performed using FlowJo (BD Biosciences, v10.6.1). Genome sizes were determined by comparing the mean fluorescence intensity of the G1 peak of each sample to that of the standard. Three independent biological replicates were analyzed for each accession, and for each a total of 10000 nuclei were counted.

**Phylogenetic analysis and protein domain annotation:**

Putative *Wolffia brasiliensis* orthologues / paralogues of *Arabidopsis thaliana* and *Spirodela polyrhiza* silencing components were identified using homology-based searches. Query nucleotide and amino-acid sequences were retrieved from TAIR10 and the Spirodela dataset deposited at [https://doi.org/10.5281/zenodo.14825003] (Dombey *et al.*, 2025) and used as inputs against the *Wolffia brasiliensis* proteome with BLAST+(Camacho *et al.*, 2009); (v2.8.1, blastp, -qcov_hsp_perc 60). These protein queries were also mapped to the *W. brasiliensis* genome assembly using tblastn (default settings) to capture loci not represented or incompletely represented in the predicted protein set. Hits were retained as candidates when they contained the expected conserved domains, final gene sets were organized into orthogroups by comparison with curated angiosperm proteins from each orthogroup.

To validate candidates, homologous protein sequences from other angiosperms were collected from UniProt (EMBL-EBI) (https://www.uniprot.org) (Consortium *et al.*, 2025) and used as BLAST queries. Multiple sequence alignments were generated in CLC Main Workbench v20 (QIAGEN) using a gap opening penalty of 10 and a gap extension penalty of 1. Phylogenetic relationships were first summarized with unrooted Neighbor-Joining trees computed under a Jukes–Cantor distance model, and maximum-likelihood trees were additionally inferred with 1,000 bootstrap replicates. Pairwise protein identity analysis of DCL proteins on *Spirodela polyrhiza* and *Wolffia brasiliensis* was performed using CLC Main Workbench. Domain architecture was annotated using Pfam-A v35 (Mistry *et al.*, 2020) within CLC Main Workbench and cross-checked with independent InterPro-based predictions to confirm and extend domain calls.

**TE annotation**

Transposable elements in the *Spirodela polyrhiza* genome were annotated using a combined *de novo* approach that integrated EDTA (Ou *et al.*, 2019); (v2.1.3, --species others --sensitive 1) and RepeatModeler2 (Flynn *et al.*, 2020) (v 2.0.1, -engine ncbi -

LTRStruct). To ensure accuracy and sensitivity, both tools were run in triplicate and the resulting annotation libraries were aligned and merged using CDHit (v4.8.1). The classification of TEs was further refined using MCHelper (Orozco-Arias *et al.*, 2024). TEs were then annotated with RepeatMasker ((Smit *et al.*, n.d.) v4.1.0; -xsmall -a -e ncbi -q -no_is -norna -nolow -div 40 -cutoff 225). Using TEsorter (Zhang *et al.*, 2022) (version 1.3, -db rexdb-plant -st nucl), the protein domains and individual TE copies classification were determined.

**Transcriptome analysis**

Approximately 100 mg of plant material from *W. brasiliensis* was flash-frozen and ground to a fine powder using 1mm ø glass beads (Carl Roth, #A554.1) on a Silamat S6 (Ivoclar Vivadent). RNA was then extracted with the Rneasy Plant Mini Kit (Qiagen, #74904) following manufacturer's instructions.

Illumina short read libraries were prepared using the NEBNext® Ultra™ II RNA Library Prep Kit for Illumina® (NEB, #E7775), and sequenced on a NovaSeq S4 to generate 150bp paired-end reads. Short-read data were adapter/quality trimmed and assessed with fastp v0.20.1 (Chen *et al.*, 2018) after which reads were quantified against the *W.brasiliensis* genome to estimate transcript abundance. Gene and transposable element expression were quantified separately using Kallisto (Bray *et al.*, 2016), and transcript abundance values (TPM) were averaged across three biological replicates.

For *Arabidopsis thaliana* seedlings, and *Spirodela polyrhiza* fronds publicly available datasets were used (GSM6892967, GSM6892968, GSM6892969, SRX26200034, SRX26200035, SRX26200036) (Li *et al.*, 2023; Dombey *et al.*, 2025).

The Iso-Seq library was prepared according to manufacturer's instructions using the SMRTBell prep kit v2 (Pacific Biosciences, PN 100-938-900) and sequenced on PacBio® Sequel® II. Reads were processed with the nf-core/isoseq pipeline (v1.0.0; nf-core/isoseq), and high-quality transcripts were aligned to the *W.brasiliensis* genome using minimap2 v2.17 using default parameters (Li, 2021).

**Agrobacterium-mediated transient transformation**

Transformation of *W.brasiliensis* fronds was performed according to the protocol described in (Dombey *et al.*, 2025; Barragán-Borrero *et al.*, 2026) with minor

modifications. In brief, overnight *Agrobacterium tumefaciens* strain EHA105 bacterial cultures were pelleted and resuspended to $OD_{600}$ of 0.6 in Agroinfiltration Medium (10 mM $MgCl_2$, 5% w/v sucrose, 200 µM acetosyringone (Sigma-Aldrich, #D134406), pH 5.6). Before infiltration, Silwet L-77 (Kurt-Obermeier GmbH, #7060-10) was added to a final 0.02% (v/v) concentration. Approximately 1 g of *W. brasiliensis* fronds from axenic cultures was incubated with 1 mm ø glass beads (Carl Roth, #A554.1) in agitation (200 rpm, 15 minutes, 21ºC), and then transferred to ice-cold 20 mg/L L-glutamine solution (PanReac AppliChem, #A1420) for 20 min (Yang *et al.*, 2018). Following the treatment, fronds were submerged in 10 mL of the *Agrobacterium* suspension inside a Gosselin screw-cap container (Corning, #TPC30C-002) placed in a vacuum desiccator, and vacuum infiltrated at −80 kPa twice for 30 min.
Following infiltration, the *Agrobacterium* solution was aspirated and replaced with 5 mL of N-medium supplemented with 100 µM acetosyringone, 1% sorbitol, 5% sucrose. Plants in N-media were transferred to culture dishes (100 × 20 mm; Greiner Bio-One, #664160) containing 2 layers of sterile filter paper, sealed with parafilm and cultivated in darkness for 4 days. Fronds were then rinsed 2-3 times in antibiotic-free N-media before being transferred to Magentas™ and cultivated in the same conditions described under “Plant material”.

For the image-based analysis, plants were scanned using a Sapphire Imaging system (Azure Biosystems) and fluorescence quantification was performed in Fiji software(Schindelin *et al.*, 2012). Chlorophyll autofluorescence was used to identify plants that were viable and generate segmented masks. Segmentation was performed using a gaussian blur filter, followed by binary thresholding, watershed separation and particle analysis (default parameters). GFP fluorescence was then quantified over the resulting segmented areas. At each time point, plants were classified as GFP-positive when their fluorescence intensity exceeded the mean GFP signal of control samples by more than two standard deviations.

**DNA methylation**

For the assessment of DNA methylation, EM-seq was performed as in (Dombey *et al.*, 2025) with minor modifications. In summary, DNA extraction was performed according to the instructions provided in the DNeasy® Plant mini kit (Qiagen, #69104). Libraries were prepared with the NEBNext® Enzymatic Methyl-seq Kit (NEB, #M7634) according to the manufacturer's protocol, and sequenced on an Illumina NovaSeq S4 to produce 150-bp paired-end reads. Reads were then processed with fastp to remove adaptors and low quality reads, and aligned with Bismark (Krueger and Andrews, 2011) (v0.22.2; --non_directional *-q* --score-min *L,0,–0.4*) to the *Wolffia brasiliensis* genome. samtools v1.9 was used for alignment handling. Alignments were then deduplicated and used to compute per-cytosine weighted methylation levels (deduplicate_bismark, bismark_methylation_extractor). Conversion efficiency was monitored using *W. brasiliensis* chloroplast sequences retrieved from NCBI (MN850406.1) revealing a >98.6% conversion rate in all replicates. Replicates were then merged, and cytosines with a coverage <4 were filtered out for downstream analyses. Global 5mC levels, and feature-specific 5mC levels were calculated with DeepTools (multiBigwigSummary, computeMatrix and plotProfile) using default parameters unless specified in the figure legends. When calculating DNA methylation on features, to ensure robustness in the analysis, only those covered with a minimum of 4 cytosines were considered.

**Gene and TE overlap analysis**

Classification of TEs based on the overlap with genes was performed using ParasiTE (Berthelier *et al.*, 2023), using the Iso-Seq reads as input. In brief, fully exonic TEs were those with a >80% overlap with exons, as Intronic if <1% of the TE length overlapped with an exon and partially exonic those TEs within 1%-80% overlap with exonic sequences. Intragenic TEs that were not overlapping with a predicted gene model were filtered out as Intergenic.

**Visualization and Statistical analysis:**

Data visualization plots were prepared as described in the relevant sections. All statistical analyses were conducted in R (v4.2.2; R Project). The statistical tests applied are specified in the corresponding figure legends. Figures were generated in R using circlize (Gu *et al.*, 2014) ;v0.4.15) and ggplot2 (Wickham, 2016); v3.4.4)

**Transposon divergence**

To study the divergence of TEs, Kimura substitution scores, that account for the degree of sequence divergence of individual TE sequences from the consensus, were calculated from the output of RepeatMasker using the calcDivergenceFromAlign.pl script (Smit *et al.*, n.d.).

##### Annotation of Inverted Repeats

Inverted repeats were identified using the EMBOSS einverted (Rice *et al.*, 2000) – (maxrepeat 20000 -gap 12 -threshold 50 -match 3 -mismatch -4) on small RNA clusters identified with Shortstack (--foldsize 1000, --pad 150, --mincov 2rpm). Self-dotplots for Inverted repeat visualization were generated using Gepard (Krumsiek *et al.*, 2007) v2.1 with a 100bp window size.

##### Plasmids used in this study

The pZmUbq:GFP plasmid (EPR Plasmid #869) was previously generated in the lab (Barragán-Borrero *et al.*, 2026). pGGSun-pUBIPars-hpScarlet was a gift from Marco Incarbone (Max-Planck-Institute of Plant Physiology).

##### Jbrowser

Genome screenshots were obtained from Jbrowse2 (Diesh *et al.*, 2023) https://jbrowse.org/jb2/download/
